# Temporal and spatial segregations between phenotypes of the Diablotin Black-capped Petrel *Pterodroma hasitata* during the breeding and non-breeding periods

**DOI:** 10.1101/2022.06.02.491532

**Authors:** Yvan G. Satgé, Bradford S. Keitt, Chris P. Gaskin, J. Brian Patteson, Patrick G.R. Jodice

## Abstract

**Aim:** Despite growing support for ecosystem-based approaches, conservation is mostly implemented at the species level. However, genetic differentiation exists within this taxonomic level, putting genetically distinct populations at risk of local extinction. In seabirds, reproductive isolation is one of the principal drivers of genetic structure. In the Diablotin Black-capped Petrel *Pterodroma hasitata*, an endangered gadfly petrel endemic to the Caribbean, two phenotypes have been described: a smaller dark form and a heavier light form, which are genetically distinct. We hypothesized that color forms have a similar non-breeding distribution at sea but distinct nesting distributions.

**Location:** Western North Atlantic and northern Caribbean islands.

**Methods:** In May 2019, we captured 5 adult Black-capped Petrels of each phenotype at sea and equipped them with satellite transmitters. We used generalized linear mixed models to test the importance of phenotype on geographic distribution. Using kernel density estimations, we located use areas, quantified spatial overlap between forms, and assessed form-specific exposure to marine threats. Finally, we used tracking data to estimate the distribution and timing of nesting.

**Results:** Petrels were tracked for 11 – 255 d (mean = 102.1 d±74.2). During the non-breeding period, all individuals ranged from 28.4 – 43.0 degrees latitude. Phenotypes had significantly distinct non-breeding distributions. In the western North Atlantic, the dark form was exposed to more marine threats than the light form. We recorded two trips (1 individual of each form) to known breeding areas, with the light form initiating breeding 1.5 months before the dark form.

**Main conclusions:** Phenotypic differences in the Black-capped Petrel were linked to differences in nesting phenology, non-breeding marine distribution, and at-sea threat exposure. To sustain the species’ representation, redundancy, and resiliency in the light of environmental changes, it is likely that the evolutionary processes that resulted in genetic differentiation will also need to be conserved.

## INTRODUCTION

Whether due to financial, political, or logistical reasons, and despite growing support for ecosystem-based approaches (Pörtner et al. 2021), conservation is still implemented most frequently at the species level (IUCN 2008, Arponen 2012, Burfield et al. 2017). All species are not, however, homogeneous with respect to genetic structure, and spatial or temporal partitioning can occur within a species, often leading to phenotypic differentiation and, potentially, speciation (Frankham et al. 2002). In cases where phenotypic differentiation leads to speciation, this genetic differentiation coevolves with local adaptation to habitats which subsequently should lead to increased fitness. Therefore, identifying populations’ traits such as distribution, habitat use, or trophic niche is a necessary step to inform and prioritize conservation, particularly for species that may not demonstrate homogenous traits among subpopulations or morphs. A poor appreciation of genetic differentiation within a species and a lack of conservation actions focused at the population level could cause the reduction or loss of populations, resulting in a significant loss of genetic variation (Ennos et al. 2005). Thus, even when subspecies status is not warranted, populations that exhibit genetic differentiation may require adapted conservation actions (Gaston 2001, but see Winker 2010; Ennos et al. 2005; Danckwerts et al. 2021). Therefore, in addition to understanding a species’ representation (the existence of genetic and phenotypic diversity within a species) and redundancy (its capacity to persist despite the loss of a population), understanding the physical scales at which genetic differentiation occurs has direct implications for conservation.

In vagile animals, geographic distance between breeding populations was originally assumed to lead to isolation and hence was proposed as the main driver of genetic differentiation (Wright 1943). Studies of highly mobile taxa have demonstrated that this mechanism is complicated, however, by natural and anthropological features of the landscape (Bridle et al. 2004, Deiner et al. 2007), complex dispersal patterns within populations (Hellberg 2009, Burridge and Waters 2020), sociality and philopatry (Randall et al. 2010, Ribeiro et al. 2012), migratory behavior (Rolshausen et al. 2009, Burridge and Waters 2020), or combinations of these factors (Mancilla-Morales et al. 2020). In seabirds, which are some of the most highly vagile animals, isolation by distance among breeding sites is only a weak predictor of genetic structure among populations at diverse geographic scales (Friesen et al. 2007a). Seabirds are geographically constrained to a limited number of terrestrial nesting areas (e.g., available oceanic islands or coastal areas) during the breeding season, but most species travel long distances during the non-breeding season, sometimes to both hemispheres (Croxall et al. 2005, Shaffer et al. 2006, Alerstam et al. 2019). Thus genetic differentiation may result from (a) isolation due to processes such as non-breeding distributions (Friesen et al. 2007a, Friesen 2015), with birds using different areas less likely to mix at sea or to return to similar breeding colonies, (b) selection of distinct foraging locations during the breeding period (Wiley et al. 2012, Lombal et al. 2018, but see Lombal et al. 2020 for a discussion of inconsistencies in the effects of spatial segregation during foraging), or (c) differences in feeding strategies and niche partitioning (Ryan et al. 2014). In addition, seabirds tend to have high levels of colony philopatry; therefore, mechanisms of reproductive isolation, including temporal mechanisms (e.g., breeding allochrony in the sympatric *Oceanodroma castro* and *O. monteiro*: Monteiro and Furness 1998, Friesen et al. 2007b, Bolton et al. 2008), remain one of the principal drivers for accumulating genetic structure (Friesen et al. 2007a).

Our ability to investigate these potential drivers of isolation has improved with advances in the miniaturization of biologging devices (Hussey et al. 2015, Kays et al. 2015). Nevertheless, few tracking studies have been conducted with many species that are globally threatened or endangered (Bernard et al. 2021) or showing geopolitical bias in the location of breeding sites (Mott and Clarke 2018). In seabirds, tracking is commonly initiated on individuals captured at nest sites and subsequently used to map migratory paths or locate core use areas. However, tracking has also been used to attempt to locate unknown nesting areas of species captured on non-breeding grounds (Kanai et al. 2002, Rayner et al. 2020). Remote tracking technology is particularly well suited for seabirds captured at sea when there is no a priori assumption of possible nesting grounds, or when the potential nesting range is too broad (Rayner et al. 2015, Rayner et al. 2020). In addition, capturing seabirds at sea prior to or during the breeding season has the advantage of not predetermining which populations are sampled and birds can be tracked regardless of their breeding location. Given adequate spatiotemporal resolution, data gathered from such tracking studies can also be used to assess at-sea distributions and exposure to threats at sea (Hays et al. 2019). Exposure to marine threats (including but not limited to fisheries bycatch, overfishing, pollution, and attraction and collisions to ships and structures: Ronconi et al. 2015, Dias et al. 2019) can occur at different geographic scales. Macro-exposure is defined as the occurrence of the population of concern within the geographical area of interest (Burger et al. 2011) and is frequently calculated as spatiotemporal overlap between a population and the threats impacting it (Fischer et al. 2021, Pereira et al. 2021). Spatial overlap does not necessarily imply interaction with marine threats, but this parameter can be used as a proxy for potential exposure (Le Bot et al. 2018).

The Diablotin, or Black-capped Petrel *Pterodroma hasitata*, is an endangered gadfly petrel endemic to the Caribbean and occurs in waters of the western North Atlantic Ocean, Caribbean Sea, and Gulf of Mexico (Simons et al. 2013, Jodice et al. 2015, Jodice et al. 2021). The species is considered Endangered throughout its range (BirdLife International 2018) and is being considered by the U.S. Fish and Wildlife Service for listing as Threatened under the Endangered Species Act (U.S. Fish and Wildlife Service 2018). Two color forms have been described (dark and light, with intermediate phenotypes) that differ in size and by the amount of white plumage on the face, back of the neck, and underwing feathers (Howell and Patteson 2008) (Figure S1). The species has a fixed population structure, with a strong genetic divergence between the two forms, with light and intermediate forms belonging to a unit separate from the dark form (Manly et al. 2013). This phylogenetic structure suggests the existence of two distinct populations that are isolated geographically and/or temporally (Howell and Patteson 2008, Manly et al. 2013). The dark form begins the breeding period with visits to nesting grounds around mid-November; egg-laying occurs in mid-January, hatching in mid-March, and fledging around mid-June (Simons et al. 2013). Analysis of molt patterns suggest that the light form may be breeding 1 to 1.5 months earlier (Howell and Patteson 2008, Manly et al. 2013) (Figure S2). Observations at sea suggest that both forms use similar non-breeding areas (Howell and Patteson 2008, Simons et al. 2013) and no differences in foraging strategies have been described (see Haney 1987), so reproductive isolation has been described as the most probable mechanism for genetic differentiation (Haney 1987, Manly et al. 2013). Breeding populations are fragmented into five distinct breeding areas on the island of Hispaniola, with suspected additional breeding areas in Dominica and Cuba (Wheeler et al. 2021). To date, the dark form is more commonly observed at sea (Howell and Patteson 2008) and on land (from camera trapping, stranding, or captures at nest sites; Simons et al. 2013, Satgé et al. 2019, Wheeler et al. 2021). Observations of the light and intermediary forms have occurred primarily at sea (Howell and Patteson 2008). Light individuals have recently been confirmed to nest at breeding areas in the central Dominican Republic (E. Rupp, Grupo Jaragua Inc., oral communication, 2021) but no clear pattern of the breeding distribution of the two phenotypes has emerged. Therefore, to better understand the species’ biogeography and inform conservation, in the spring of 2019 we attempted to locate unknown nesting areas using satellite telemetry on individuals captured at sea. The objectives of this study were to: 1) assess the connectivity of Black-capped Petrels between their non-breeding foraging locations in the Gulf Stream and their breeding locations in the Caribbean; 2) evaluate differences in the nesting and non-breeding distributions of both forms; 3) assess differences in nesting phenology between forms; and 4) assess macro-scale exposure to marine threats in both forms. We hypothesized that color forms would have a similar non-breeding distribution, and thus similar exposure to marine threats, but that they would have separate nesting distributions.

## METHODS

### Fieldwork

At-sea captures occurred during May 2019 in Gulf Stream waters, ∼ 60 km southeast of Cape Hatteras, North Carolina, USA, an area where foraging Black-capped Petrels are commonly found during the non-breeding seasons (Simons et al. 2013, Jodice et al. 2015; Figure 1). We chose to capture birds during the boreal spring because individuals of the light form of Black-capped Petrel appear to be more common off Hatteras at this time of the year, whereas dark-form petrels appear to be more common during the late summer and fall (Howell and Patteson 2008). After sunrise, we located Black-capped Petrels from a ∼ 20-m research vessel, with the aid of chumming (a mixture of fish meal and shark liver oil). Upon detection of petrels, we deployed a metal cage (∼ 20 × 20 × 40 cm) secured to a drifting vinyl mooring buoy and containing blocks of frozen chum, and launched a ∼ 3-m motorized inflatable boat with two occupants, a pilot and a catcher. We positioned the inflatable boat upwind from the buoy, keeping the bow of the boat and the catcher facing downwind. We attempted to capture Black-capped Petrels that had flown upwind along the oil slick and approached within 10 – 15 m forward of the bow. We used a modified air-propelled whale tagger (ARTS Whale tagger, Restech, Norway) custom-fitted to launch four narrow PVC tubes (approx. 50 cm x 1.5 cm diameter; designed to float) attached to the corners of a 4 × 4 m mist net (adapted from Rayner et al. 2020). The net launcher was powered by compressed air from a dive tank. Upon capture, we transferred petrels to the research vessel for processing and transmitter deployment. This transfer lasted < 3 min.

**Figure 1.**
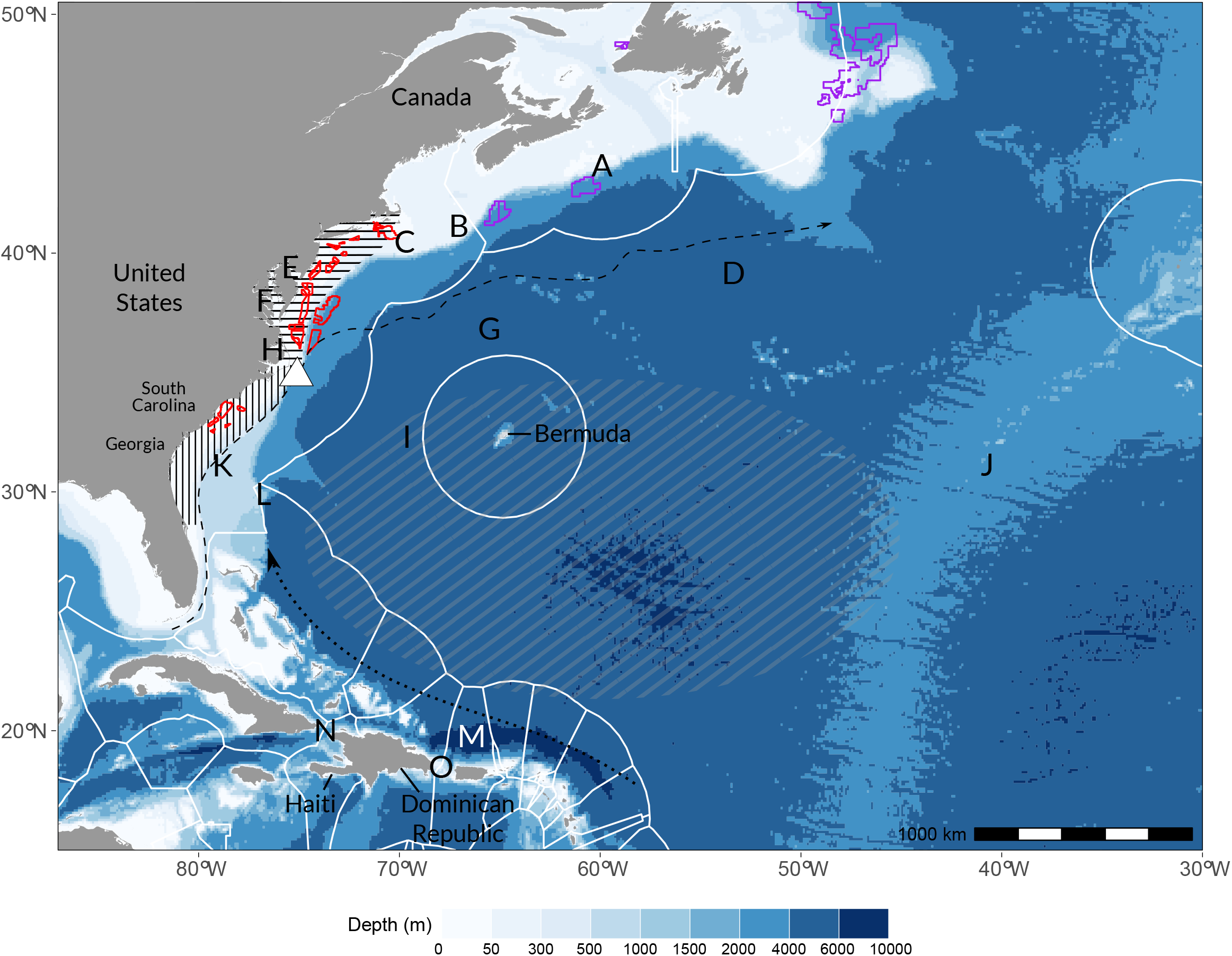
Study area. White triangle indicates capture location. Letters indicate geographic and oceanographic features. A: Banquereau Bank; B: Georges Bank; C: Nantucket Shoals; D: Sohm Plains; E: Delaware Bay; F: Chesapeake Bay; G: Caryn Seamount; H: Cape Hatteras; I: Hatteras Plains; J: Mid-Atlantic Ridge; K: Charleston Bump; L: Blake Spur; M: Puerto Rico Trench; N: Windward Passage; O: Mona Passage. Black lines indicate the general location of the western edge of the Gulf Stream (dashed) and of the Antilles Current (dotted). Black horizontal dashed area indicates the Middle Atlantic Bight, and black vertical dashed area the South Atlantic Bight. Grey dashed area locates the general extent of the Sargasso Sea. White polygons represent exclusive economic zones. Purple polygons locate petroleum leases, and red polygons offshore wind leases. (Mollweide equal-area projection).

After assessing captured petrels for general condition, we measured body mass (± 5 g), tarsus (± 0.1 mm), wing chord (± 1 mm), exposed culmen length (± 0.1 mm), and bill depth at gonys (± 0.1 mm). We banded petrels with individually numbered metal bands (U.S. Geological Survey Bird Banding Laboratory, Maryland, USA). We collected a few drops of blood from one metatarsal vein for genetic sexing. We photographed the birds’ profiles and upper- and under-wings, and classified them as dark, intermediate, or light forms. In addition, we assessed the age classes of the birds based on the shape and aspect of secondary flight feathers: we assumed that first-year juveniles had pointy feathers with a uniform aspect (due to limited wear and uniform growth), and immature—adults had worn secondaries with a varied aspect (due to scattered molting of flight feathers) (K. Sutherland, Univ. of North Carolina Wilmington, written communication, 2022). Finally, we recorded any molting of flight feathers. We used a t-test to compare each morphometric measurement between forms (for all statistical analyses, we grouped light and intermediate forms following Manly et al. 2013).

We deployed solar-powered PTTs (GT-5GS n = 8: GeoTrak Inc., North Carolina, USA, 5 g; 5g-Solar-PTT n = 2: Microwave Telemetry Inc., Maryland, USA, 5 g) on petrels whose body mass > 350 g. For the GeoTrak PTTs, we chose a duty cycle of 6 h on, 28 h off to benefit from the most extensive tracking time while optimizing battery usage; Microwave Telemetry PTTs were donated to the study with a pre-set duty cycle of 5 h on, 48 h off. All PTTs were custom-fitted with a base of marine-grade epoxy of ∼ 2 mm in thickness, in which 4 tubular channels were made perpendicular to the length of the PTT. Customized PTTs weighed 8.5 g (≤ 2.5 % of body mass of lightest petrel in Simons et al. 2013, Jodice et al. 2015, and Satgé et al. 2019). We deployed PTTs dorsally between the wings and centered laterally above the vertebrae, using four subcutaneous sutures and a small amount of glue (*sensu* Jodice et al. 2015). Before release, we placed equipped birds in a holding crate lined with a dry cloth towel until chest feathers were preened (∼ 20 minutes). Molecular sexing was performed at the Centro de Ecologia, Evolução e Alterações Ambientais, University of Lisbon, Portugal, following Fridolfsson and Ellegren (1999) with primers 2550F and 2718R. All animal handling was performed under Clemson University’s Animal Care and Use protocol AUP2019-033. Banding and PTT deployment were authorized by the USGS Bird Banding Lab (permit #22408).

### Spatial analysis of tracks

Tracking data were imported to Movebank.org *via* Service Argos. Argos applies a Kalman Filter model and provides error statistics and a quality class (3, 2, 1, 0, B, A, Z, in decreasing order of quality) for each estimated location. To improve the accuracy of Argos location estimates, we filtered locations and estimated the most probable “true” location using a continuous-time random walk state-space model (package *foieGras* in R, Jonsen et al. 2019). The mean duration between two subsequent Argos locations was 32 minutes therefore we used a time-step parameter of 30 minutes; we also included a maximum flight speed of 20 m/s as a model parameter. Fitting algorithms can estimate locations during periods when PTTs are off but the precision of fitted locations decreases during long periods without transmissions and unrealistic estimations may occur. Therefore, instead of filtering fitted locations to an arbitrary time period (e.g., 1 to 2 h from first or last Argos location during an on period), we selected fitted locations in on and off periods based on their spatial standard errors. Hence, we kept only fitted locations where the standard error for longitude and latitude was less than the 95^th^ percentile of the error radius of location classes 0-3 (10.3 km, Table S1).

We restricted analysis of location data to a period of time during which most birds were tracked, removing data from individuals tracked for less than 100 days (n = 4), and removing outlying data from a single individual travelling eastward towards the mid-Atlantic ridge, a behavior associated with vagrancy (Simons et al. 2013). This filtering resulted in a study time frame of 14 May to 25 August 2019, included 70% of all locations, and included 7 individual birds (dark: n = 3 dark; light: n = 4). After determining that the distributions of latitude and longitude in each phenotype were not normally distributed (Shapiro Wilk normality test with p<0.005 for both groups), we compared longitudinal and latitudinal distributions between dark and light forms using Wilcoxon rank sum tests. We estimated the magnitude of difference between groups by calculating Cohen’s *d* effect size (function *cohen*.*d* in package *effsize* in R, Torchiano 2020). Cohen’s *d* is defined as the difference between two means divided by a standard deviation for the data. Cohen (1988) suggests quantitative descriptors as follows: *d* ≤0.2 represents a small effect, 0.2< *d* ≤0.5 a moderate effect, and *d* >0.5 a large effect (but see Sawilowsky 2009 for a discussion of descriptors). Because of the imbalanced number of individuals of each sex (n = 3 females and n = 7 males), we did not assess the effect of sex on petrel distribution.

The use of foraging areas may also depend on the time of year, an individual’s phenology, or on individual variability. Therefore, to compare latitudinal and longitudinal distributions with phenotype, we used generalized linear mixed models with a gamma distribution and inverse link, and included date and individual as random effects (function *glm* in package *stats* in R). To avoid any variations within individuals within dates, we used the latitude and longitude of the mean daily location for each individual. We did not include longitude as a fixed effect in the model of latitude nor did we include latitude as a fixed effect in the model of longitude because the distribution of Black-capped Petrel is geographically constrained by the continental coastline of the eastern United States (roughly a southwest-to-northeast diagonal) and latitude and longitude were correlated (Spearman’s correlation index ρ = 0.71, *p* <0.005). We estimated the importance of phenotype as the absolute value of the t-statistic for each model (package *caret* in R, Kuhn 2020).

We calculated utilization distributions (UD) for both phenotypes using kernel density estimations in package *adehabitat* in R (Calenge 2006; with smoothing parameter h = 0.3 and grid = 1,000). Within a form, all individuals were grouped. We estimated the amount of spatial overlap among home range (90% UD) and core (50% UD) areas between the two forms. We also quantified the extent of spatial overlap in the home range and core areas of both forms using Bhattacharyya’s affinity (BA). BA is a function of the product of the utilization distributions of two populations (here dark and light forms) that assumes each population uses space independently from the other (Fieberg & Kochanny 2005). A BA value = 0 indicates no overlap while a BA value = 1 indicates complete overlap. Finally, we estimated overlap of the two populations with exclusive economic zones (EEZ; VLIZ 2019) and marine ecoregions (Spalding et al. 2007) by calculating the proportion of the 50% and 90% UDs of each phenotype within EEZs and international waters, and ecoregions and high seas, respectively. We obtained shapefiles for EEZs and ecoregions from marineecoregions.org.

### Colony visits

Black-capped Petrels access breeding sites after dusk (Jodice et al. 2015) and, when at the colony, are active at night but spend daylight hours underground (Simons et al. 2013). Thus, when at breeding sites, solar-powered PTTs cannot adequately communicate with satellites and cannot adequately recharge. We assumed that locations near known breeding sites were likely to indicate some form of breeding activity (e.g., prospecting or nest initiation). We also assumed that breeding activity (e.g., occupancy of a burrow) would lead to gaps in communications between the satellite tag and satellite system and to low voltage levels of the satellite tags. Therefore, for locations within one month before and one month after a suspected visit to a breeding site, we: 1) calculated the distance to the nearest breeding site with function *distGeo* in package *geosphere* in R (Hijmans 2019); 2) used raw PTT data (location and metadata from Service Argos) to assess any gap in satellite communication; and 3) compared mean daily voltage with the overall mean voltage during that period.

### Overlap with habitat features and marine-based threats

The Black-capped Petrel is considered to be strongly associated with the Gulf Stream off the southeast coast of the United States (Haney 1987, Jodice et al. 2015, Winship et al. 2018), so we examined differences between dark and light forms of Black-capped Petrels in their use of two key habitat features: ocean depth and sea surface temperature (SST). We did so by calculating spatial statistics for ocean depth (raster ETOPO1; 1 arc minute; Amante and Eakins 2009) and SST (raster HYCOM; 0.08 arc degree; May – September 2019; Cummings and Smedstad 2013) for all raster cells overlapping with 50% and 90% UDs of each form. We also compared differences in the distributions of environmental values (i.e. the depth and SST raster values extracted from all cells overlapping UDs) between phenotypes using Wilcoxon rank sum tests.

We then sought to quantify differences between dark and light forms of Black-capped Petrels in terms of their macro-exposure to marine-based threats in the western North Atlantic. We defined macro-scale exposure as the level of occurrence of a threat within the home range and core areas of each phenotype, *sensu* Burger et al. (2011) and Waggit and Scott (2014). We included assessments of mercury, plastics, fisheries, marine energy, and shipping. We assessed the potential exposure of Black-capped Petrel to mercury using, as a proxy, a model of the spatial concentration of total mercury developed by Zhang et al. (2014). We used a raster file of modelled present-day mercury concentrations in the mixed layer (i.e. from the ocean surface to 50 m depth; Figure 3a in Zhang et al. 2014) with a resolution of 1 × 1° (between 80 × 100 km and 100 × 122 km, depending on latitude). We quantified potential exposure to plastics using, as a proxy, global models of the spatial distribution of micro-plastics (van Sebille et al. 2015). For each 1 × 1° cell, we averaged concentrations of micro-plastics (g/km^2^) predicted by the Maximenko, Lebreton, and van Sebille models (Figure 3 in van Sebille et al. 2015). We obtained data of daily commercial fishing effort as fishing hours at 0.1° cell resolution (between 8 × 9.5 km and 10 × 12.5 km depending on latitude) from Global Fishing Watch (http://globalfishingwatch.org/; accessed 1 July 2021). Global Fishing Watch combines satellite tracking of commercial fishing vessels equipped with automatic identification systems and convolutional neural networks to classify the activity of vessels larger than 15 m as fishing or not fishing (Kroodsma et al. 2018). This dataset represents >50−70% of the global fishing effort (n > 70 000 vessels; Kroodsma et al. 2018). We summed all available daily effort data from May to August 2019, which corresponded to the period when most birds were tracked. To assess overlap with marine traffic, we obtained data of vessel transit from the U.S. Office for Coastal management through the U.S. Marine Cadastre (https://www.fisheries.noaa.gov/inport/item/55365; accessed 1 July 2021), as transit counts at 100 m cell resolution. This dataset represents annual vessel transits counts collected from automatic identification systems, where a single transit count corresponds to each time a vessel track passes through, starts, or stops within a grid cell (Office for Coastal Management 2021). We aggregated raster data attributed to cargo, tanker, and all other vessel types (which includes fishing, passenger, pleasure craft and sailing, and tug and towing), for 2017 (the latest available year) at a resolution of 10 × 10 km. We obtained spatial datasets of active oil and gas exploration areas from the Canada-Nova Scotia Offshore Petroleum Board (https://www.cnsopb.ns.ca/resource-library/maps-and-coordinates; accessed 1 November 2021) and the Canada-Newfoundland and Labrador Offshore Petroleum Board (https://www.cnlopb.ca/information/shapefiles/; accessed 1 November 2021), and of wind energy production from the U.S. Bureau of Ocean Energy Management (https://www.boem.gov/renewable-energy/mapping-and-data; accessed 1 May 2022). To simplify analyses, we merged individual lease plots occurring within a U.S. state, and occurring in Newfoundland and Labrador (Canadian hydrocarbon areas) into project-scale lease areas (i.e. one lease area per administrative area). We also merged lease areas occurring in the U.S. states of Rhode Island and Massachusetts into a single lease unit.

For each of the above attributes, we calculated spatial statistics from the values of all raster cells overlapping with 50% and 90% UDs: mean, minimum, maximum, standard deviation, and interquartile range (IQR). For fishing effort and ship transit, we also calculated the percentage of cells exposed to the threat, and the sum of fishing effort and ship transit in both UDs. Shapiro-Wilk tests indicated that habitat features and threats followed non-parametric distributions; therefore we assessed differences in the numeric distribution of threat values between phenotypes using Wilcoxon rank sum tests. We estimated the magnitude of difference between groups by calculating Cohen’s *d* effect size. We assessed exposure to marine energy production by calculating the proportion of the 50% and 90% UDs of each phenotype within the footprint of hydrocarbon and wind leases. We also calculated the shortest distance between a tracking location and a lease area.

All statistical and spatial analyses were performed in R version 3.6.3 (R Core Team 2020).

## RESULTS

Capture attempts occurred on 08, 09, 11, and 14 May 2019, within a 25-km radius of 34.78°N, 75.33°W, along the continental slope of the eastern United States and the western edge of the Gulf Stream. Capture effort ranged from 3.0 – 6.5 h on each of the four capture days. We captured 2 birds on 08 May, 4 birds on 09 May, 0 birds on 11 May, and 4 birds on 14 May. Sea state varied from Beaufort 2 to 5 among the four capture days with the lowest sea state occurring on 11 May when no birds were captured. Approximately 50 % of attempts resulted in a successful capture. Sex (n = 3 females, n = 7 males) and morphometric data are summarized in Table 1. Of the 10 Black-capped Petrels we instrumented, we classified five birds as dark-forms, four as light-forms, and one as intermediate (Table 1). None of the petrels were first-year juveniles but we could not assess age further and separate between immatures (1-4 years) and adults (>4 years; Simons et al. 2013). Morphometrics did not differ between sexes (*p* > 0.05 for all tests) (Table 1). Dark forms appeared smaller than light and intermediate forms in all metrics, but the differences were only significant for wing cord (mean _Dark (5)_ = 289.3 mm vs mean_Light (5)_ = 303.2; *p* < 0.05; *t* = -3.6) and culmen length (mean_Dark (5)_ = 32.5 mm vs mean_Light (5)_ = 35.4; *p* < 0.05; *t* = - 5.0) (Table 1; *p* > 0.05 for all other comparisons). All light and intermediate forms were molting at least two flight feathers but none of the dark forms were molting (Table 1). Deployed PTTs ranged from 1.85 % – 2.30 % of body mass (mean: 2.16 %). Processing time ranged from 13 – 23 min (mean: 18 min) per individual.

**Table 1.** Phenotype and morphometrics of Black-capped Petrels captured off Cape Hatteras, North Carolina, USA, May 2019.

| Bird ID | Capture date | Sex | Mass<br>(g) | Tarsus<br>(mm) | Culmen<br>(mm) | Bill depth<br>(mm) | Wing cord<br>(mm) | Molt <sup>a</sup> |
| --- | --- | --- | --- | --- | --- | --- | --- | --- |
| **Dark forms** |  |  |  |  |  |  |  |  |
| **469** | 8 May 2019 | M | 370 | 39.5 | 32.6 | 14.4 | 295 | N |
| **464** | 8 May 2019 | F | 390 | 38.7 | 32.5 | 13.7 | 292 | N |
| **466** | 9 May 2019 | M | 380 | 41.7 | 31.5 | 14.0 | 290 | N |
| **442** | 14 May 2019 | M | 380 | 39.2 | 34.8 | 13.6 | 280 | N |
| **441** | 14 May 2019 | M | 380 | 39.0 | 32.1 | 14.0 | 287 | N |
| **Average** |  |  | 380.0 | 39.6 | 32.7** | 13.9 | 288.8** |  |
| **SD** |  |  | 6.32 | 1.07 | 1.12 | 0.28 | 5.11 |  |
| **Light and intermediate forms** |  |  |  |  |  |  |  |  |
| **468** | 9 May 2019 | M | 420 | 39.1 | 35.3 | 14.3 | 300 | P1-2 |
| **462** | 9 May 2019 | M | 375 | 40.2 | 35.0 | 14.0 | 299 | P3-4, S3 |
| **467<sup>b</sup>** | 9 May 2019 | M | 410 | 40.6 | 34.6 | 13.6 | 297 | P2-4 |
| **465** | 14 May 2019 | F | 390 | 41.1 | 36.0 | 14.1 | 315 | P2-3 |
| **463** | 14 May 2019 | F | 460 | 41.4 | 36.0 | 14.4 | 305 | P1-2 |
| **Average** |  |  | 411.0 | 40.5 | 35.4** | 14.1 | 303.2** |  |
| **SD** |  |  | 29.05 | 0.80 | 0.55 | 0.28 | 6.46 |  |

<sup>a</sup> All molting feathers were in sheaths and growing. P = primary and S = secondary flight feathers, and N = none.
<sup>b</sup> Intermediate form.
\*\*Significant difference between dark and light forms at $p < 0.05$ .

Black-capped Petrels were tracked for 11 – 255 d ± 74.2 (mean: 102.1 d; median: 108.5 d; Table 2 and Figure S3), resulting in 1,021 bird-tracking days. We tallied 4,656 locations and 73% of all locations were calculated from four (or more) Argos messages (location classes 0 – 3; Table S1). Of all locations, 37% were accurate to < 1,500 m, and 84 % were accurate to < 10 km. The maximum error radius for locations of class LC 0 (defined by an error radius > 1,500 m but with no upper limit) was 216 km (Table S1), but the 95^th^ percentile for this class was at 17.5 km. Refitted locations (n = 4142) had a mean precision of 5.1 km ± 2.5 for longitude and 4.4 km ± 2.4 for latitude.

**Table 2.** Summary of tracking period and geographic range of Black-capped Petrels tracked from May 2019 – January 2020.

| **Bird ID <sup>a</sup>** | **First transmission date** | **Last transmission date** | **No. of tracking days** | **Latitudinal range (°)** | **Longitudinal range (°)** | **General areas used** |
| --- | --- | --- | --- | --- | --- | --- |
| **469** | 2019-05-08 | 2019-05-25 | 17 | 30.3 – 35.8 | -80.2 – -74.5 | South Atlantic Bight: continental shelf |
| **464** | 2019-05-08 | 2019-08-25 | 109 | 29.4 – 38.5 | -78.2 – -70.3 | Off Virginia: high seas |
| 468 | 2019-05-09 | 2019-08-31 | 114 | 35.6 – 40.4 | -75.4 – -62.8 | Off Delaware – New Jersey: high seas |
| **466** | 2019-05-09 | 2019-05-20 | 11 | 31.5 – 37.4 | -78.9 – -72.8 | South Atlantic Bight: continental shelf |
| 462 | 2019-05-09 | 2019-11-14 | 189 | 17.4 – 43.1 | -75.9 – -56.3 | Off Delaware – Connecticut: high seas; Dominican Republic |
| 467 | 2019-05-09 | 2019-09-09 | 123 | 34.7 – 40.6 | -75.4 – -64.0 | Off Virginia – New Jersey: high seas |
| **442** | 2019-05-14 | 2020-01-24 | 255 | 17.3 – 38.0 | -80.1 – -64.1 | Off South Carolina, North Carolina: continental slope; Haiti |
| **441** | 2019-05-14 | 2019-08-30 | 108 | 30.4 – 38.2 | -79.3 – -65.8 | Off Georgia – North Carolina: continental slope; off Virginia: high seas |
| 465 | 2019-05-14 | 2019-08-01 | 79 | 31.1 – 39.8 | -77.2 – -50.5 | Off Delaware, Maryland: high seas; mid-Atlantic |
| 463 | 2019-05-14 | 2019-05-30 | 16 | 32.1 – 34.6 | -77.4 – -74.2 | Off North Carolina: continental slope |

<sup>a</sup> Bold lettering indicates dark phenotypes.

### Spatial analysis

Seven individuals were included in the spatial analysis: 3 dark forms and 4 light forms. With the exception of two trips to the vicinity of Hispaniola, all individuals ranged from 28.4° – 43.0° latitude (Figure 2 and Figure S4, Table 2). Tracked petrels remained west of -60° longitude, except for one individual that was last located at approximately -50° longitude (ID 465). Locations were concentrated along the western wall of the Gulf Stream, over the outer continental shelf of the United States. Petrels were located in the interior of the Stream itself but the eastern wall of the Stream and the Sargasso Sea remained largely unused. One individual (ID 462) utilized the western half of the Sohm Plains, travelling as far north as the Canadian continental shelf off Banquereau Bank.

**Figure 2.**
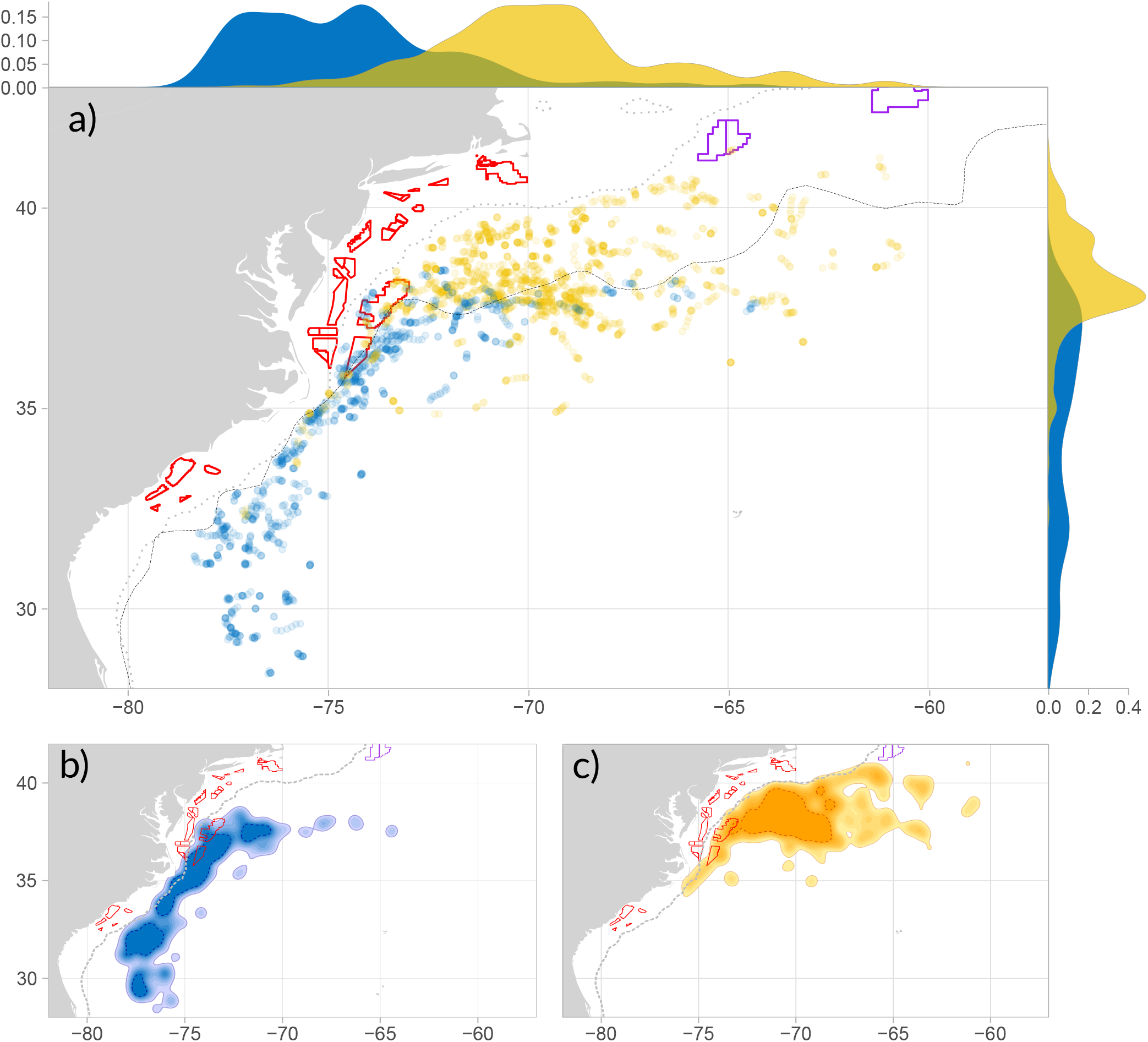
Distribution of tracked Black-capped Petrels in the western North Atlantic. For all panels, blue represents dark forms and yellow represents light forms. (a) Locations of all Black-capped Petrels tracked between May 2019-January 2020 (outliers were removed) with frequency distributions of longitude and latitude of dark form birds and light form birds overlaid on the top and right border of the panel. (b) Utilization distributions of dark Black-capped Petrels. (c) Utilization distributions of light Black-capped Petrels. For (b) and (c), petrels were tracked from May 2019 – August 2019; colored dashed lines indicate home ranges (95% UD: thin line) and core area (50% UD: thick line). Solid grey line shows the general location of the western edge of the Gulf Stream. Dotted grey line indicates the -250-m isobath. Grey polygons represent exclusive economic zones. Purple polygons locate petroleum leases, and red polygons offshore wind leases. (Mollweide equal-area projection).

Overall, the dark form (core area: 88,058 km^2^; home range: 292,472km^2^) occupied a more extensive core area than the light form (core area: 81,145 km^2^; home range: 355,915 km^2^) but had a more limited home range. Dark and light forms had significantly distinct distributions (Table 3 and Figure 2). The light form had a narrower and significantly more northerly latitudinal range than the dark form, and utilized deeper waters in its core area (Table 3, Figures 2, S5-7). In addition, both forms showed extended longitudinal ranges, with the dark form having a significantly more westerly distribution (Table 3 and Figure 2). The dark morph utilized significantly warmer waters than the light form in its core area (Table 3, Figures S6 and S7). Phenotype was a strong predictor of both latitude (|t| = 19.4, *p* < 0.005) and longitude (|t| =18.8, *p* < 0.005).

**Table 3.** **Characteristics of geographic distribution, environmental variables, and threat exposure for Black-capped Petrels tracked from May 2019 – August 2019, grouped by phenotype (n = 3 dark, n = 4 light). For latitude/longitude, n = number of locations. For core area (50% UD) and home range (90%UD), sample size n = number of raster cells in UD). Percentage of cells exposed to the threat (Prop.), sum of fishing effort and vessel counts (Sum), mean, range (Min. = minimum value, Max. = maximum value), standard deviation, interquartile range (IQR), and p-value of Wilcoxon rank sum tests and Cohen’s *d* effect size are provided. Bold lettering denotes significant difference between light and dark forms and moderate or large effects. Effect size: * small (d ≤0.2), ** moderate (0.2< *d* ≤0.5), *** large (d >0.5). UD = Utilization distribution**.

| Dark |  |  |  |  |  |  |  |  |  | Light |  |  |  |  |  |  |  |  |  |
| --- | --- | --- | --- | --- | --- | --- | --- | --- | --- | --- | --- | --- | --- | --- | --- | --- | --- | --- | --- |
| Variable | UD | n | Prop | Sum | Mean | Min. | Max. | SD | IQR | n | Prop. | Sum | Mean | Min. | Max. | SD | IQR | p-value | Effect size |
| Latitude (°) | - | 1133 | - | - | 34.50 | 28.40 | 38.40 | 2.73 | 4.72 | 1771 | - | - | 38.1 | 32.30 | 41.40 | 1.24 | 1.38 | <0.005 | 1.86*** |
| Longitude (°) | - | 1133 | - | - | -74.4 | -78.4 | -64.3 | 2.62 | 3.21 | 1771 | - | - | -69.6 | -77.10 | -60.70 | 2.82 | 2.86 | <0.005 | 1.78*** |
| Depth (m) | 50 | 26755 | - | - | -2090.8 | -4152 | -6 | 1124.3 | 2111 | 25870 | - | - | -3428.6 | -5030 | -1934 | 663.6 | 1149 | <0.005 | 1.44*** |
|  | 90 | 88858 | - | - | -2639.9 | -5143 | -5 | 1486.6 | 2752 | 114897 | - | - | -3505.9 | -5670 | -5 | 1338.1 | 1923 | <0.005 | 0.61*** |
| SST (°C) | 50 | 1072 | - | - | 27.41 | 23.81 | 28.61 | 1.10 | 1.34 | 1036 | - | - | 24.43 | 21.53 | 26.93 | 1.46 | 2.79 | <0.005 | 2.30*** |
|  | 90 | 3554 | - | - | 26.92 | 22.28 | 28.68 | 1.46 | 1.83 | 4595 | - | - | 24.13 | 14.87 | 28.24 | 2.31 | 3.51 | <0.005 | 1.40*** |
| Hg (x10<sup>-10</sup> mol/m3) | 50 | 6 | - | - | 9.75 | 9.51 | 9.96 | 0.16 | 0.13 | 7 | - | - | 9.44 | 9.00 | 10.20 | 0.40 | 0.40 | 0.074 | 0.98*** |
|  | 90 | 28 | - | - | 9.55 | 8.55 | 1.04 | 0.40 | 0.42 | 38 | - | - | 9.10 | 8.03 | 10.20 | 0.56 | 0.81 | <0.005 | 0.90*** |
| Plastic (g/km<sup>2</sup>) | 50 | 6 | - | - | 149.37 | 86.62 | 205.85 | 44.75 | 59.21 | 7 | - | - | 132.99 | 82.02 | 198.08 | 37.86 | 37.44 | 0.43 | 0.40** |
|  | 90 | 28 | - | - | 247.55 | 85.71 | 730.80 | 152.95 | 154.77 | 38 | - | - | 154.92 | 46.30 | 386.46 | 74.54 | 79.41 | <0.005 | 0.81*** |
| Fishing effort (h) | 50 | 113 | 13.0 | 1866.61 | 16.52 | 0.18 | 197.21 | 29.64 | 15.60 | 20 | 2.4 | 149.48 | 7.47 | 0.61 | 23.83 | 8.04 | 7.66 | 0.600 | 0.33* |
|  | 90 | 299 | 10.4 | 3693.79 | 12.35 | 0.18 | 197.21 | 22.30 | 12.71 | 510 | 13.7 | 22323.28 | 43.77 | 0.15 | 1987.09 | 138.88 | 38.65 | <0.005 | 0.28** |
| Ship traffic (count) |  |  |  |  |  |  |  |  |  |  |  |  |  |  |  |  |  |  |  |
| Cargo | 50 | 1271 | 97.3 | 182346 | 143.47 | 0.04 | 1910.99 | 258.54 | 108.42 | 1316 | 99.7 | 21554 | 16.38 | 0.04 | 150.11 | 15.76 | 14.84 | <0.005 | 0.70*** |
|  | 90 | 4164 | 96.5 | 371888 | 89.31 | 0.03 | 1910.99 | 201.21 | 59.94 | 5173 | 88.5 | 225726 | 43.64 | 0.01 | 1910.99 | 145.64 | 23.90 | <0.005 | 0.26** |
| Tanker | 50 | 1245 | 95.3 | 32711 | 26.27 | 0.07 | 166.32 | 29.47 | 27.91 | 1240 | 93.9 | 10026 | 8.09 | 0.02 | 63.71 | 7.81 | 8.56 | <0.005 | 0.84*** |
|  | 90 | 3922 | 90.9 | 69277 | 17.66 | 0.01 | 195.00 | 24.28 | 17.71 | 4508 | 77.2 | 56786 | 12.60 | 0.01 | 166.32 | 19.82 | 13.28 | <0.005 | 0.23** |
| Other | 50 | 1288 | 98.6 | 305290 | 237.03 | 0.04 | 2701.45 | 409.81 | 175.70 | 1320 | 100.0 | 36676 | 27.78 | 0.09 | 234.70 | 26.12 | 26.65 | <0.005 | 0.73*** |
|  | 90 | 4253 | 98.5 | 650198 | 152.88 | 0.01 | 4193.95 | 334.49 | 95.50 | 5400 | 92.4 | 429026 | 79.45 | 0.02 | 2883.33 | 245.20 | 44.55 | <0.005 | 0.25** |

Bhattacharyya’s affinities indicated limited overlap in home ranges (BA = 0.29) and almost no overlap in core areas (BA = 0.03) of dark and light forms. The area of overlap corresponded to 6.13% and 6.65% (50% UD), and 35.57 % and 29.23 % (90% UD) of the areas used by dark and light forms respectively (Table S2).

In the western North Atlantic, Black-capped Petrels occurred in the EEZ of three countries, and international waters (Table 4 and Figure S8). U.S. waters accounted for most of the core areas (dark form: 99.1%; light form: 74.2%) and home ranges (dark form: 77.3% light form: 55.7%). The core areas (dark form: 0.9%; light form: 25.8%), and home ranges of both forms (dark form: 15.3%; light form: 42.4%) overlapped with international waters. The Canadian EEZ was utilized by one individual of the light form (home range: 1.9%). The home range of the dark form overlapped with the Bahamian EEZ (7.4%).

**Table 4.** **Area and proportion of core area (50% UD) and home range (90% UD) of dark and light forms of Black-capped Petrels tracked from May 2019 – August 2019 overlapping with exclusive economic zones of the western North Atlantic. INT: International waters. UD = Utilization distribution**.

| UD | Dark |  |  | Light |  |  |
| --- | --- | --- | --- | --- | --- | --- |
|  | EEZ | Area (km<sup>2</sup>) | Proportion in EEZ (%) | EEZ | Area (km<sup>2</sup>) | Proportion in EEZ (%) |
| 50 | USA | 87243 | 99.1 | USA | 60231 | 74.2 |
|  | INT | 787 | 0.9 | INT | 20914 | 25.8 |
| 90 | USA | 226172 | 77.3 | USA | 198362 | 55.7 |
|  | INT | 44669 | 15.3 | INT | 150883 | 42.4 |
|  | Bahamas | 21632 | 7.4 | Canada | 6670 | 1.9 |

Black-capped Petrels occurred in six marine ecoregions (including high seas) (Table 5, Figure S9). The dark form was most present in the Carolinian (50.4% of core area and 39.2% of home range) and Virginian regions (36.2% of core area and 28.3% of home range). The light form was mostly limited to the Virginian region (70.7% of core area and 39.7% of home range) and high seas (29.3% of core area and 47.9% of home range). The dark form made incursions into the high seas (5.9% of core area and 20.9% of home range) and Bahamian region (7.5% of core area and 11.6% of home range). In its home range, the light form seldom utilized the Gulf of Maine/Bay of Fundy (8.6%), Carolinian (3.7%), and Scotian Shelf regions (0.1%).

**Table 5.** **Area and proportion of core area (50% UD) and home range (90% UD) of dark and light forms of Black-capped Petrels tracked from May 2019 – August 2019 within marine ecoregions of the western North Atlantic. UD = Utilization distribution**.

| UD | Dark |  |  | Light |  |  |
| --- | --- | --- | --- | --- | --- | --- |
|  | Ecoregion | Area (km<sup>2</sup>) | Proportion in ecoregion (%) | Ecoregion | Area (km<sup>2</sup>) | Proportion in ecoregion (%) |
| 50 | Carolinian | 44392 | 50.4 | Virginian | 57374 | 70.7 |
|  | Virginian | 31899 | 36.2 | High Seas | 23771 | 29.3 |
|  | Bahamian | 6589 | 7.5 |  |  |  |
|  | High Seas | 5178 | 5.9 |  |  |  |
| 90 | Carolinian | 114745 | 39.2 | High Seas | 170482 | 47.9 |
|  | Virginian | 82620 | 28.3 | Virginian | 141397 | 39.7 |
|  | High Seas | 61129 | 20.9 | GoM/BoF* | 30617 | 8.6 |
|  | Bahamian | 33979 | 11.6 | Carolinian | 13058 | 3.7 |
|  |  |  |  | Scotian Shelf | 363 | 0.1 |

\* Gulf of Maine/Bay of Fundy

### Visits to breeding sites

Two PTTs were still transmitting at the onset of the breeding season in September 2019: ID 442 (male dark form) and ID 462 (male light form). Both individuals made southerly trips to the Caribbean (Figure 3). Based on a combination of locations, location error classes, battery voltage and satellite communication schedules (which can be used to infer burrow occupancy) we infer that ID 442 was at a breeding site during 9 – 22 November, and 29 November – 3 December, and that ID 462 was at a breeding site during 2 – 8 October and 9 – 15 October. It remains inconclusive whether ID 442 visited a breeding site between 19 – 21 December. Details can be found in the Supporting online information..

**Figure 3.**
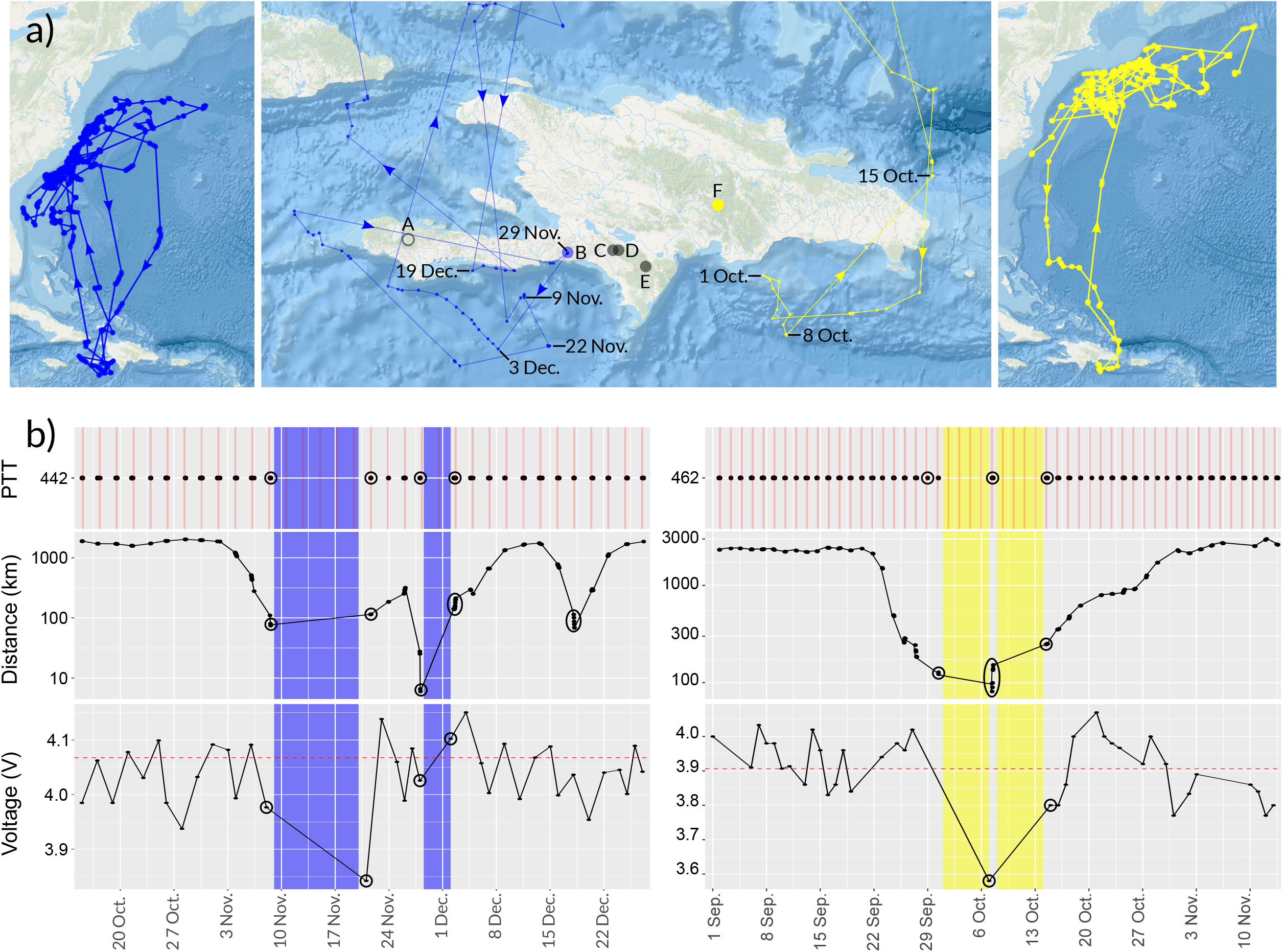
Movements and accessory tracking information of two Black-capped Petrels tracked to the vicinity of known breeding sites on Hispaniola, September-December 2019. For all panels, blue represents dark-form petrel ID 442, and yellow represents light-form petrel ID 462. (a) Left and right panels: movements of both tracked petrels for the entire tracking period (ID 442: 14 May 2019 to 24 January 2020; ID 462: 5 May to 14 November 2019). Center panel: movements of both tracked petrels in close proximity of Hispaniola. Dates represent dates when petrels were closest to a known breeding site. Circles represent known (filled) or suspected (outlined) breeding sites on Hispaniola: A: Pic Macaya, Haiti; B: La Visite Haiti; C: Morne Vincent, Haiti; D: Loma del Toro, Dominican Republic; E: Loma Quemada, Dominican Republic; and F: Valle Nuevo, Dominican Republic. The blue circle represents the most likely breeding site visited by petrel ID 442, and the yellow circle the most likely breeding site visited by ID 462, where nests of light-form petrels were located following this study. (b) For all panels, black ovals represent dates shown on the center map in (a); shaded rectangles show periods of suspected visits to breeding sites; left panels show accessory data for ID 442; and right panels represent accessory data for ID 462.Top panels: dates of communications between PTTs and satellites. Red bars represent scheduled communications, and black dots represent actual communications. Middle panels: distance to most likely breeding site. Bottom panels: battery levels of PTTs. Dashed red lines represent mean battery level for the duration of the period shown.

### Marine threats

Global mercury models showed that high levels of mercury were present in the mixed layer of the western North Atlantic (Zhang et al. 2014). Highest levels within the region were concentrated in the Gulf of Saint Lawrence and the Gulf of Maine, two areas not used by Black-capped Petrels (Figure S6c). Yet, petrels in our study were exposed to concentrations in the 90^th^ quantile of global levels in both their core areas and home ranges (dark: mean_Hg_ = 9.75 × 10^-10^ mol/m^3^; light: mean_Hg_ = 9.44 × 10^-10^ mol/m^3^) and home ranges (dark: mean_Hg_ = 9.55 × 10^-10^ mol/m^3^; light: mean_Hg_ = 9.10 × 10^-10^ mol/m^3^; 90th quantile: 9.17 × 10^-10^ mol/m^3^) (Table 3 and Figure 4). At the home range level, the dark form was significantly more exposed to mercury than the light form, with a large effect size (*p* = 0.001, |d| = 0.90). Differences between forms in the 50% UD, or between UDs within forms, were not significant (*p* > 0.005).

**Figure 4.**
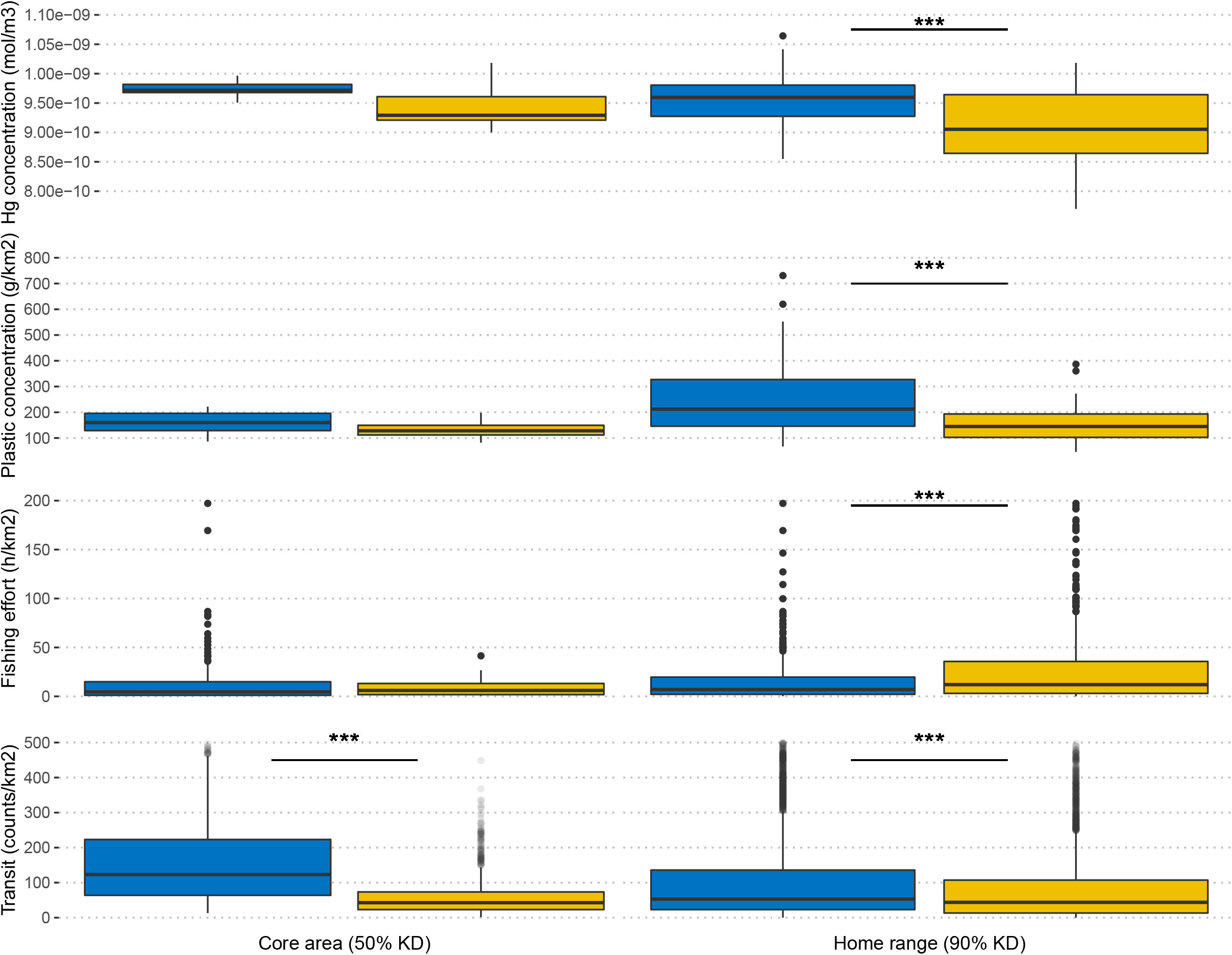
Distribution of values of threat exposure in core area and home range of Black-capped Petrels tracked from May 2019 – August 2019. Blue represents dark forms and yellow represents light forms. P-value of Wilcoxon sum rank test: *** <0.005. Boxes depict the median and quartiles, whiskers depict the 5th and 95th percentiles, and circles depict data beyond the 5th and 95th.

In the western North Atlantic, the highest levels of micro-plastic were concentrated in the Atlantic gyre of the Sargasso Sea (Figure S6d), an area not used by petrels in our study, although lower levels did overlap with use areas. At the home range level, the dark form was significantly more exposed to micro-plastic than the light form (dark; mean_Plastic_ = 247.55 g/km^2^; light: mean_Plastic_ = 154.92 g/km^2^),, with a large effect size (*p* < 0.005, |d| = 0.88) (Table 3, Figure 4). Differences between forms in the 50% UD, or between UDs within forms, were not significant (*p* > 0.05).

High commercial fishing effort was present in the U.S. and Canadian EEZ, with highest effort located on the continental plateau and shelf of the Georges Bank, the Nantucket Shoals, and locally off Delaware Bay (Figure S6e). Effort was generally limited to neritic and semi-pelagic waters (from the coastline to the bottom of the continental shelf), though some effort was recorded in the Sohm plains and southeastern parts of the Sargasso Sea. Petrels were exposed to fisheries in a limited portion of their distribution, in a strip of waters located along the continental shelf from Georges Bank to Hatteras, and on the continental plateau off the U.S. states of South Carolina and Georgia. Fishing effort occurred within 13.0 % and 2.4% of the core areas, and 10.4% and 13.7% of the home ranges of dark and light forms, respectively (Table 3). In the dark form, average fishing effort equated to 16.5 fishing hours per 0.1°cell in the core area, and 12.4 fishing hours per cell in the home range. Summed fishing effort equated to 1,866.6 fishing hours (50% UD) and 3,693.8 fishing hours (90% UD). In the light form, average fishing effort equated to 7.5 fishing hours per 0.1°cell in the core area, and 43.8 fishing hours per cell in the home range. Summed fishing effort equated to 149.5 fishing hours (50% UD) and 22,323.3 fishing hours (90% UD). At the home range level, the light form was significantly more exposed to fisheries than the dark, with a moderate effect size (*p* < 0.005, |d| = 0.24) (Table 3 and Figure 4).

High levels of shipping activity occurred in the study area (Figure S6f), with the highest densities of vessels localized along the shipping routes linking major U.S. shipping areas (from north to south: Boston, New York City harbor, Delaware Bay, Chesapeake Bay, Charleston, and Savannah). Although petrels were exposed to shipping activity in most of their core areas and home range (range: 73.81-94.64 % of overlap; Table 3), major shipping routes generally did not overlap with petrel distribution, except for the Chesapeake-Charleston segment in the area of Cape Hatteras. On average in core areas, the dark form was largely more exposed than the light form to cargo ships, tankers, and other types of vessels (Table 3 and Figure S10). On average in home ranges, the dark form was moderately more exposed than the light form to cargo ships, tankers, and other types of vessels (Table 3 and Figure S10).

One Black-capped Petrel overlapped with active hydrocarbon exploration areas 2435 (n = 3 locations) and 2436 (n=5 locations), in Nova Scotia, Canada (Table S3 and Figure 2). No individuals overlapped with active leases for wind energy. The nearest active wind energy lease area to a petrel location was in North Carolina (29.5 km from location). Five individuals overlapped with proposed lease area E, and six with proposed lease area F, for wind energy production in the Central Atlantic DRAFT Call for Information and Nomination Area (https://www.regulations.gov/document/BOEM-2022-0023-0001; accessed 1 May 2022). Three individuals were present in both leases. The home range of the dark form slightly overlapped with the active wind lease area in North Carolina (138.5 km^2^, 0.05 % of the home range). Drafted Central Atlantic wind areas overlapped with core areas (dark: 4.5% of core area; light: 0.9%) and home ranges (dark: 3.4% of home range; light: 2.7%) of both forms (Figure 2 and Table S4). 95-100% of the drafted Central Atlantic wind areas E and F were in the home range of both forms, and 90% of area F was in the core area of the dark form (Table S4 and Figure S11).

## DISCUSSION

We used tracks from 7 Black-capped Petrels captured at sea to characterize the spatial distribution of the species in the western North Atlantic. We provide an updated assessment of marine areas used by dark and light forms and new information from two individuals on the apparent allochrony of color forms at sympatric breeding sites. Contrary to our hypotheses, color forms shared similar breeding areas in the Caribbean but had distinct non-breeding distributions, and thus may experience phenotypically specific exposure to marine threats.

### Marine distribution

More than 5,500 at-sea observations of Black-capped Petrels have been confirmed in the U.S. EEZ since 1938 (Sussman and U.S. Geological Survey 2014, Jodice et al. 2021). Together with data from the Caribbean Sea, these records have been used to infer the global range and distribution of the species (Simons et al. 2013, Winship et al. 2018, Leopold et al. 2019, Jodice et al. 2021)(Figure S12). At-sea surveys are limited, however, by season, weather, time of day, sea state, and bias due to research funding. Additionally, surveys often do not distinguish individual characteristics such as phenotype, sex, or breeding status, and cannot provide information on connectivity among oceanic basins, and between breeding sites and marine areas. Unlike systematic surveys, opportunistic pelagic trips, which specifically look for and attract seabirds at a short distance, tend to collect information on phenotypes but are often geographically limited to foraging hotspots that are easily accessible in a single day.

Given adequate sampling, individual-based tracking can provide detailed information on movements and marine usage at the level of the individual and the population but is restricted by the logistics and difficulties of capturing target individuals, ethical considerations of animal research, and the costs of tracking technology. In the Black-capped Petrel, tracking data are limited to the results of two studies, each with a sample size of 3 tracked individuals (Jodice et al. 2015, Satgé et al. 2019). Covering a much larger spatial extent than these previous efforts, our study provides a substantial increase in data available to assess the marine distribution of the species. Our dataset showed high overlap with data from at-sea surveys in and around the Hatteras hotspot (i.e. an area similar to the core area of the dark form in our study). Both systematic and opportunistic at-sea survey data, however, highlighted an area absent from use in our study, the Charleston Bump/Hoyt Hills (Figure 1). This area, located ∼100 km southeast of Charleston, South Carolina and to the northwest of the Blake Spur, has been a historical hotspot for Black-capped Petrel observations (Haney 1987, eBird 2021) but petrels in our study and in Jodice et al. (2015) showed very limited use of this area. In contrast, tracking data showed a much wider use of pelagic waters off the Middle Atlantic Bight and most of the Virginian marine ecoregion compared to at-sea surveys. Most of the population is strongly concentrated in Gulf Stream waters in the Carolinian ecoregion, but petrels in the Virginian region utilized an area of pelagic waters between the northern wall of the Gulf Stream and the continental slope. Finally, except in very limited locations within ca. 100 km of Cape Hatteras, tracked Black-capped Petrels never occurred on the continental shelf. This is consistent with at-sea surveys in which, except in two areas around Cape Hatteras and the Charleston Bump, petrels were only recorded on the continental slope and rise.

By specifically targeting individuals of distinct phenotypes, we also were able to test our hypothesis that both phenotypes had similar non-breeding distributions, and to gain insight on the connectivity with breeding sites. Observations from opportunistic surveys suggest that dark form individuals primarily use central/Carolinian waters and light form individuals primarily use northern/Virginian waters (eBird 2021), but these records are insufficient for range determination due to their anecdotal nature and inconsistent methodology. Nevertheless, our dataset supports this suggestion by showing the same spatial separation in the distributions of the two forms. The dark form occupies pelagic Carolinian waters of the South Atlantic Bight and is concentrated within a ∼ 200-km strip of waters extending eastward from the continental shelf into the Gulf Stream; the light form occupies pelagic Virginian waters of the Middle Atlantic Bight, extending over a wider area between the continental shelf and the northern edge of the Gulf Stream. Consequently, the dark form frequented warm surface waters of the Gulf Stream and, in fact, appears to be one of the seabirds in the western North Atlantic most strongly associated with the Gulf Stream (Winship et al. 2018). In contrast, the light form utilized an area of significantly deeper and colder waters showing a wider range of temperatures and more complex oceanic processes (Lai and Richardson 1977, Kang and Curchitser 2013). Despite this geographic separation between the two forms, light-form petrels remain influenced by the Gulf Stream, whether by latitudinal variability in the path of the Gulf Stream itself or through anticyclonic eddies that diffuse northward (Kang and Curchitser 2013, Seidov et al. 2019). Therefore, the difference in sea surface temperature that we observed between phenotypes may be more an indication of macroscale differences in habitat availability than active selection of temperate waters by light-form petrels. Indeed, within an overall area of temperate oceanic waters, it is likely that light-form petrels select areas directly influenced by warm oligotrophic waters of anticyclonic eddies (Kang and Curchitser 2013). This selection of eddies has been observed in several species of subtropical and tropical seabirds (e.g. Haney 1986, Hyrenbach et al. 2006, Tew Kai and Marsac 2010). Generally, anticyclonic eddies tend to capture material drifting at the surface (including plankton and fish larvae) and cyclonic eddies tend to expel it to peripheral fronts, hence both potentially increasing prey availability to seabirds at the mesoscale (Cotté et al. 2007). Thus, the more pelagic distribution of the light form may be a response to the lower predictability of anticyclonic eddies (Tew Kai and Marsac 2010) rather than to Gulf Stream fronts, and to a patchier geographic availability of prey across the area (Weimerskirch 2007, Clay et al. 2017).

### Influence of breeding connectivity, phenology, and distribution on genetic structure

Two PTTs, deployed on a male petrel of each phenotype (hereafter, light and dark individual), were still transmitting when the petrels travelled to suspected breeding areas. In late September 2019, the light individual travelled to eastern Hispaniola. Location data and tag performance suggest this petrel was active at a breeding site (possibly Valle Nuevo in the Cordillera Central of the Dominican Republic) during 02 – 08 and 09 – 15 October 2019. Valle Nuevo was confirmed as a nesting area in 2018, when three nests of the dark form were discovered. In October 2019, the dark individual travelled to the La Visite breeding area of eastern Haiti. Location data and tag performance suggest this petrel was active at this breeding site between 09 – 22 November, and 29 November – 03 December 2019. Connectivity between the breeding area of Loma del Toro, at the border between the Dominican Republic and Haiti, and Gulf Stream waters of the western North Atlantic was revealed by Jodice et al. (2015) but our study is the first to explicitly link the marine and terrestrial areas of a light-form Black-capped Petrel, and to show that this phenotype is likely to breed in sympatry with the dark form. Our study also confirms for the first time the link between Gulf Stream waters and the breeding area of La Visite. Moreover, our small dataset supports suggestions based on molt and phenology in the Gulf Stream that light-form petrels may be breeding 1 to 1.5 months earlier than the dark form (Howell and Patteson 2008, Manly et al. 2013).

Our results also suggest that the dual marine distribution of dark- and light-forms may be related more to phenotype than date. All individuals were captured within a 25 km radius within six days in May 2019. Following captures, most light-form petrels travelled north and stayed in the Virginian ecoregion until the fall, except for one individual that remained in the Carolinian region until June, at which point data transmission ceased. Dark-form petrels remained in the Carolinian region until late-July/early-August, when they moved to the Virginian region for ca. one month (n = 3). Information is lacking for the remainder of the annual cycle but the visits to breeding sites in the Caribbean at the end of the tracking period suggest that some degree of tracking nevertheless occurred for most of the non-breeding period. Therefore, although limited by a small sample size, our analysis confirms that phenotypes of the Black-capped Petrels have distinct non-breeding distributions, independently of the time of the year.

In vagile species, genetic differentiation may be a consequence of distribution (Friesen 2015, Wiley et al. 2012), phenology (Friesen et al. 2007b), dispersal (Burridge and Waters 2020), or ecological niche (Ryan et al. 2014). Our tracking data suggest that petrels experience temporal but not spatial reproductive isolation at breeding sites on Hispaniola. The lack of spatial segregation is also supported by monitoring data collected after our study which confirmed that nests of both phenotypes occurred within an area of 0.14 km^2^ at Valle Nuevo (Wheeler et al. 2021). Allochrony in sympatry exists in seabirds over time frames of both several months (e.g., Band-Rumped Storm-Petrels *Oceanodroma sp*. that breed in a cool or warm season: Friesen et al 2007b, Sato et al. 2010, Krüger et al. 2016, Wallace et al. 2017) and on the order of several weeks (Tomkins and Milne 1991, Brooke and Rowe 1996, Brown et al. 2015).

The spatial segregation we observed in marine distributions of Black-capped Petrel phenotypes may also contribute to non-breeding isolation. Although our limited dataset prevents us from inferring whether individuals in our study were from sympatric or distinct breeding areas, segregation in marine distributions is common within species with allopatric breeding (Rayner et al. 2011, Priddel et al. 2014, Clay et al. 2016, Hipfner et al. 2020) and between species breeding in sympatry (Friesen 2007a, McFarlane et al. 2015, Jones et al. 2020). Furthermore, differences in morphometrics between dark and light forms (Howell and Patteson 2008, this study) suggest that niche partitioning may also be occurring (Ryan et al. 2014, Rayner et al. 2016, Miller et al. 2018). For example, differences in foraging strategies in response to use of different oceanic habitats and distinct availability of prey may also influence spatial segregation (Wiley et al. 2012, Navarro et al. 2015, Pontón-Cevallos et al. 2017, Jones et al. 2020). Therefore, it appears that multiple mechanisms may be operating that could contribute to genetic differentiation in this species, although the causes of their occurrence and the importance of each mechanism remain untested and warrant additional investigation.

### Representativeness of captured individuals

Among pelagic seabirds, differences in the distribution of unsuccessful, successful, or non-breeding adults may occur and therefore could contribute to the dual distribution we observed in our study (e.g., a confounding effect of breeding status). For example, failed breeders may have similar or different distributions compared to non-breeders (Phillips et al. 2005, Clay et al. 2016), failed breeders may disperse farther than successful breeders (Bogdanova et al. 2011), or failed breeders may remain resident within the breeding range or migrate but return earlier to breeding sites than successful breeders (Catry et al. 2013). The reproductive status of petrels tracked in this study was not known at the time of capture and therefore we are unable to accurately assess the potential impact of reproductive status on distribution. For example, petrels utilizing waters off Cape Hatteras in May could be either active breeders (Jodice et al. 2015, Satgé et al. 2019), failed breeders, non-breeders (including skip-breeders, i.e. mature petrels who bred successfully the previous year and skipped breeding this current year: Hunter et al. 2000, Dillingham and Fletcher 2011, Taylor et al. 2011), or immatures (< 4 years of age, Simons et al. 2013). Except for the two individuals that were tracked to Hispaniola (which were likely adults of breeding age), we cannot tell if the petrels were immatures or adults. Nonetheless, we suggest that the dark individuals we captured in May 2019 were unlikely to be breeding adults. For example, juveniles of dark-form petrels fledge after mid-June (Simons et al. 2013; E. Rupp, Grupo Jaragua Inc., oral communication, 2021) suggesting foraging ranges of breeders are more likely to be restricted during May. Additionally, Jodice et al. (2015) showed that chick-rearing petrels made repeated foraging trips within the Caribbean Sea and were less likely to occupy Gulf Stream waters, while failed breeders vacated the Caribbean and travelled to Gulf Stream waters. These data would support dark form birds occurring readily in Gulf Stream waters in May to be non-breeders or failed breeders. The breeding status of light form birds is more challenging to predict due to limited data from this phenotype. Preliminary data gathered from camera traps from nests of light-form petrels in Valle Nuevo suggest, however, that fledging occurs in late April and hence our captures may have been post-breeders (E. Rupp, Grupo Jaragua Inc., oral communication, 2021). These data, along with the limited extent of the species distribution in the western North Atlantic and the fact that the dual phenotypic distribution is supported by at-sea surveys, suggest that differences in the distribution of unsuccessful, successful, or non-breeders may be limited in scale in Black-capped Petrels and that the dual distribution we observed may be driven primarily by phenotype.

### Marine threats

Conservation planning benefits from current assessments of vulnerability of species to extrinsic factors such as anthropogenic threats (Butt et al. 2016). Although trait-based approaches are preferred for threat assessments (Butt and Gallagher 2018, Zhou et al. 2019, Richards et al. 2021), substantial data gaps on the biology and ecology of Black-capped Petrels and uncertainties about the effect of threats, notably marine ones (Wheeler et al. 2021), limit their applicability. Nonetheless, a review of marine threats and likely exposure based on spatial distribution of the species in the western North Atlantic is warranted (Wheeler et al. 2021). Here, we assess exposure at the macro-scale *sensu* Burger et al. (2011) and Waggit and Scott (2014), i.e. the occurrence of the species of concern within the geographical area of interest (e.g., the broad area where the threat occurs and overlaps with the species). We assume here that spatial overlap provides an adequate proxy to assess exposure, with the caveat that overlap at the macroscale likely over-estimates actual overlap at the meso- and microscale (i.e. at finer spatial and temporal scales, Torres et al. 2013). Therefore, our metrics provide an initial estimate of potential risk for dark and light forms of the species.

### Marine plastic

Marine plastic is a global and increasing issue for seabirds (Dias et al. 2019). The Black-capped Petrel is expected to be susceptible because of its foraging behavior and its limited capacity to regurgitate plastic fragments (Furness 1985, Moser and Lee 1992, Rodríguez et al. 2019). Currently, our understanding of plastic exposure in the Black-capped Petrel is limited to a single study (Moser and Lee 1992) that showed the frequency of occurrence of plastics in the digestive tract of 57 individual petrels was 1.8%, well below other procellariforms in the study. Our data indicate that macro-scale exposure to plastics was higher in the dark form compared to the light form. For example, use areas of the dark form overlapped with the waters influenced by the southwestern region of the North Atlantic Subtropical Gyre where the Gulf Stream and Antilles Current converge and where micro-plastics accumulate (Law et al. 2010, Enders et al. 2015, van Sebille et al. 2015). In contrast, in the more northern areas utilized by the light form, plastic accumulation was limited, due to limited ingress from less-populated areas in northern North America, and oceanic and atmospheric processes limiting accumulation in the upper oceanic layer (Law et al. 2010). Nevertheless, research suggests that anticyclonic eddies concentrate micro-plastics (Brach et al. 2018). Since the majority of anticyclonic eddies originating from the Gulf Stream are located in the northern areas of the western North Atlantic Ocean, light-form Black-capped Petrels may be locally more exposed than dark-form individuals at the meso- and micro-scale.

### Fisheries

Because of their foraging behavior, *Pterodroma* petrels, including the Black-capped Petrel, are generally considered less susceptible (although not immune) to bycatch than larger pelagic species (Waugh et al. 2012, Simons et al. 2013; but see Trebilco et al. 2010, Richard et al. 2017). Nonetheless, studies assessing overlap between fisheries and the distribution of *Pterodroma* petrels are lacking (Bugoni et al. 2008, Ramos et al. 2017). Black-capped Petrels have not been historically recorded as bycatch in the western North Atlantic (Li et al. 2006), but Zhou et al. (2019) predict the species may be at risk of bycatch in the pelagic longline fishery in the region. Moreover, foraging by Black-capped Petrels on chum and discards of offal may expose the species to collisions with cables used in trawl fisheries (including net-sonde cables which are used in the region), a type of bycatch that is often overlooked by on-board observers (González-Zevallos et al. 2007, Adasme et al. 2019). Our macro-scale assessment shows limited spatial overlap (4-16 % of core areas and 12-14 % of home ranges) but relatively high levels of fishing effort (9.5-14 h/cell in core areas and 15.5-51 h/cell in home ranges), although, in our study area, pelagic waters are subject to relatively low monthly fishing effort compared to global effort (Guiet et al. 2019). Within our study area, four fisheries were responsible for most of the fishing effort in Black-capped Petrel use areas: demersal trawl, bottom-set longlines, pelagic longlines, and line fishing (Guiet et al. 2019, Global Fishing Watch 2021). Drifting gear dominated in the South Atlantic Bight, fixed gear dominated off Cape Hatteras, and no gear type was clearly dominant throughout the Middle Atlantic Bight (Guiet et al. 2019). Trawling was responsible for the higher fishing effort observed in the home range of the light phenotype but overlap was restricted to ∼ 0.3 % of the home range area, along the oceanic edge of Georges Bank. Within petrel use areas, bottom-set longlines were mostly concentrated along the continental slope of the Middle Atlantic Bight and Georges Bank. The highest effort for pelagic longlines was concentrated off Cape Hatteras, with localized hotspots along Georges Bank, and a large area (ca. 20,000 km^2^) of moderate effort around the Charleston Bump. Finally, pole and line effort was localized along the continental edge of Georges Bank and the Nantucket Shoals. Although the relationship between overlap and bycatch is complex and requires information that is currently missing for the Black-capped Petrel (Wheeler et al. 2021), our results suggest that overlap does exist between endangered Black-capped Petrels and fisheries in the U.S. EEZ in the western North Atlantic, and that data collected at a finer spatial and temporal scale may benefit conservation assessments for the species.

### Marine traffic

In the western North Atlantic, marine traffic includes shipping (mainly cargo and tanker) and fishing vessels (Wu et al. 2017). These can have adverse effects on seabirds through attraction and collision with lighted vessels (Glass and Ryan 2013, Ryan et al. 2021), pollution (Heubeck et al. 2003, Fox et al. 2016, Lieske et al. 2020, King et al. 2021), and displacement (Lieske et al. 2020), leading to lethal or sublethal effects (Matcott et al. 2019, King et al. 2021). Filtered satellite imagery shows that some level of marine lighting is present in most of the petrel’s use areas (VIIRS boat detections, accessed through Global Fishing Watch https://globalfishingwatch.org/map/) during the non-breeding period. Moreover, most, if not all, of the use areas of Black-capped Petrels are impacted by some level of marine traffic (between 80-100 % of use areas), mostly from cargo ships and other vessels (including fishing vessels). The area of highest overlap was in neritic waters off Cape Hatteras. This area is a hotspot of Black-capped Petrel activity within the core area of the dark form. Our analysis highlights exposure to marine traffic in the U.S. EEZ, but our results are limited by the availability of open access marine traffic data in the Canadian EEZ (including shipping channels to and from Nova Scotia) and the high seas. Our results should therefore be considered a conservative assessment of Black-capped Petrel exposure to marine traffic in the western North Atlantic.

### Marine energy

Marine energy activities in the western North Atlantic include petroleum exploration and extraction, and production of offshore wind energy (Goodale et al. 2019). Petrels are susceptible to point-based attraction to continuous lighting of staffed oil and gas production platforms, and irregular lighting from support vessels and gas flaring, which can cause disorientation, grounding, and direct mortality (Ronconi et al. 2015, Fraser and Carter 2018). Petrels may also be exposed to accidental oil spills and regular discharge of produced waters, through direct (contact with contaminated waters) or indirect (through bioaccumulation of contaminants in the food chain) entryways (Fraser et al. 2006, Ronconi et al. 2015, Jodice et al. 2021). There is currently no active petroleum production in areas used by Black-capped Petrels in our study but active exploratory leases are present in the region and one individual in our study was present in leases 2435 and 2436. Additionally, a moratorium on petroleum extraction in the Canadian portion of Georges Bank is set to expire in 2022 (Georges Bank Protection Act (S.C. 2015, c. 39), https://laws-lois.justice.gc.ca/eng/annualstatutes/2015_39/page-1.html; accessed 1 November 2021). On the U.S. Atlantic coast, there are currently no active oil and gas leases (https://www.boem.gov/oil-gas-energy/oil-and-gas-atlantic; accessed 1 November 2021) but several leases for the production of offshore wind energy are in active states of development in the Middle Atlantic Bight (https://www.boem.gov/renewable-energy/state-activities; accessed 1 May 2022). Because wind energy production is currently logistically constrained to neritic waters, Black-capped Petrels in our study were not present within active leases. The closest location of a Black-capped Petrel to an active lease area was ca. 30-km from lease OCS-A 0508 in waters offshore of North Carolina. In early 2022, draft planning areas have been proposed on the outer continental shelf and rise of the Central Atlantic coast of the U.S. (Figure 1). These areas overlapped with home ranges and core areas of both phenotypes (Figures 2 and S11). Therefore, if future marine energy exploration and production were to occur in these pelagic waters of the western North Atlantic, our data suggest that Black-capped Petrels be considered for inclusion in ecological assessments.

## CONCLUSIONS

Our study highlights the connectivity between foraging areas of Black-capped Petrels in the western North Atlantic and their nesting sites on Hispaniola. Furthermore, our dataset demonstrates that dark and light phenotypes of the Black-capped Petrel differed in the spatial extent of waters they occupied within the western North Atlantic. The two phenotypes may be using different foraging habitats, although the small sample size and coarse scale of our dataset prevented a detailed modeling of habitat use at this time. We also demonstrated that differences in phenotype were linked to differences in breeding phenology. Allochrony may be an initial driver of speciation but may also contribute to divergence at any stage along a speciation process (Taylor and Friesen 2017). Conserving these evolutionary processes would be more likely to sustain biodiversity that responds adaptively to environmental changes (Ennos et al. 2005). In addition, differences in morphology between phenotypes suggest that dark and light petrels may select different prey, using different foraging strategies, which raises several new questions on the ecological niche and niche partitioning of Black-capped Petrels at sea. Further research could focus on detailed analyses of habitat selection, and shed light on the causes and consequences of this dual distribution between phenotypes. Additional tracking of Black-capped Petrels from known breeding areas not yet studied (e.g. Valle Nuevo, Dominican Republic, or La Visite, Haiti) could supplement our data and thus provide a more complete assessment of use areas and marine threats (Wheeler et al. 2021). As new breeding areas are discovered, individual-based tracking may be considered to further assess the global connectivity of the species and identify areas of exposure to anthropic stressors.

## ACKNOWLEDGEMENTS

This study would not have been possible without Seabirding Pelagic Trips, Hatteras, North Carolina, and the trustworthy Stormy Petrel II. We thank Kate Sutherland for help throughout the study. We are grateful to Autumn-Lynn Harrison, of the Smithsonian Institute’s Migratory Connectivity Project, for generously donating two satellite transmitters. We also thank Ari Friedlaender, of University of California Santa Cruz, for lending us the whale-tagger, and Dive Hatteras in Frisco, North Carolina for generously providing air tanks. Monica Silva, of Universidade de Lisboa, Portugal performed the molecular sexing. Zhang Yanxu, of Nanjing University, shared the mercury raster; Erik van Sebille, of Universiteit Utrecht, shared micro-plastic datasets; and Carina Gjerdrum, of Environment and Climate Change Canada, provided shapefiles of Oil and Gas leases in Canada. Teresa Militão and Andrew Read provided helpful reviews of the manuscript prior to submission. Funding for this research was provided by the bin Zayed Species Conservation Fund, American Bird Conservancy, the South Carolina Cooperative Fish and Wildlife Research Unit, and private donors who contributed to American Bird Conservancy’s fundraising campaign. The South Carolina Cooperative Fish and Wildlife Research Unit is jointly supported by the US Geological Survey, South Carolina Department of Natural Resources, and Clemson University. Any use of trade, firm, or product names is for descriptive purposes only and does not imply endorsement by the U.S. Government.

## REFERENCES

Adasme, L. M., Canales, C. M., & Adasme, N. A. (2019). Incidental seabird mortality and discarded catches from trawling off far southern Chile (39–57° S). ICES Journal of Marine Science, 76(4), 848–858.

Alerstam, T., Bäckman, J., Grönroos, J., Olofsson, P., & Strandberg, R. (2019). Hypotheses and tracking results about the longest migration: The case of the arctic tern. Ecology and Evolution, 9(17), 9511–9531.

Amante, C., & Eakins, B. W. (2009). ETOPO1 arc-minute global relief model: procedures, data sources and analysis.

Arponen, A. (2012). Prioritizing species for conservation planning. Biodiversity and Conservation, 21(4), 875–893.

Bernard, A., Rodrigues, A. S., Cazalis, V., & Grémillet, D. (2021). Toward a global strategy for seabird tracking. Conservation Letters, 14(3), e12804.

BirdLife International (2018). Pterodroma hasitata. The IUCN Red List of Threatened Species 2018: e.T22698092A132624510. https://dx.doi.org/10.2305/IUCN.UK.2018-2.RLTS.T22698092A132624510.en. Accessed on 01 July 2021.

Bogdanova, M. I., Daunt, F., Newell, M., Phillips, R. A., Harris, M. P., & Wanless, S. (2011). Seasonal interactions in the black-legged kittiwake, Rissa tridactyla: links between breeding performance and winter distribution. Proceedings of the Royal Society B: Biological Sciences, 278(1717), 2412–2418.

Bolton, M., Smith, A. L., GÓMEZ-DÍAZ, E. L. E. N. A., Friesen, V. L., Medeiros, R., Bried, J., … & Furness, R. W. (2008). Monteiro’s Storm-petrel Oceanodroma monteiroi: a new species from the Azores. Ibis, 150(4), 717–727.

Brach, L., Deixonne, P., Bernard, M. F., Durand, E., Desjean, M. C., Perez, E., … & Ter Halle, A. (2018). Anticyclonic eddies increase accumulation of microplastic in the North Atlantic subtropical gyre. Marine Pollution Bulletin, 126, 191–196.

Bridle, J. R., Pedro, P. M., & Butlin, R. K. (2004). Habitat fragmentation and biodiversity: testing for the evolutionary effects of refugia. Evolution, 58(6), 1394–1396.

Brooke, M. D. L., & Rowe, G. (1996). Behavioural and molecular evidence for specific status of light and dark morphs of the Herald Petrel Pterodroma heraldica. Ibis, 138(3), 420–432.

Brown, R. M., Techow, N. M., Wood, A. G., & Phillips, R. A. (2015). Hybridization and back-crossing in giant petrels (Macronectes giganteus and M. halli) at Bird Island, South Georgia, and a summary of hybridization in seabirds. PLoS ONE, 10(3), e0121688.

Bugoni, L., Mancini, P. L., Monteiro, D. S., Nascimento, L., & Neves, T. S. (2008). Seabird bycatch in the Brazilian pelagic longline fishery and a review of capture rates in the southwestern Atlantic Ocean. Endangered Species Research, 5(2-3), 137–147.

Burger, J., Gordon, C., Lawrence, J., Newman, J., Forcey, G., & Vlietstra, L. (2011). Risk evaluation for federally listed (roseate tern, piping plover) or candidate (red knot) bird species in offshore waters: A first step for managing the potential impacts of wind facility development on the Atlantic Outer Continental Shelf. Renewable Energy, 36(1), 338–351.

Burridge, C. P., & Waters, J. M. (2020). Does migration promote or inhibit diversification? A case study involving the dominant radiation of temperate Southern Hemisphere freshwater fishes. Evolution, 74(9), 1954–1965.

Butt, N., & Gallagher, R. (2018). Using species traits to guide conservation actions under climate change. Climatic Change, 151(2), 317–332.

Butt, N., Possingham, H. P., De Los Rios, C., Maggini, R., Fuller, R. A., Maxwell, S. L., & Watson, J. E. M. (2016). Challenges in assessing the vulnerability of species to climate change to inform conservation actions. Biological Conservation, 199, 10–15.

Calenge, C. (2006). The package “adehabitat” for the R software: a tool for the analysis of space and habitat use by animals. Ecological Modeling, 197(3-4), 516–519.

Catry, P., Dias, M. P., Phillips, R. A., & Granadeiro, J. P. (2013). Carry-over effects from breeding modulate the annual cycle of a long-distance migrant: An experimental demonstration. Ecology, 94(6), 1230–1235.

Clay, T. A., Manica, A., Ryan, P. G., Silk, J. R., Croxall, J. P., Ireland, L., & Phillips, R. A. (2016). Proximate drivers of spatial segregation in non-breeding albatrosses. Scientific Reports, 6(1), 1–13.

Clay, T. A., Phillips, R. A., Manica, A., Jackson, H. A., & Brooke, M. D. L. (2017). Escaping the oligotrophic gyre? The year-round movements, foraging behaviour and habitat preferences of Murphy’s petrels. Marine Ecology Progress Series, 579, 139–155.

Cohen, J. (1988). Statistical power analysis for the behavioral. Hillsdale, NJ: Lawrence Erlbaum Associates Sciences.

Cotté, C., Park, Y. H., Guinet, C., & Bost, C. A. (2007). Movements of foraging king penguins through marine mesoscale eddies. Proceedings of the Royal Society B: Biological Sciences, 274(1624), 2385–2391.

Croxall, J. P., Silk, J. R., Phillips, R. A., Afanasyev, V., & Briggs, D. R. (2005). Global circumnavigations: tracking year-round ranges of nonbreeding albatrosses. Science, 307(5707), 249–250.

Cummings, J. A., & Smedstad, O. M. (2013). Variational data assimilation for the global ocean. In Data Assimilation for Atmospheric, Oceanic and Hydrologic Applications (Vol. II) (pp. 303–343). Springer, Berlin, Heidelberg.

Danckwerts, D. K., Humeau, L., Pinet, P., McQuaid, C. D., & Le Corre, M. (2021). Extreme philopatry and genetic diversification at unprecedented scales in a seabird. Scientific Reports, 11(1), 1–12.

Deiner, K., Garza, J. C., Coey, R., & Girman, D. J. (2007). Population structure and genetic diversity of trout (Oncorhynchus mykiss) above and below natural and man-made barriers in the Russian River, California. Conservation Genetics, 8(2), 437–454.

Dias, Maria P., Rob Martin, Elizabeth J. Pearmain, Ian J. Burfield, Cleo Small, Richard A. Phillips, Oliver Yates, Ben Lascelles, Pablo Garcia Borboroglu, and John P. Croxall. “Threats to seabirds: a global assessment.” Biological Conservation 237 (2019): 525–537.

Dillingham, P. W., & Fletcher, D. (2011). Potential biological removal of albatrosses and petrels with minimal demographic information. Biological Conservation, 144(6), 1885–1894.

eBird (2021). eBird: An online database of bird distribution and abundance. eBird, Cornell Lab of Ornithology, Ithaca, New York. Accessed from http://www.ebird.org on 01 November 2021.

Enders, K., Lenz, R., Stedmon, C. A., & Nielsen, T. G. (2015). Abundance, size and polymer composition of marine microplastics≥ 10 μm in the Atlantic Ocean and their modelled vertical distribution. Marine Pollution Bulletin, 100(1), 70–81.

Ennos, R. A., French, G. C., & Hollingsworth, P. M. (2005). Conserving taxonomic complexity. Trends in Ecology & Evolution, 20(4), 164–168.

Evers, D. C., Kaplan, J. D., Meyer, M. W., Reaman, P. S., Braselton, W. E., Major, A., … & Scheuhammer, A. M. (1998). Geographic trend in mercury measured in common loon feathers and blood. Environmental Toxicology and Chemistry: An International Journal, 17(2), 173–183.

Fieberg, J., & Kochanny, C. O. (2005). Quantifying home-range overlap: The importance of the utilization distribution. The Journal of Wildlife Management, 69(4), 1346–1359.

Fischer, J. H., Debski, I., Spitz, D. B., Taylor, G. A., & Wittmer, H. U. (2021). Year-round offshore distribution, behaviour, and overlap with commercial fisheries of a Critically Endangered small petrel. Marine Ecology Progress Series, 660, 171–187.

Fox, C. H., O’hara, P. D., Bertazzon, S., Morgan, K., Underwood, F. E., & Paquet, P. C. (2016). A preliminary spatial assessment of risk: Marine birds and chronic oil pollution on Canada’s Pacific coast. Science of the Total Environment, 573, 799–809.

Frankham, R., Ballou, S. E. J. D., Briscoe, D. A., & Ballou, J. D. (2002). Introduction to conservation genetics. Cambridge, UK: Cambridge university press.

Fraser, G. S., & Carter, A. V. (2018). Seabird attraction to artificial light in Newfoundland and Labrador’s offshore oil fields: documenting failed regulatory governance. Ocean Yearbook Online, 32(1), 265–282.

Fraser, G. S., Russell, J., & Von Zharen, W. M. (2006). Produced water from offshore oil and gas installations on the Grand Banks, Newfoundland: are the potential effects to seabirds sufficiently known?. Marine Ornithology, 34, 147–156.

Fridolfsson, A. K., & Ellegren, H. (1999). A simple and universal method for molecular sexing of non-ratite birds. Journal of avian biology, 116–121.

Friesen, V. L., Burg, T. M., & McCoy, K. D. (2007). Mechanisms of population differentiation in seabirds. Molecular Ecology, 16(9), 1765–1785.

Friesen, V. L., Smith, A. L., Gómez-Díaz, E., Bolton, M., Furness, R. W., González-Solís, J., & Monteiro, L. R. (2007). Sympatric speciation by allochrony in a seabird. Proceedings of the National Academy of Sciences, 104(47), 18589–18594.

Friesen, V. L. (2015). Speciation in seabirds: why are there so many species… and why aren’t there more?. Journal of Ornithology, 156(1), 27–39.

Furness, R. W. (1985). Ingestion of plastic particles by seabirds at Gough Island, South Atlantic Ocean. Environmental Pollution Series A, Ecological and Biological, 38(3), 261–272.

Gaston, A. J. (2001). Taxonomy and conservation: thoughts on the latest BirdLife International listings for seabirds. Marine Ornithology, 29, 1–6.

Glass, J. P., & Ryan, P. G. (2013). Reduced seabird night strikes and mortality in the Tristan rock lobster fishery. African Journal of Marine Science, 35(4), 589–592.

Global Fishing Watch (2021). Fishing effort. Accessed from https://globalfishingwatch.org/data-download/datasets/public-fishing-effort on 01 July 2021.

González-Zevallos, D., Yorio, P., & Caille, G. (2007). Seabird mortality at trawler warp cables and a proposed mitigation measure: A case of study in Golfo San Jorge, Patagonia, Argentina. Biological Conservation, 136(1), 108–116.

Goodale, M. W., Milman, A., & Griffin, C. R. (2019). Assessing the cumulative adverse effects of offshore wind energy development on seabird foraging guilds along the East Coast of the United States. Environmental Research Letters, 14(7), 074018.

Goutte, A., Bustamante, P., Barbraud, C., Delord, K., Weimerskirch, H., & Chastel, O. (2014). Demographic responses to mercury exposure in two closely related Antarctic top predators. Ecology, 95(4), 1075–1086.

Goutte, A., Kriloff, M., Weimerskirch, H., & Chastel, O. (2011). Why do some adult birds skip breeding? A hormonal investigation in a long-lived bird. Biology letters, 7(5), 790–792.

Guiet, J., Galbraith, E., Kroodsma, D., & Worm, B. (2019). Seasonal variability in global industrial fishing effort. PloS ONE, 14(5), e0216819.

Haney, J. C. (1986). Seabird segregation at Gulf Stream frontal eddies. Marine Ecology Progress Series, 28(3), 279–285.

Haney, J. C. (1987). Aspects of the pelagic ecology and behavior of the black-capped petrel (Pterodroma hasitata). Wilson Bulletin, 99(2), 153–312.

Hays, G. C., Bailey, H., Bograd, S. J., Bowen, W. D., Campagna, C., Carmichael, R. H., … & Sequeira, A. M. (2019). Translating marine animal tracking data into conservation policy and management. Trends in Ecology & Evolution, 34(5), 459–473.

Hellberg, M. E. (2009). Gene flow and isolation among populations of marine animals. Annual Review of Ecology, Evolution, and Systematics, 40, 291–310.

Heubeck, M., Camphuysen, K. C., Bao, R., Humple, D., Rey, A. S., Cadiou, B., … & Thomas, T. (2003). Assessing the impact of major oil spills on seabird populations. Marine Pollution Bulletin, 46(7), 900–902.

Hijmans, R.J. (2019). geosphere: Spherical Trigonometry. R package version 1.5-10. https://CRAN.R-project.org/package=geosphere

Hipfner, J. M., Prill, M. M., Studholme, K. R., Domalik, A. D., Tucker, S., Jardine, C.,. & Burg, T. M. (2020). Geolocator tagging links distributions in the non-breeding season to population genetic structure in a sentinel North Pacific seabird. PloS ONE, 15(11), e0240056.

Howell, S. N. G., & Patteson, J. B. (2008). Variation in the Black-capped Petrel–one species or more. Alula, 14, 70–83.

Hunter, C. M., Moller, H., & Fletcher, D. (2000). Parameter uncertainty and elasticity analyses of a population model: setting research priorities for shearwaters. Ecological Modelling, 134(2-3), 299–324.

Hussey, N. E., Kessel, S. T., Aarestrup, K., Cooke, S. J., Cowley, P. D., Fisk, A. T., … & Whoriskey, F. G. (2015). Aquatic animal telemetry: a panoramic window into the underwater world. Science, 348(6240), 1255642.

Hyrenbach, K. D., Veit, R. R., Weimerskirch, H., & Hunt Jr, G. L. (2006). Seabird associations with mesoscale eddies: the subtropical Indian Ocean. Marine Ecology Progress Series, 324, 271–279.

IUCN Species Survival Commission. Species Conservation Planning Task Force. (2008). Strategic planning for species conservation: a handbook, version 1.0. IUCN.

Jodice, P. G., Ronconi, R. A., Rupp, E., Wallace, G. E., & Satgé, Y. (2015). First satellite tracks of the endangered black-capped petrel. Endangered Species Research, 29(1), 23–33.

Jodice, P. G., Michael, P. E., Gleason, J. S., Haney, J. C., & Satgé, Y. G. (2021). Revising the marine range of the endangered black-capped petrel Pterodroma hasitata: occurrence in the northern Gulf of Mexico and exposure to conservation threats. Endangered Species Research, 46, 49–65.

Jones, C. W., Phillips, R. A., Grecian, W. J., & Ryan, P. G. (2020). Ecological segregation of two superabundant, morphologically similar, sister seabird taxa breeding in sympatry. Marine Biology, 167(4), 1–16.

Jonsen, I. D., McMahon, C. R., Patterson, T. A., Auger-Méthé, M., Harcourt, R., Hindell, M. A., & Bestley, S. (2019). Movement responses to environment: fast inference of variation among southern elephant seals with a mixed effects model. Ecology, 100(1):e02566

Kanai, Y., Nagendran, M., Ueta, M., Markin, Y., Rinne, J., Sorokin, A. G., … & Archibald, G. W. (2002). Discovery of breeding grounds of a Siberian Crane Grus leucogeranus flock that winters in Iran, via satellite telemetry. Bird Conservation International, 12(4), 327–333.

Kang, D., & Curchitser, E. N. (2013). Gulf Stream eddy characteristics in a high-resolution ocean model. Journal of Geophysical Research: Oceans, 118(9), 4474–4487.

Kays, R., Crofoot, M. C., Jetz, W., & Wikelski, M. (2015). Terrestrial animal tracking as an eye on life and planet. Science, 348(6240), aaa2478.

King, M. D., Elliott, J. E., & Williams, T. D. (2021). Effects of petroleum exposure on birds: A review. Science of The Total Environment, 755, 142834.

Kroodsma, D. A., Mayorga, J., Hochberg, T., Miller, N. A., Boerder, K., Ferretti, F., … & Worm, B. (2018). Tracking the global footprint of fisheries. Science, 359(6378), 904–908.

Krüger, L., Paiva, V. H., Colabuono, F. I., Petry, M. V., Montone, R. C., & Ramos, J. A. (2016). Year-round spatial movements and trophic ecology of Trindade Petrels (Pterodroma arminjoniana). Journal of Field Ornithology, 87(4), 404–416.

Kuhn, M. (2020). caret: Classification and Regression Training. R package version 6.0-86. https://CRAN.R-project.org/package=caret

Lai, D. Y., & Richardson, P. L. (1977). Distribution and movement of Gulf Stream rings. Journal of Physical Oceanography, 7(5), 670–683.

Law, K. L., Morét-Ferguson, S., Maximenko, N. A., Proskurowski, G., Peacock, E. E., Hafner, J., & Reddy, C. M. (2010). Plastic accumulation in the North Atlantic subtropical gyre. Science, 329(5996), 1185–1188.

Le Bot, T., Lescroël, A., & Grémillet, D. (2018). A toolkit to study seabird–fishery interactions. ICES Journal of Marine Science, 75(5), 1513–1525.

Leopold, M. F., Geelhoed, S. C., Scheidat, M., Cremer, J., Debrot, A. O., & Van Halewijn, R. (2019). A review of records of the Black-capped Petrel Pterodroma hasitata in the Caribbean Sea. Marine Ornithology, 47, 235–241.

Li, Y., Jiao, Y., & Browder, J. A. (2016). Assessment of seabird bycatch in the US Atlantic pelagic longline fishery, with an extra exploration on modeling spatial variation. ICES Journal of Marine Science, 73(10), 2687–2694.

Lieske, D. J., Tranquilla, L. M., Ronconi, R. A., & Abbott, S. (2020). “Seas of risk”: Assessing the threats to colonial-nesting seabirds in Eastern Canada. Marine Policy, 115, 103863.

Lombal, A. J., O’dwyer, J. E., Friesen, V., Woehler, E. J., & Burridge, C. P. (2020). Identifying mechanisms of genetic differentiation among populations in vagile species: historical factors dominate genetic differentiation in seabirds. Biological Reviews, 95(3), 625–651.

Lombal, A. J., Wenner, T. J., Lavers, J. L., Austin, J. J., Woehler, E. J., Hutton, I., & Burridge, C. P. (2018). Genetic divergence between colonies of Flesh-footed Shearwater Ardenna carneipes exhibiting different foraging strategies. Conservation Genetics, 19(1), 27–41.

Mancilla-Morales, M. D., Romero-Fernández, S., Contreras-Rodríguez, A., Flores-Martínez, J. J., Sánchez-Cordero, V., Herrera M L. G., … & Ruiz, E. A. (2020). Diverging Genetic Structure of Coexisting Populations of the Black Storm-Petrel and the Least Storm-Petrel in the Gulf of California. Tropical Conservation Science, 13, 1940082920949177.

Manly, B., Arbogast, B. S., Lee, D. S., & Van Tuinen, M. (2013). Mitochondrial DNA analysis reveals substantial population structure within the endangered Black-capped Petrel (Pterodroma hasitata). Waterbirds, 36(2), 228–233.

Matcott, J., Baylis, S., & Clarke, R. H. (2019). The influence of petroleum oil films on the feather structure of tropical and temperate seabird species. Marine Pollution Bulletin, 138, 135–144.

McFarlane Tranquilla, L., Montevecchi, W. A., Hedd, A., Regular, P. M., Robertson, G. J., Fifield, D. A., & Devillers, R. (2015). Ecological segregation among Thick-billed Murres (Uria lomvia) and Common Murres (Uria aalge) in the Northwest Atlantic persists through the nonbreeding season. Canadian Journal of Zoology, 93(6), 447–460.

Miller, M. G., Silva, F. R., Machovsky-Capuska, G. E., & Congdon, B. C. (2018). Sexual segregation in tropical seabirds: drivers of sex-specific foraging in the brown booby Sula leucogaster. Journal of Ornithology, 159(2), 425–437.

Monteiro, L. R., Granadeiro, J. P., & Furness, R. W. (1998). Relationship between mercury levels and diet in Azores seabirds. Marine Ecology Progress Series, 166, 259–265.

Moser, M. L., & Lee, D. S. (1992). A fourteen-year survey of plastic ingestion by western North Atlantic seabirds. Colonial Waterbirds, 83–94.

Mott, R., & Clarke, R. H. (2018). Systematic review of geographic biases in the collection of at-sea distribution data for seabirds. Emu-Austral Ornithology, 118(3), 235–246.

Navarro, J., Cardador, L., Brown, R., & Phillips, R. A. (2015). Spatial distribution and ecological niches of non-breeding planktivorous petrels. Scientific reports, 5(1), 1–5.

Office for Coastal Management (2021). 2017 Vessel Transit Counts. Accessed from https://www.fisheries.noaa.gov/inport/item/55365 on 01 July 2021.

Pereira, J. M., Ramos, J. A., Marques, A. M., Ceia, F. R., Krüger, L., Votier, S. C., & Paiva, V. H. (2021). Low spatial overlap between foraging shearwaters during the breeding season and industrial fisheries off the west coast of Portugal. Marine Ecology Progress Series, 657, 209–221.

Phillips, R. A., Silk, J. R., Croxall, J. P., Afanasyev, V., & Bennett, V. J. (2005). Summer distribution and migration of nonbreeding albatrosses: individual consistencies and implications for conservation. Ecology, 86(9), 2386–2396.

Pontón-Cevallos, J., Dwyer, R. G., Franklin, C. E., & Bunce, A. (2017). Understanding resource partitioning in sympatric seabirds living in tropical marine environments. Emu-Austral Ornithology, 117(1), 31–39.

Pörtner, H. O., Scholes, R. J., Agard, J., Archer, E., Arneth, A., Bai, X., … & Ngo, H. T. (2021). IPBES-IPCC co-sponsored workshop report on biodiversity and climate change; IPBES and IPCC. Intergovernmental Science-Policy Platform on Biodiversity and Ecosystem Services (IPBES), 24.

Priddel, D., Carlile, N., Portelli, D., Kim, Y., O’Neill, L., Bretagnolle, V., … & Raynei, M. J. (2014). Pelagic distribution of Gould’s Petrel (Pterodroma leucoptera): linking shipboard and onshore observations with remote-tracking data. Emu-Austral Ornithology, 114(4), 360–370.

R Core Team (2020). R: A language and environment for statistical computing. R Foundation for Statistical Computing, Vienna, Austria. https://www.R-project.org/.

Ramos, R., Carlile, N., Madeiros, J., Ramírez, I., Paiva, V. H., Dinis, H. A., … & González-Solís, J. (2017). It is the time for oceanic seabirds: Tracking year-round distribution of gadfly petrels across the Atlantic Ocean. Diversity and Distributions, 23(7), 794–805.

Randall, D. A., Pollinger, J. P., Argaw, K., Macdonald, D. W., & Wayne, R. K. (2010). Fine-scale genetic structure in Ethiopian wolves imposed by sociality, migration, and population bottlenecks. Conservation Genetics, 11(1), 89–101.

Rayner, M. J., Hauber, M. E., Steeves, T. E., Lawrence, H. A., Thompson, D. R., Sagar, P. M., … & Shaffer, S. A. (2011). Contemporary and historical separation of transequatorial migration between genetically distinct seabird populations. Nature Communications, 2(1), 1–7.

Rayner, M. J., Gaskin, C. P., Fitzgerald, N. B., Baird, K. A., Berg, M. M., Boyle, D., … & Ismar, S. M. (2015). Using miniaturized radiotelemetry to discover the breeding grounds of the endangered New Zealand Storm Petrel Fregetta maoriana. Ibis, 157(4), 754–766.

Rayner, M. J., Carlile, N., Priddel, D., Bretagnolle, V., Miller, M. G. R., Phillips, R. A., … & Torres, L. G. (2016). Niche partitioning by three Pterodroma petrel species during non-breeding in the equatorial Pacific Ocean. Marine Ecology Progress Series, 549, 217–229.

Rayner, M. J., Baird, K. A., Bird, J., Cranwell, S., Raine, A. F., Maul, B., … & Gaskin, C. P. (2020). Land and sea-based observations and first satellite tracking results support a New Ireland breeding site for the Critically Endangered Beck’s Petrel Pseudobulweria beckii. Bird Conservation International, 30(1), 58–74.

Ribeiro, A. M., Lloyd, P., Feldheim, K. A., & Bowie, R. C. (2012). Microgeographic socio-genetic structure of an African cooperative breeding passerine revealed: integrating behavioural and genetic data. Molecular Ecology, 21(3), 662–672.

Richard, Y., Abraham, E. R., & Berkenbusch, K. (2017). Assessment of the risk of commercial fisheries to New Zealand seabirds, 2006-07 to 2014-15. Ministry for Primary Industries, Manatu Ahu Matua.

Richards, C., Cooke, R. S., & Bates, A. E. (2021). Biological traits of seabirds predict extinction risk and vulnerability to anthropogenic threats. Global Ecology and Biogeography, 30(5), 973–986.

Rodríguez, A., Arcos, J. M., Bretagnolle, V., Dias, M. P., Holmes, N. D., Louzao, M., … & Chiaradia, A. (2019). Future directions in conservation research on petrels and shearwaters. Frontiers in Marine Science, 6:94.

Rolshausen, G., Segelbacher, G., Hobson, K. A., & Schaefer, H. M. (2009). Contemporary evolution of reproductive isolation and phenotypic divergence in sympatry along a migratory divide. Current Biology, 19(24), 2097–2101.

Ronconi, R. A., Allard, K. A., & Taylor, P. D. (2015). Bird interactions with offshore oil and gas platforms: review of impacts and monitoring techniques. Journal of Environmental Management, 147, 34–45.

Ryan, P. G., Bourgeois, K., Dromzée, S., & Dilley, B. J. (2014). The occurrence of two bill morphs of prions Pachyptila vittata on Gough Island. Polar Biology, 37(5), 727–735.

Ryan, P. G., Ryan, E. M., & Glass, J. P. (2021). Dazzled by the light: the impact of light pollution from ships on seabirds at Tristan da Cunha. Ostrich, 92(3), 218–224.

Satgé, Y. G., Rupp, E., & Jodice, P. G. (2019). A preliminary report of ongoing research of the ecology of Black-capped Petrel (Pterodroma hasitata) in Sierra de Bahoruco, Dominican Republic–I: GPS tracking of breeding adults. South Carolina Cooperative Fish and Wildlife Research Unit.

Sato, F., Karino, K., Oshiro, A., Sugawa, H., & Hirai, M. (2010). Breeding of Swinhoe’s Storm-petrel Oceanodroma monorhis in the Kitsujima Islands, Kyoto, Japan. Marine Ornithology, 38, 133–136.

Sawilowsky, S. S. (2009). New effect size rules of thumb. Journal of Modern Applied Statistical Methods, 8(2), 26.

Seidov, D., Mishonov, A., Reagan, J., & Parsons, R. (2019). Resilience of the Gulf Stream path on decadal and longer timescales. Scientific Reports, 9(1), 1–9.

Shaffer, S. A., Tremblay, Y., Weimerskirch, H., Scott, D., Thompson, D. R., Sagar, P. M., … & Costa, D. P. (2006). Migratory shearwaters integrate oceanic resources across the Pacific Ocean in an endless summer. Proceedings of the National Academy of Sciences, 103(34), 12799–12802.

Shimada, T., Thums, M., Hamann, M., Limpus, C. J., Hays, G. C., FitzSimmons, N. N., … & Meekan, M. G. (2021). Optimising sample sizes for animal distribution analysis using tracking data. Methods in Ecology and Evolution, 12(2), 288–297.

Simons, T. R., Lee, D. S., & Haney, J. C. (2013). Diablotin Pterodroma hasitata: a biography of the endangered Black-capped Petrel. Marine Ornithology, 41, 1–43.

Soanes, L. M., Arnould, J. P., Dodd, S. G., Sumner, M. D., & Green, J. A. (2013). How many seabirds do we need to track to define home-range area?. Journal of Applied Ecology, 50(3), 671–679.

Spalding, M. D., Fox, H. E., Allen, G. R., Davidson, N., Ferdaña, Z. A., Finlayson, M. A. X., … & Robertson, J. (2007). Marine ecoregions of the world: a bioregionalization of coastal and shelf areas. BioScience, 57(7), 573–583.

Sussman, Allison & US Geological Survey (2014). Atlantic Offshore Seabird Dataset Catalog, Atlantic Coast and Outer Continental Shelf, from 1938-01-01 to 2013-12-31 (NCEI Accession 0115356).

NOAA National Centers for Environmental Information. Dataset. Accessed from https://accession.nodc.noaa.gov/0115356 on 01 November 2020.

Tartu, S., Goutte, A., Bustamante, P., Angelier, F., Moe, B., Clément-Chastel, C., … & Chastel, O. (2013). To breed or not to breed: endocrine response to mercury contamination by an Arctic seabird. Biology Letters, 9(4), 20130317.

Taylor, G. A., Elliott, G. P., Walker, K.J., & Bose, S. (2011). Year-round distribution, breeding cycle, and activity of white-headed petrels (Pterodroma lessonii). Notornis 67(2011), 369–386

Taylor, R. S., & Friesen, V. L. (2017). The role of allochrony in speciation. Molecular Ecology, 26(13), 3330–3342.

Tew Kai, E., & Marsac, F. (2010). Influence of mesoscale eddies on spatial structuring of top predators’ communities in the Mozambique Channel. Progress in Oceanography, 86(1-2), 214–223.

Tomkins, R. J., & Milne, B. J. (1991). Differences among Dark-rumped Petrel (Pterodroma phaeopygia) populations within the Galapagos Archipelago. Notornis, 38(1), 1–35.

Torchiano M (2020). effsize: Efficient Effect Size Computation. R package version 0.8.1, https://CRAN.R-project.org/package=effsize.

Torres, L. G., Sagar, P. M., Thompson, D. R., & Phillips, R. A. (2013). Scaling down the analysis of seabird-fishery interactions. Marine Ecology Progress Series, 473, 275–289.

Trebilco, R., Gales, R., Lawrence, E., Alderman, R., Robertson, G., & Baker, G. B. (2010). Characterizing seabird bycatch in the eastern Australian tuna and billfish pelagic longline fishery in relation to temporal, spatial and biological influences. Aquatic Conservation: Marine and Freshwater Ecosystems, 20(5), 531–542.

U.S. Fish and Wildlife Service (2018). Endangered and Threatened Wildlife and Plants; Threatened Species Status for Black-capped Petrel: 83 FR 50560. Federal Register. 83(195): 50560–50574

VLIZ (Flanders Marine Institute) (2019). Maritime boundaries geodatabase, version 11. Accessed from www.marineregions.org/ on 1 July 2021.

Wallace, S. J., Morris-Pocock, J. A., González-Solís, J., Quillfeldt, P., & Friesen, V. L. (2017). A phylogenetic test of sympatric speciation in the Hydrobatinae (Aves: Procellariiformes). Molecular Phylogenetics and Evolution, 107, 39–47.

Waugh, S. M., Filippi, D. P., Kirby, D. S., Abraham, E., & Walker, N. (2012). Ecological Risk Assessment for seabird interactions in Western and Central Pacific longline fisheries. Marine Policy, 36(4), 933–946.

Weimerskirch, H. (2007). Are seabirds foraging for unpredictable resources?. Deep Sea Research Part II: Topical Studies in Oceanography, 54(3-4), 211–223.

Wheeler, J., Satgé, Y., Brown, A., Goetz, J., Keitt, B., Nevins, H., & Rupp, E. (2021). Black-capped Petrel (Pterodroma hasitata) Conservation Update and Action Plan: Conserving the Diablotin. International Black-capped Petrel Conservation Group, BirdsCaribbean.

Wiley, A. E., Welch, A. J., Ostrom, P. H., James, H. F., Stricker, C. A., Fleischer, R. C., … & Swindle, K. A. (2012). Foraging segregation and genetic divergence between geographically proximate colonies of a highly mobile seabird. Oecologia, 168(1), 119–130.

Winker, K. (2010). Chapter 1: Subspecies represent geographically partitioned variation, a gold mine of evolutionary biology, and a challenge for conservation. Ornithological Monographs, 67(1), 6–23.

Winship, A. J., Kinlan, B. P., White, T. P., Leirness, J., & Christensen, J. (2018). Modeling At-Sea Density of Marine Birds to Support Atlantic Marine Renewable Energy Planning: Final Report. U.S. Department of the Interior, Bureau of Ocean Energy Management, Office of Renewable Energy Programs, Sterling, VA.

Wright, S. (1943). Isolation by distance. Genetics, 28(2), 114.

Wu, L., Xu, Y., Wang, Q., Wang, F., & Xu, Z. (2017). Mapping global shipping density from AIS data. The Journal of Navigation, 70(1), 67–81.

Zhang, Y., Jaeglé, L., Thompson, L., & Streets, D. G. (2014). Six centuries of changing oceanic mercury. Global Biogeochemical Cycles, 28(11), 1251–1261.

Zhou, C., Jiao, Y., & Browder, J. (2019). Seabird bycatch vulnerability to pelagic longline fisheries: ecological traits matter. Aquatic Conservation: Marine and Freshwater Ecosystems, 29(8), 1324–1335.

