## Supporting Online Information for "Temporal and spatial segregations between phenotypes of the Diablotin Black-capped Petrel *Pterodroma hasitata* during the breeding and non-breeding periods"

A. Additional methods and results supporting the assessment of colony visits.

B. Supplementary tables and figures.

### A. Additional methods and results supporting the assessment of colony visits.

#### Methods

Satellite transmitters can inform about visits to general breeding areas but they may lack temporal and spatial precision to confirm breeding activity based solely on location data (Northrup et al. 2018). Indeed, the quality of locations obtained by Doppler effect (as is the case with PTT data) is unpredictable and depends on the number and location of Argos satellites within view and reach during the limited period when the PTT is on (CLS 2016). Therefore, in seabirds that spend time underground at the breeding site, the number and quality of locations in breeding areas is generally decreased. Furthermore, the operation of solar-powered transmitters is limited by the amount of solar radiations received because their batteries cannot fully sustain an adequate voltage when exposed to low or limited light levels (Ens et al. 2008, Spencer et al. 2014). Therefore, with solar-powered transmitters, it is possible to reversely use voltage data to estimate periods of low light input, and infer breeding activity in species breeding in densely forested areas and/or underground. Black-capped Petrels forage actively during daylight hours (Haney 1987, Simons et al. 2013) therefore attached solar-powered transmitters can recharge adequately during flight and foraging activity. Black-capped Petrels access breeding sites after dusk (Jodice et al. 2015) and, when at the colony, are active at night but spend daylight hours underground (Simons et al. 2013). Thus, when at breeding sites, attached solar-powered PTTs cannot adequately communicate with satellites and cannot adequately recharge. To determine if tracked petrels may have been visiting breeding sites, we assessed if 1) petrels were located in the vicinity of known breeding sites in the Caribbean, 2) scheduled communications failed to occur, and 3) battery levels showed periods of low voltage. We assumed that locations near known breeding sites were likely to indicate some form of breeding activity (e.g., prospecting or nest initiation). We also assumed that breeding activity (e.g., occupancy of a burrow) would lead to gaps in communications between the satellite tag and satellite system and to low voltage levels of the satellite tags. Therefore, for locations within a month of a suspected visit to a breeding site, we (1) calculated distance to the nearest breeding site with function *distGeo* in package *geosphere* in R (Hijmans 2019), (2) used raw PTT data (location and metadata collection) to assess any gap in satellite communication, and (3) compared mean daily voltage with the overall mean voltage during that period.

#### Results

Two PTTs were still transmitting at the onset of the breeding season in September 2019 (Figure 4:  ID 442 (male dark form) and ID 462 (male light form). Both individuals subsequently made southerly trips to the Caribbean (Figure 3). ID 462 left pelagic waters along Georges Bank between 21 – 22 September and reached the Puerto Rico Trench on 27 September, going through the Mona Passage between 27 – 28 September. On 29 September, a class-A Argos location (the only location recorded over land for this individual) was recorded ca. 8 km to the east of the known breeding area of Valle Nuevo National Park, Dominican Republic. This location had an error radius of 21.5 km (which includes coastal areas in the Dominican Republic) and was not retained by our continuous-time random walk state-space model. On 1 October, ID 462 was 122 km away from the nearest known breeding site in Valle Nuevo National Park, Dominican Republic (Figure 4A). Between 1 – 15 October, scheduled satellite communications did not occur, except for a short burst on 8 October: on that day, ID 462 was 118 km (range: 81-152 km, n = 8) directly south of a known breeding site of Valle Nuevo, Dominican Republic, and the battery level was 3.58 V (mean overall battery voltage: $\bar{x}_{2019\text{-}01\text{-}01 – 2019\text{-}10\text{-}15}$ = 3.91 V ± 0.10)(Figure 4B). On 15 October, ID 462 was heading north of the Mona Passage, 248 km from the Valle Nuevo site, and the battery level was 3.8 V. ID 462 made multiple stops in the southern reaches of the Sargasso Sea, along the Antilles Current and, on 1 November, it had returned to pelagic waters off the Virginian ecoregion, where it remained until tracking stopped on 14 November. Battery levels stayed high for the remainder of the tracking duration ($\bar{x}_{2019\text{-}10\text{-}16 – 2019\text{-}11\text{-}14}$ = 3.88 V; range: 3.66-4.10 V), and satellite communications occurred as scheduled. Between 27 September and 14 November, the accuracy of locations was < 10km. Based on this information, we inferred that ID 462 was at a breeding site during 2 – 8 October and 9 – 15 October.

ID 442 made two trips to the Caribbean. Between 31 October and 2 November, it left Gulf Stream waters to the northeast of Hatteras and travelled southward through the Hatteras Plains and the Sargasso Sea. On 7 November, it reached the Turks and Caicos shelf, and went through the Windward Passage. On 9 November, ID 442 was 83 km (range: 74-110 km, n = 4) southwest of the nearest known breeding site in La Visite escarpment of Haiti (Figure 4A). Between 9 – 22 November, scheduled satellite communications did not occur (Figure 4B). On 22 November, ID 442 was 114 km south of La Visite (range: 114-115 km, n = 4), and the battery level was 3.84 V ($\bar{x}_{2019\text{-}10\text{-}15 – 2019\text{-}12\text{-}29}$ = 4.04 V ± 0.06). Between 24 – 27 November, ID 442 stayed in neritic and semi-pelagic waters south and west of the Haitian peninsula, and within 251-261 km of La Visite. At the start of 29 November, ID 442 was within range: 6 – 7 km (n = 2 locations) of the La Visite breeding site. Satellite communications failed to occur between 29 November and 3 December. On December 3, ID 442 was 160 km (range: 139-213 km) southwest of La Visite, travelling west, and the battery level was 4.1 V. On 5 December, it passed through the Windard Passage and travelled through the western extent of the Hatteras Plains to Gulf Stream Waters south of Cape Hatteras. It remained there until 14 December, when it initiated a second trip to the Caribbean. Limited details are available but, on 17 December, ID 442 was travelling south through the Antilles Current. On 19 December, it was located 2.5-10 km offshore of the southcentral coast of the Haitian peninsula and 86 km (range: 68-114 km, n = 6) away from La Visite; the battery level was 4.05 V. On 21 December, ID 442 was travelling north through the Windward Passage, 291 km (range: 283-302 km, n = 8) away from La Visite; the battery level was 3.95 V. On 25 December, it had returned to Gulf Stream waters of the Carolinian ecoregion. Except for a trip to the vicinity of the Caryn Seamount, in the northern extent of the Hatteras Plains, ID 442 remained there until tracking stopped on 24 January 2020. Battery levels stayed high for the remainder of the tracking duration ($\bar{x}_{2019\text{-}12\text{-}21 – 2019\text{-}01\text{-}24}$ = 3.98 V; range: 3.85-4.12 V), and satellite communications occurred as scheduled. Between 31 October and 24 January 2020, the accuracy of locations was < 10km. Based on this information, we infer that ID 442 was at a breeding site during 9 – 22 November, and 29 November – 3 December. It remains inconclusive whether it visited a breeding site between 19 – 21 December.

Any use of trade, firm, or product names is for descriptive purposes only and does not imply endorsement by the U.S. Government.

**References**

CLS. (2016). Argos User's Manual: Worldwide tracking and environmental monitoring by satellite.

Cohen, J., 1988. Statistical power analysis for the behavioral sciences. Lawrence Erlbaum Associates. ESBN 0-8058-0283-5.

Ens B.J., Bairlein F., Camphuysen C.J., Boer P. de, Exo K.-M., Gallego N., Hoye B., Klaassen R., Oosterbeek K., Shamoun-Baranes J., Jeugd H. van der & Gasteren H. van. (2008).

Tracking of individual birds. Report on WP 3230 (bird tracking sensor characterization) and WP 4130 (sensor adaptation and calibration for bird tracking system) of the FlySafe basic activities project. SOVON-onderzoeksrapport 2008/10. SOVON Vogelonderzoek Nederland, Beek-Ubbergen.

Haney, J. C. (1987). Aspects of the pelagic ecology and behavior of the black-capped petrel(Pterodroma hasitata). *Wilson bulletin*, *99*(2), 153-312.

Hijmans, R.J. (2019). *geosphere: Spherical Trigonometry*. R package version 1.5-10. https://CRAN.R-project.org/package=geosphere

Jodice, P. G., Ronconi, R. A., Rupp, E., Wallace, G. E., & Satgé, Y. (2015). First satellite tracks of the endangered black-capped petrel. *Endangered Species Research*, *29*(1), 23-33.

Northrup, J. M., Rivers, J. W., Nelson, S. K., Roby, D. D., & Betts, M. G. (2018). Assessing the utility of satellite transmitters for identifying nest locations and foraging behavior of the threatened Marbled Murrelet Brachyramphus marmoratus. *Marine Ornithology*, *46*, 47-55.

Simons, T. R., Lee, D. S., & Haney, J. C. (2013). Diablotin Pterodroma hasitata: a biography of the endangered Black-capped Petrel. *Marine Ornithology*, *41*, 1-43.

Spencer, N. C., Gilchrist, H. G., & Mallory, M. L. (2014). Annual movement patterns of endangered ivory gulls: the importance of sea ice. *PLoS One*, *9*(12), e115231.

**B. Supplementary tables and figures**

Table S1. Number and accuracy of locations of tracked Black-capped Petrels captured off Cape Hatteras, North Carolina, USA, May 2019, based on Argos location classes (LC).

| **Bird ID** [a] | **No. of locations** | **LC 3 < 250m** | **LC 2 250-500m** | **LC 1 500-1500m** | **LC 0 > 1500m** | **LC A -** | **LC B -** |
|---|---|---|---|---|---|---|---|
| **441** | 442 | 16 | 42 | 84 | 168 | 39 | 93 |
| **442** | 1132 | 45 | 100 | 222 | 447 | 101 | 215 |
| 462 | 916 | 83 | 99 | 155 | 342 | 64 | 173 |
| 463 | 84 | 5 | 7 | 15 | 30 | 10 | 17 |
| **464** | 309 | 28 | 32 | 67 | 105 | 28 | 48 |
| 465 | 374 | 53 | 58 | 78 | 84 | 27 | 74 |
| **466** | 55 | 5 | 3 | 10 | 15 | 9 | 13 |
| 467 | 695 | 65 | 82 | 139 | 236 | 60 | 113 |
| 468 | 566 | 32 | 42 | 119 | 214 | 51 | 108 |
| **469** | 83 | 1 | 6 | 12 | 49 | 5 | 10 |
| **Total** [b] | 4656 | 333 | 471 | 901 | 1690 | 394 | 864 |
| **Proportion** [c] **(%)** | 100.0 | 7.2 | 10.1 | 19.4 | 36.3 | 8.5 | 18.6 |
| **Error radius (m)** | | | | | | | |
| **Mean** | | 192.1 | 354.6 | 975.6 | 5288.1 | | |
| **Minimum** | | 127 | 250 | 501 | 1502 | | |
| **Maximum** | | 249 | 500 | 1500 | 216041 | | |

[a] Bold lettering indicates dark phenotypes.
[b] Total number of locations includes n = 3 location classes Z. Location class (3, 2, 1, 0, B, A, Z, in decreasing order of quality) was assigned by Argos.
[c] Total proportion of locations includes 0.1% of location classes Z.

Table S2. Area and proportion of overlap, and Bhattacharyya's affinity for core area (50%UD) and home range (90%UD) of dark and light forms of Black-capped Petrels tracked from May 2019 – August 2019. UD = Utilization distribution, BA = Bhattacharyya's affinity.

| | **Area ($km^2$)** | | | **Proportion of UD overlapping (%)** | | |
|---|---|---|---|---|---|---|
| **UD** | **Dark** | **Light** | **Overlap** | **Dark** | **Light** | **BA** |
| 50 | 88058.1 | 81144.8 | 5399.9 | 6.13 | 6.65 | 0.03 |
| 90 | 292472.4 | 355915.0 | 104022.3 | 35.57 | 29.23 | 0.29 |

Table S3. Distance of tracking locations to nearest marine energy lease area, and area and proportion of core area (50% UD) and home range (90% UD) of petrels within marine energy lease areas in the United States and Canada. Distance to nearest lease was calculated from tracking locations between May 2019 and January 2020; overlap with leases was calculated from tracking locations between May – August 2019. Bold lettering denotes overlap. UD = Utilization distribution.

| | | Overall | Dark | | | | Light | | | |
|---|---|---|---|---|---|---|---|---|---|---|
| | | | 50% UD | | 90% UD | | 50% UD | | 90% UD | |
| | | **Distance to nearest lease (km)** | **Area of overlap (km2)** | **Prop. in lease area (%)** | **Area of overlap (km2)** | **Prop. in lease area (%)** | **Area of overlap (km2)** | **Prop. in lease area (%)** | **Area of overlap (km2)** | **Prop. in lease area (%)** |
| **Wind** | | | | | | | | | | |
| Active | Delaware | 94.0 | 0 | 0 | 0 | 0 | 0 | 0 | 0 | 0 |
| | Massachusetts[a] | 94.7 | 0 | 0 | 0 | 0 | 0 | 0 | 0 | 0 |
| | Maryland | 74.2 | 0 | 0 | 0 | 0 | 0 | 0 | 0 | 0 |
| | **North Carolina** | **29.5** | **0** | **0** | **138.5** | **0.05** | **0** | **0** | **0** | **0** |
| | New Jersey | 106.2 | 0 | 0 | 0 | 0 | 0 | 0 | 0 | 0 |
| | New York | 135.7 | 0 | 0 | 0 | 0 | 0 | 0 | 0 | 0 |
| | Virginia | 65.0 | 0 | 0 | 0 | 0 | 0 | 0 | 0 | 0 |
| Planned | North Carolina | 70.8 | 0 | 0 | 0 | 0 | 0 | 0 | 0 | 0 |
| | New Jersey | 80.4 | 0 | 0 | 0 | 0 | 0 | 0 | 0 | 0 |
| | New York | 90.2 | 0 | 0 | 0 | 0 | 0 | 0 | 0 | 0 |
| | South Carolina | 76.7 | 0 | 0 | 0 | 0 | 0 | 0 | 0 | 0 |
| Drafted[b] | **OCS** | **14.3** | **0** | **0** | **669.0** | **0.2** | **0** | **0** | **0** | **0** |
| | **CR North** | **0.0** | **1157.5** | **1.3** | **6211.3** | **2.1** | **748.0** | **0.9** | **6363.6** | **1.8** |
| | **CR South** | **0.0** | **2787.0** | **3.2** | **3089.3** | **1.1** | **0** | **0** | **3089.3** | **0.9** |
| **Oil and gas[c]** | | | | | | | | | | |
| | NS: 2434R | 126.7 | 0 | 0 | 0 | 0 | 0 | 0 | 0 | 0 |
| | **NS: 2435** | **0.0** | **0** | **0** | **0** | **0** | **0** | **0** | **0** | **0** |
| | **NS: 2436** | **0.0** | **0** | **0** | **0** | **0** | **0** | **0** | **0** | **0** |
| | Newfoundland | 648.5 | 0 | 0 | 0 | 0 | 0 | 0 | 0 | 0 |

[a] Corresponds to lease areas in the U.S. states of Rhode Island and Massachusetts

[b] OCS (Outer Continental Shelf): corresponds to areas A-C in BOEM's "Central Atlantic DRAFT Call for Information and Nomination Area" (https://www.regulations.gov/document/BOEM-2022-0023-0001; accessed 1 May 2022); CR (Continental Rise) North: corresponds to area E; CR South: corresponds to area F.

[c] NS = Nova Scotia

Table S4. Proportion of lease areas proposed in BOEM's Central Atlantic DRAFT Call for Information and Nomination within core area (50% UD) and home range (90% UD) of dark and light forms of Black-capped Petrels tracked from May – August 2019. UD = Utilization distribution.

| | **Dark** | | | | **Light** | | | |
|---|---|---|---|---|---|---|---|---|
| | **50% UD** | | **90% UD** | | **50% UD** | | **90% UD** | |
| | **Area of overlap (km2)** | **Proportion in UD (%)** | **Area of overlap (km2)** | **Proportion in UD (%)** | **Area of overlap (km2)** | **Proportion in UD (%)** | **Area of overlap (km2)** | **Proportion in UD (%)** |
| OCS | 0 | 0 | 669.0 | 10.1 | 0 | 0 | 0 | 0 |
| OCR North | 1157.5 | 17.7 | 6211.3 | 94.8 | 748.0 | 11.4 | 6363.6 | 97.1 |
| OCR South | 2787.0 | 90.2 | 3089.3 | 100.0 | 0 | 0 | 3089.3 | 100.0 |

OCS (Outer Continental Shelf): corresponds to areas A-C in BOEM's "Central Atlantic DRAFT Call for Information and Nomination Area" (https://www.regulations.gov/document/BOEM-2022-0023-0001; accessed 1 May 2022); CR (Continental Rise) North: corresponds to area E; CR South: corresponds to area F.

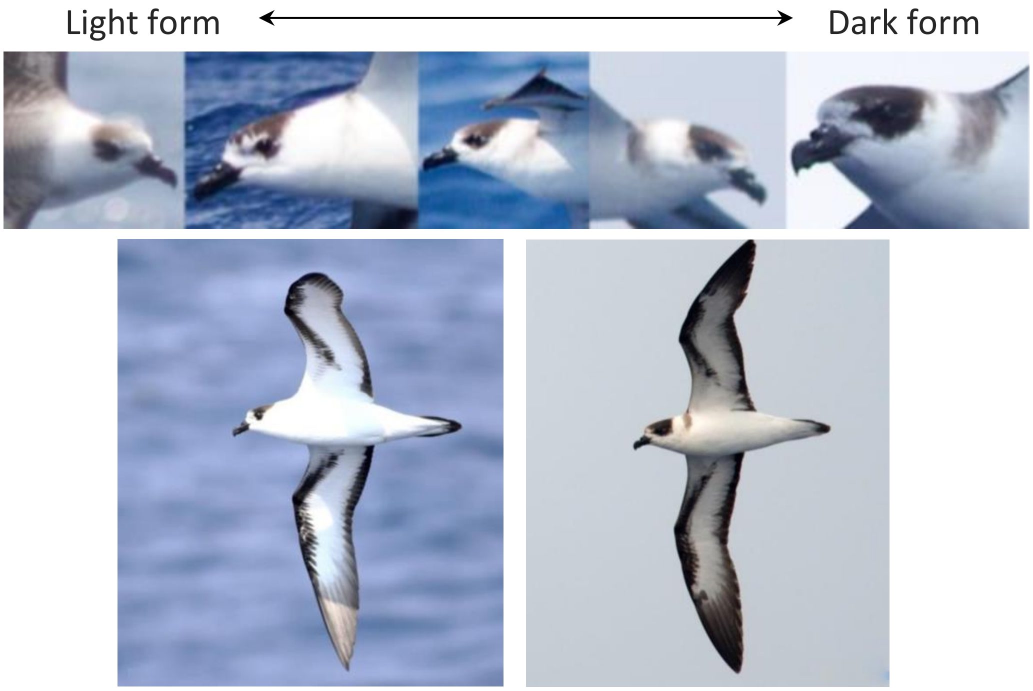


Figure S1. Photographs showing variation in Black-capped Petrel phenotypes. Image credits: Kate Sutherland.

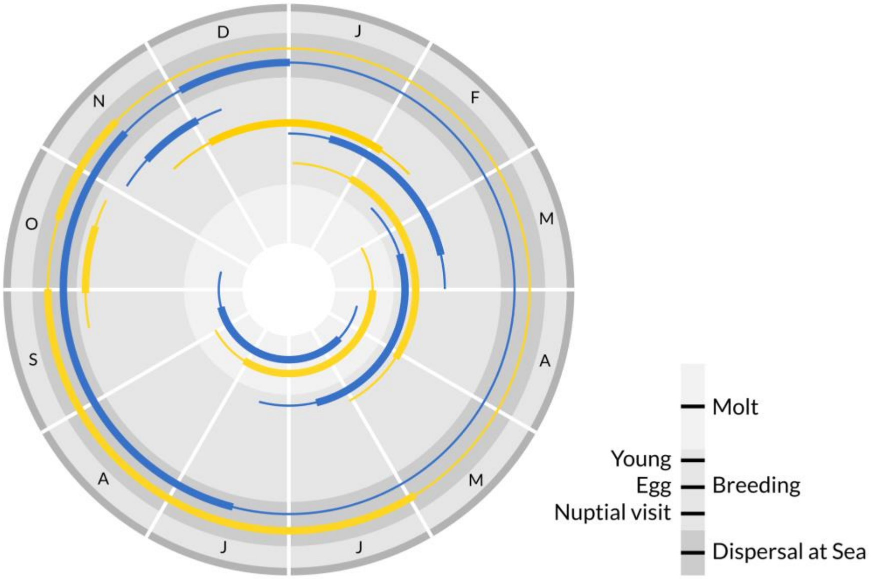


Figure S2. Phenology diagram for dark and light Black-capped Petrel forms. Blue: dark form; yellow: light form. Adapted from Simons et al. (2013). The diagram for the light form is based on Howell and Patteson (2008), Manly et al. (2013), Wheeler et al. (2021) and this study.

References:

Howell, S. N. G., & Patteson, J. B. (2008). Variation in the Black-capped Petrel–one species or more. *Alula*, *14*, 70-83.

Manly, B., Arbogast, B. S., Lee, D. S., & Van Tuinen, M. (2013). Mitochondrial DNA analysis reveals substantial population structure within the endangered Black-capped Petrel (*Pterodroma hasitata*). *Waterbirds*, *36*(2), 228-233.

Wheeler, J., Satgé, Y., Brown, A., Goetz, J., Keitt, B., Nevins, H., & Rupp, E. (2021). *Black-capped Petrel (*Pterodroma hasitata*) Conservation Update and Action Plan: Conserving the Diablotin*. International Black-capped Petrel Conservation Group, BirdsCaribbean.

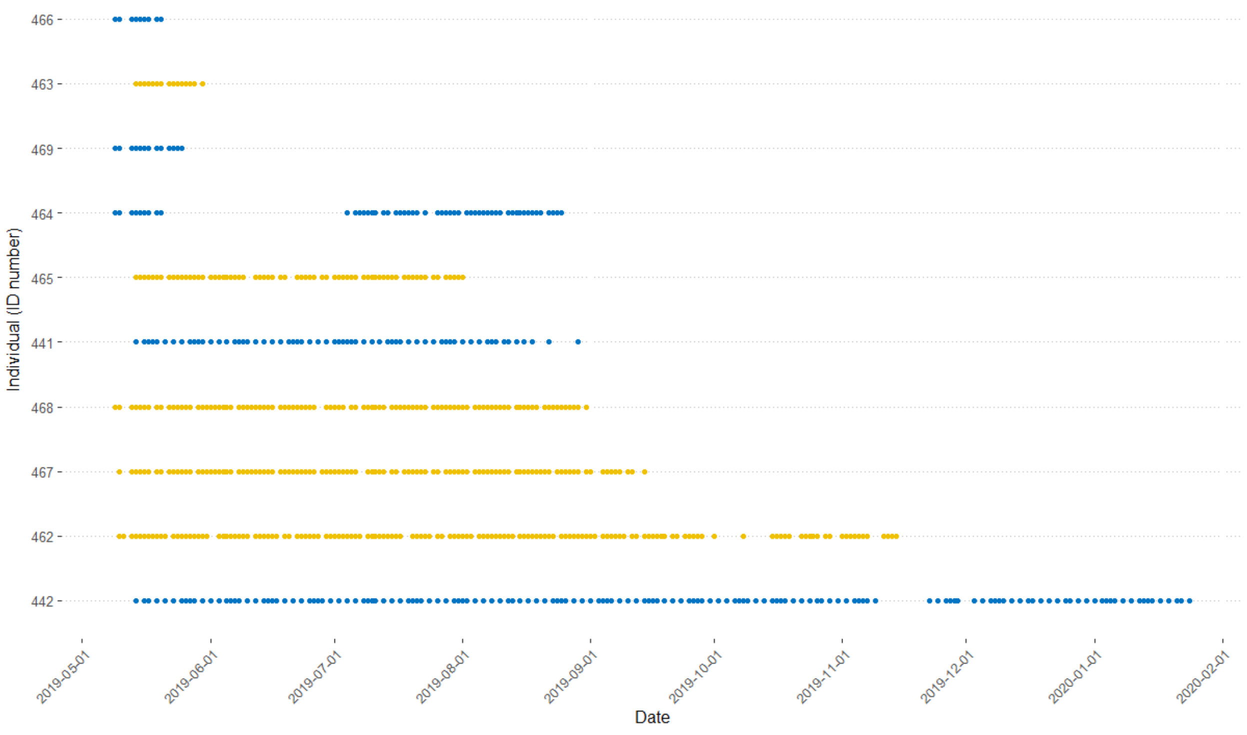


Figure S3. Tracking durations for Black-capped Petrels captured at sea off Cape Hatteras, North Carolina, USA, May 2019. Blue: dark form; yellow: light form. Individuals are ordered by tracking duration.

Figure S4 (next pages). Maps of individual movements of Black-capped Petrels tracked in 2019. In each map, the black circle represents the first tracked location, and the black triangle represents the last tracked location. Blue: dark form; yellow: light form. Black dashed lines indicate the general location of the western edge of the Gulf Stream. Dotted grey line indicates the -250-m isobath.

441: 2019−05−14 to 2019−08−29

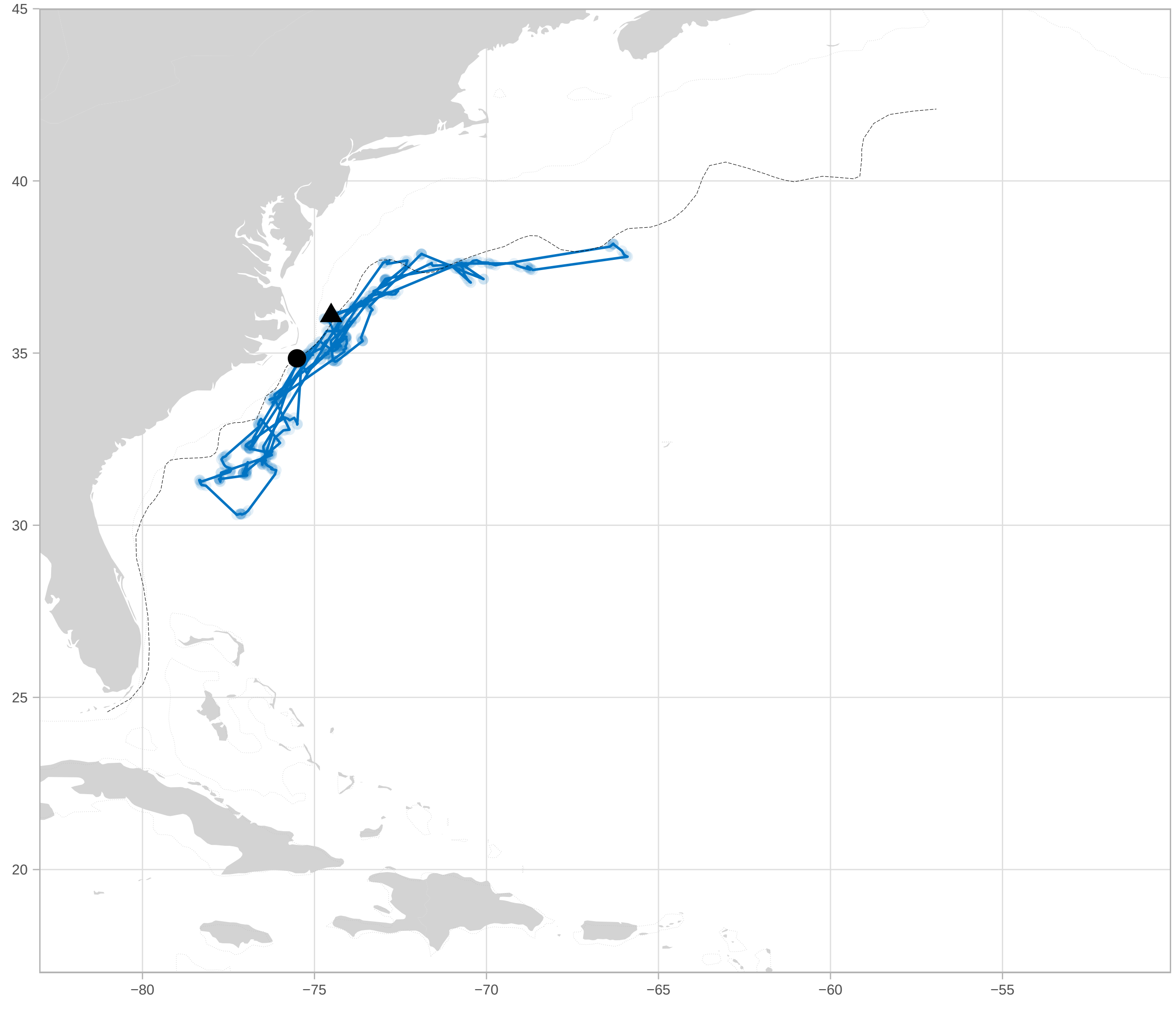

442: 2019−05−14 to 2020−01−24

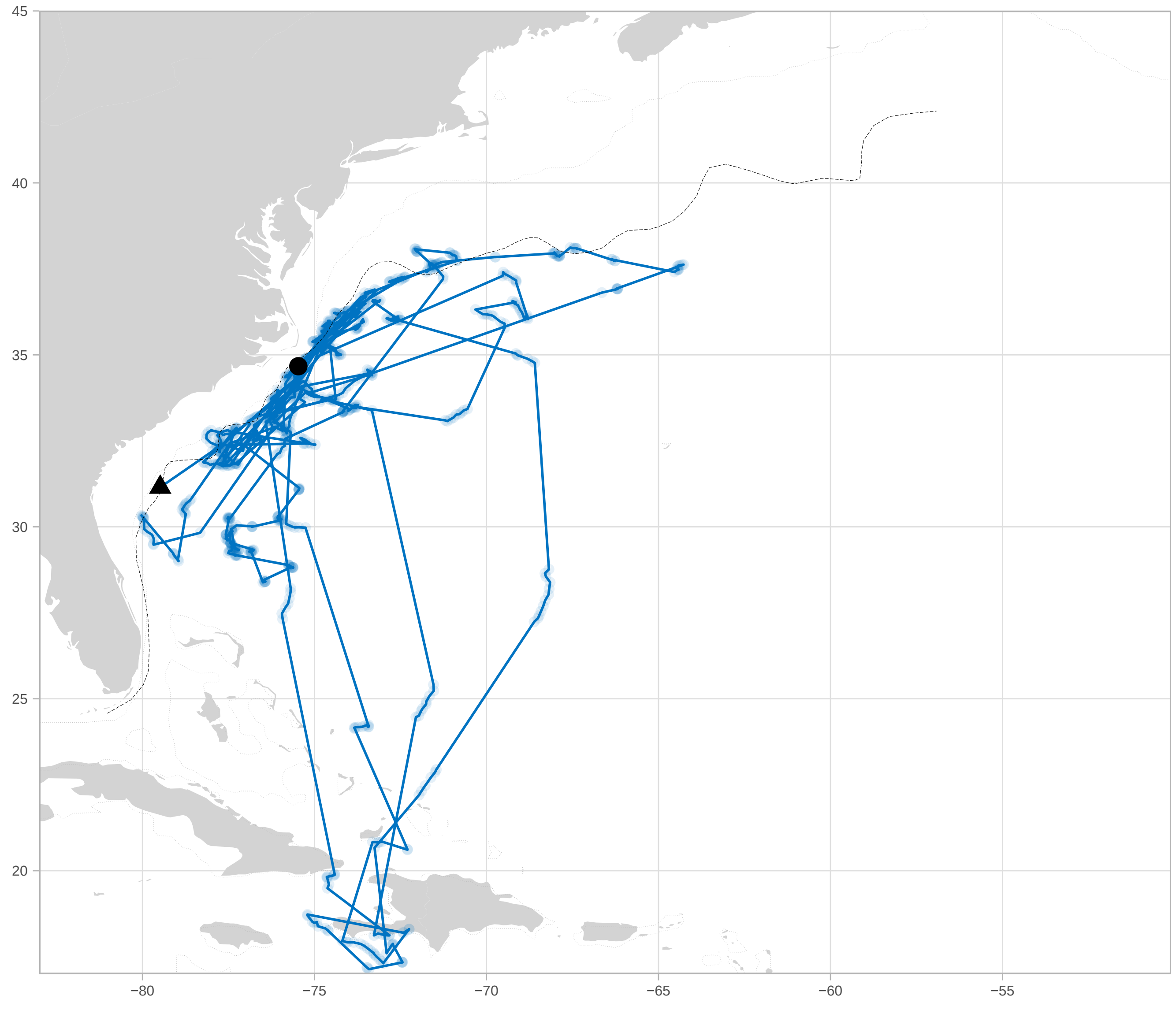

462: 2019−05−10 to 2019−11−14

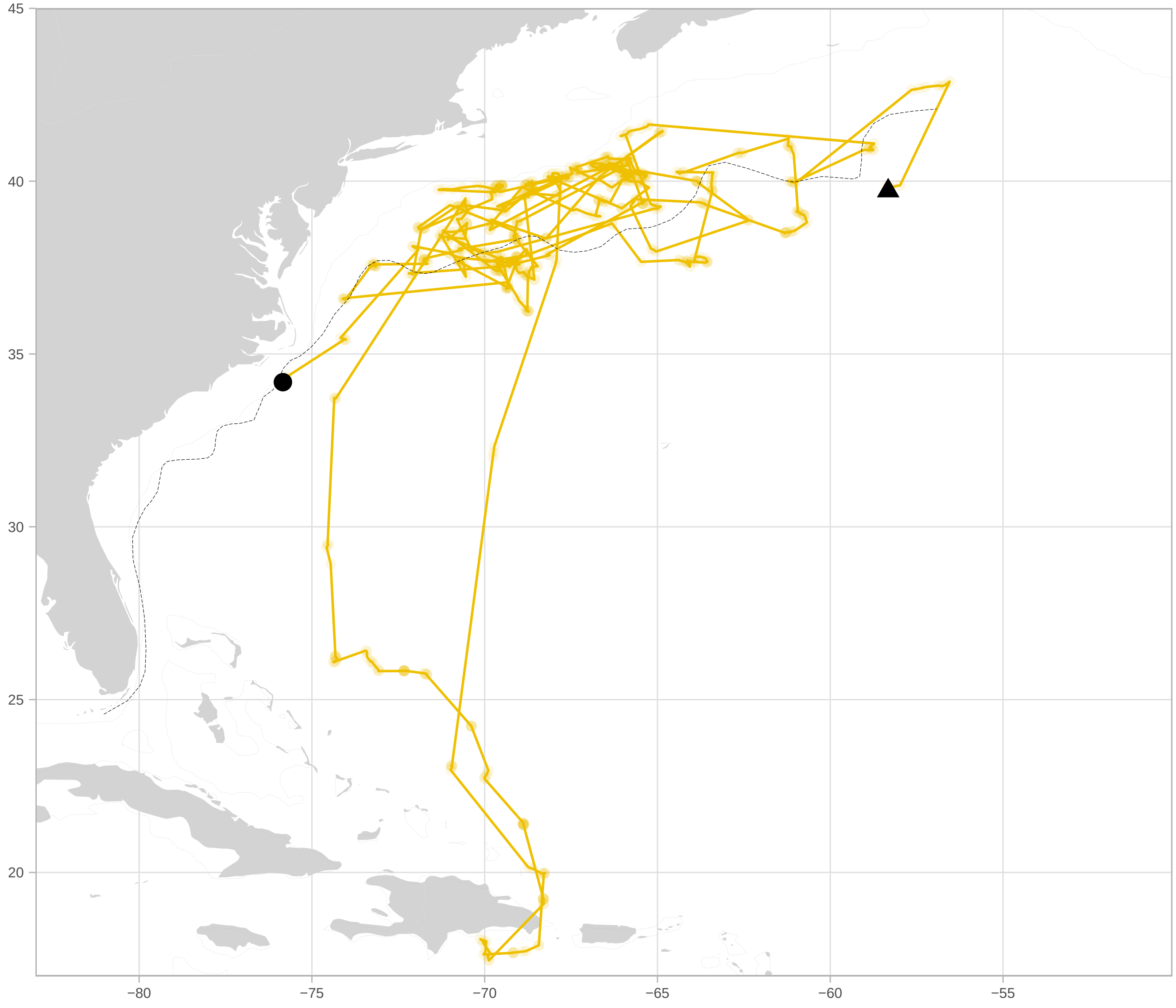

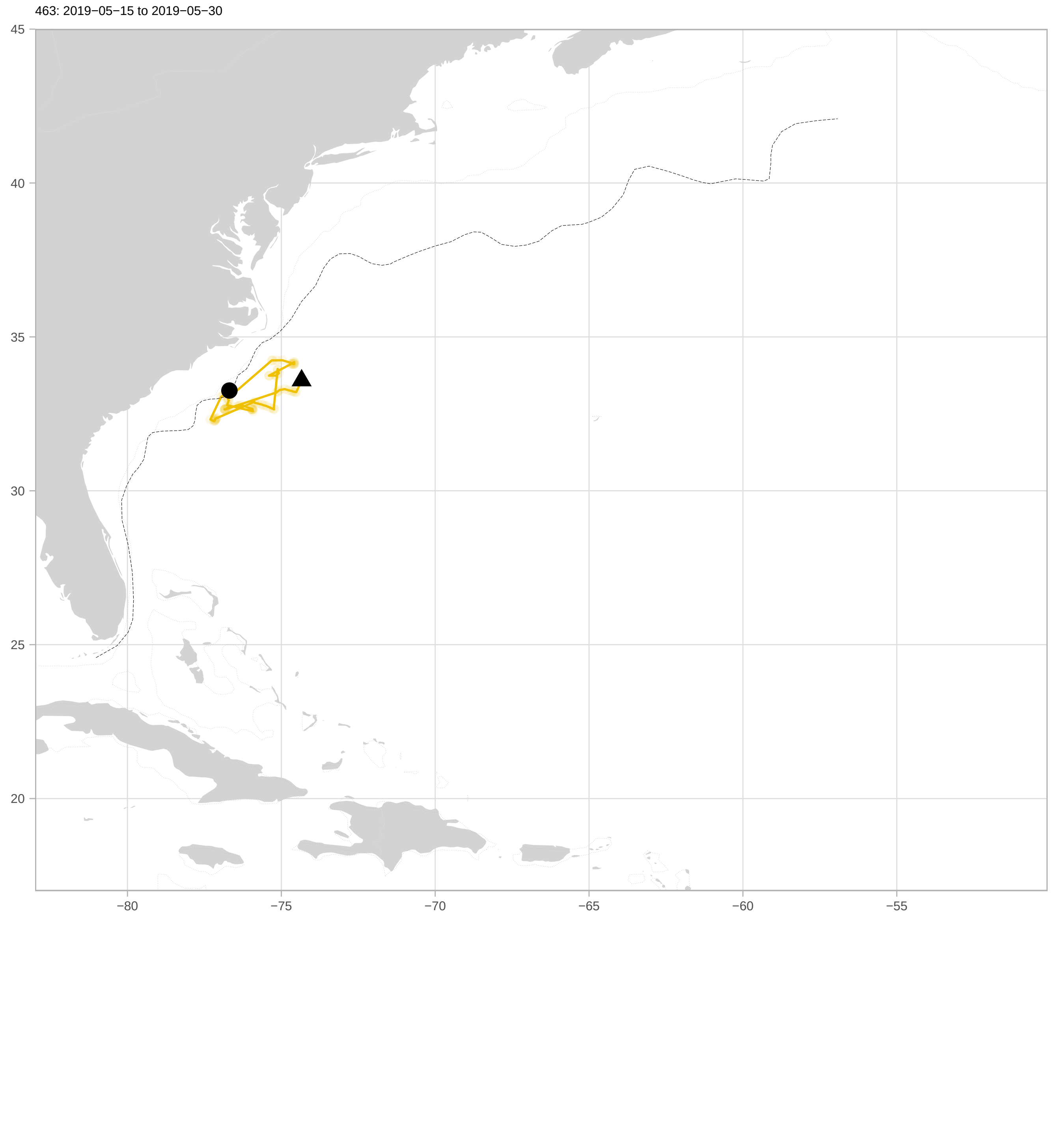
463: 2019−05−15 to 2019−05−30
45
40
35
30
25
20
−80
−75
−70
−65
−60
−55

464: 2019−05−09 to 2019−08−25

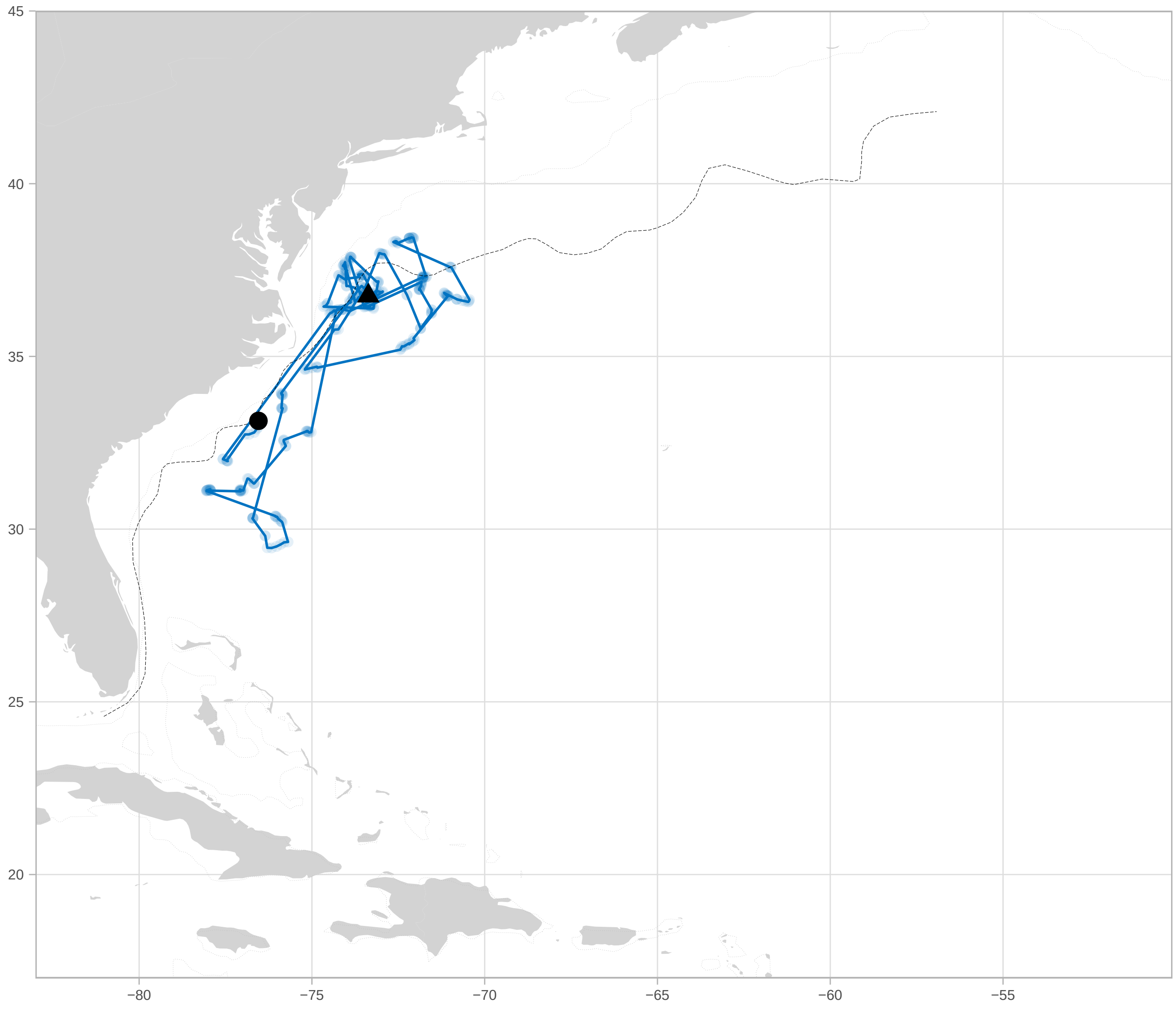

465: 2019−05−15 to 2019−08−01

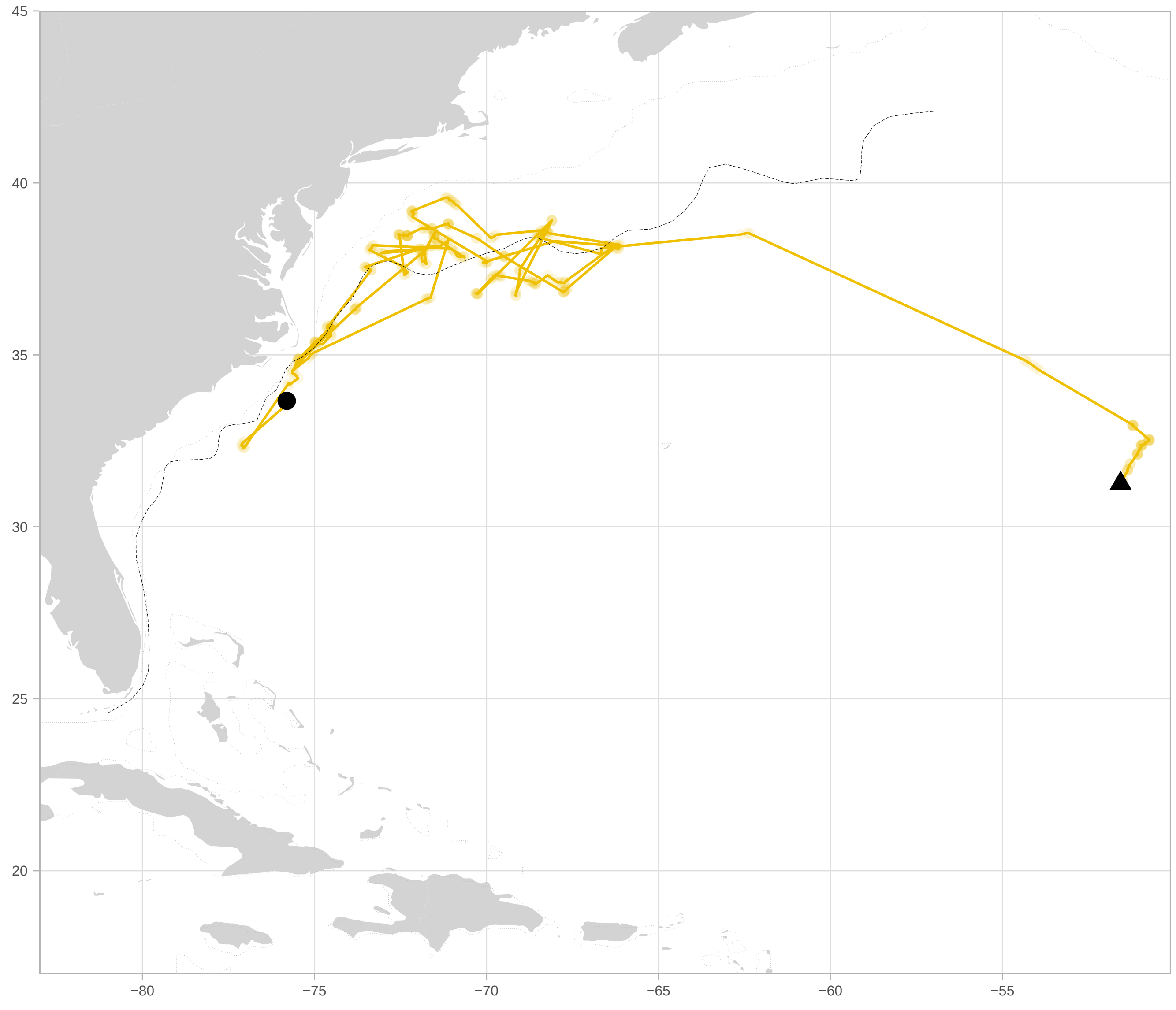

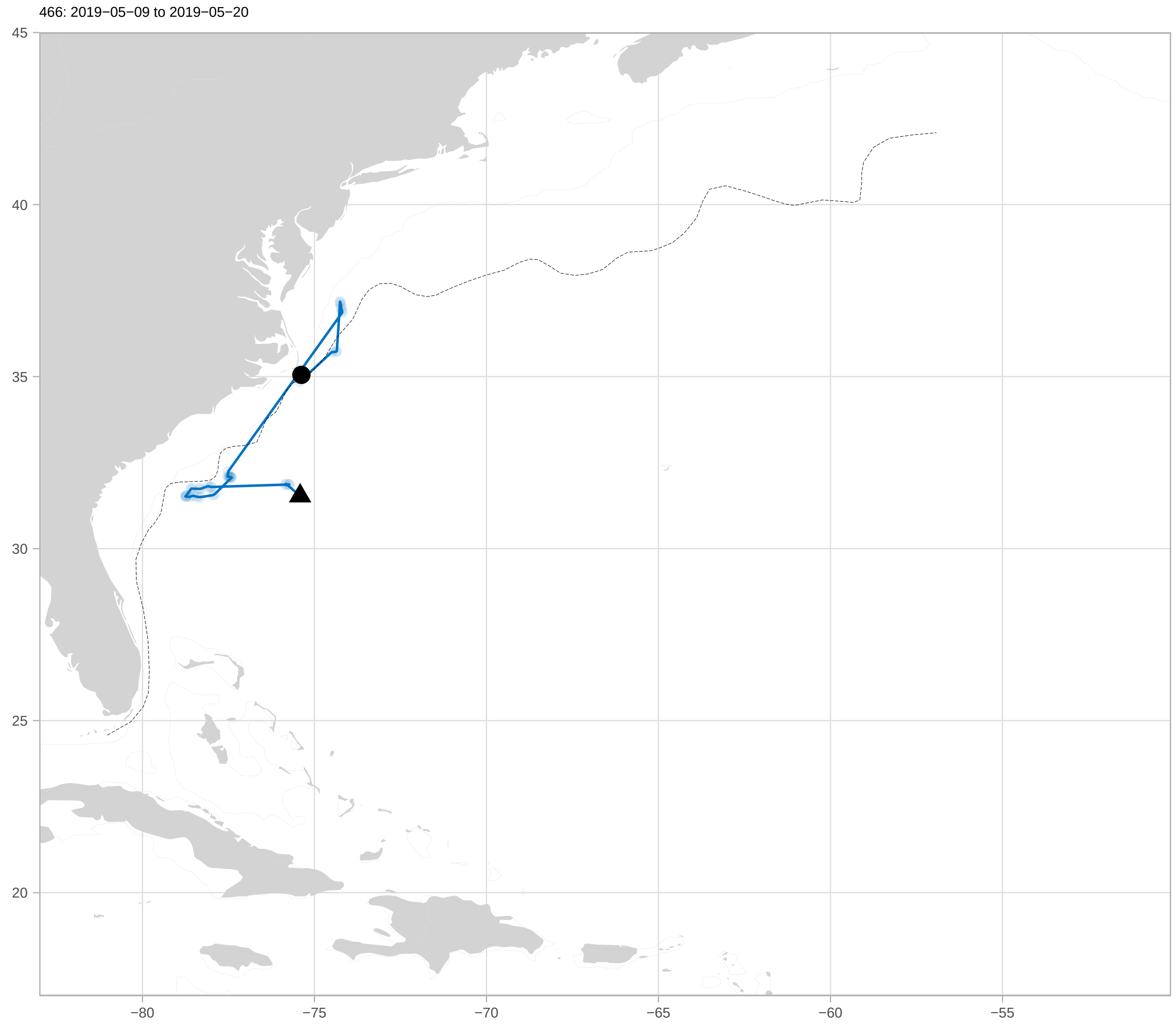
466: 2019−05−09 to 2019−05−20
45
40
35
30
25
20
−80
−75
−70
−65
−60
−55

467: 2019−05−10 to 2019−09−14

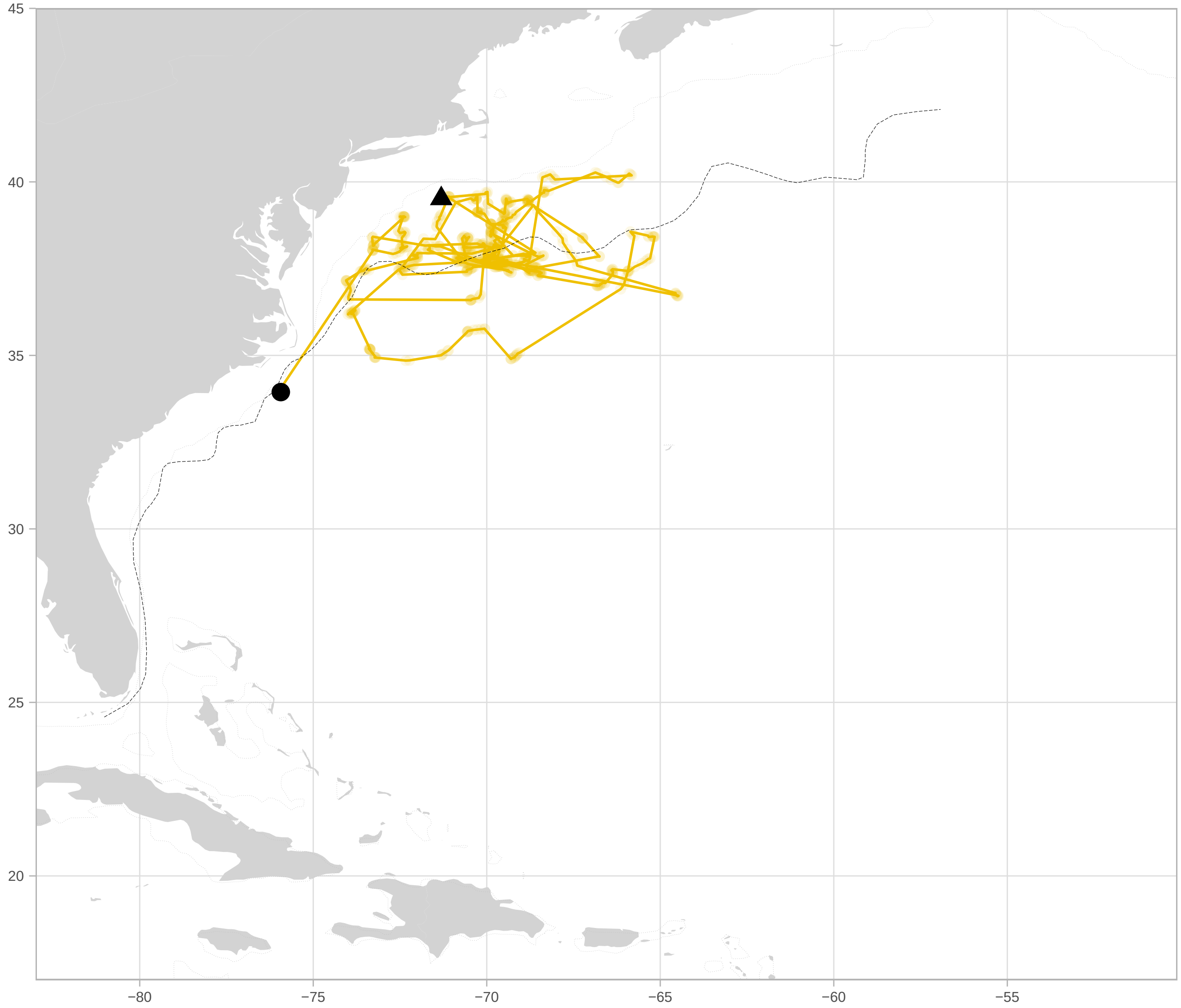

468: 2019−05−09 to 2019−08−31

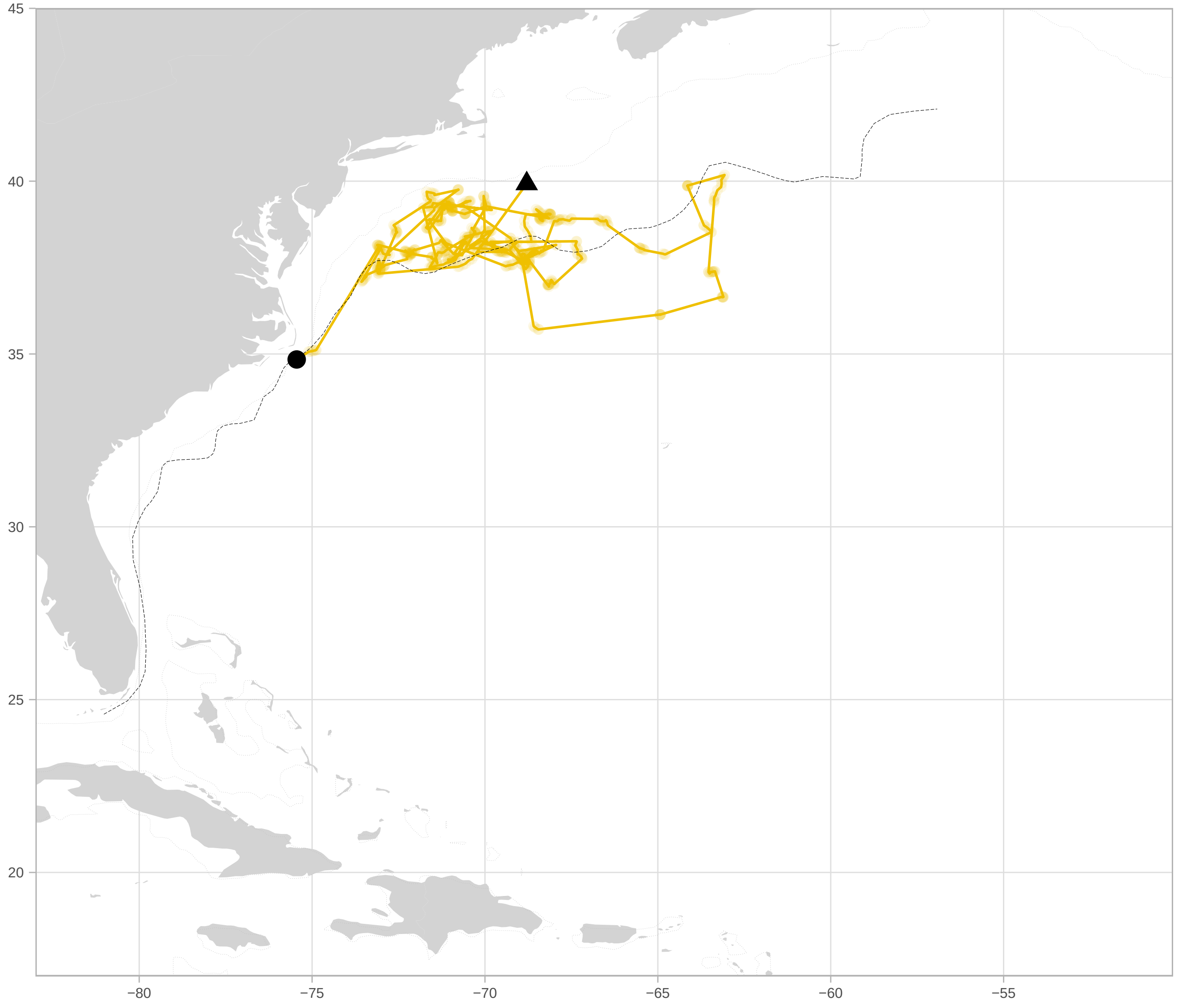

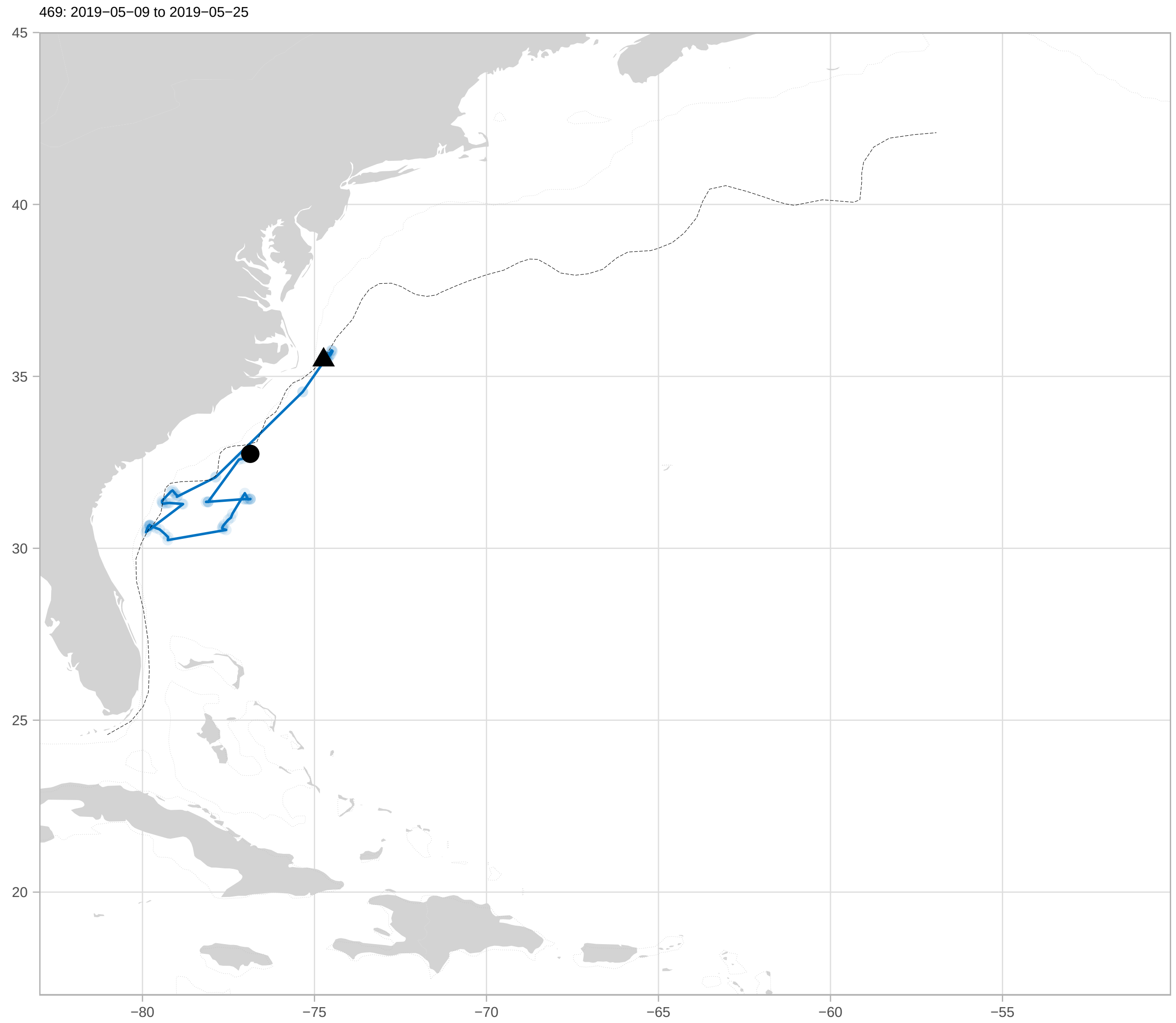

469: 2019−05−09 to 2019−05−25
45
40
35
30
25
20
−80
−75
−70
−65
−60
−55

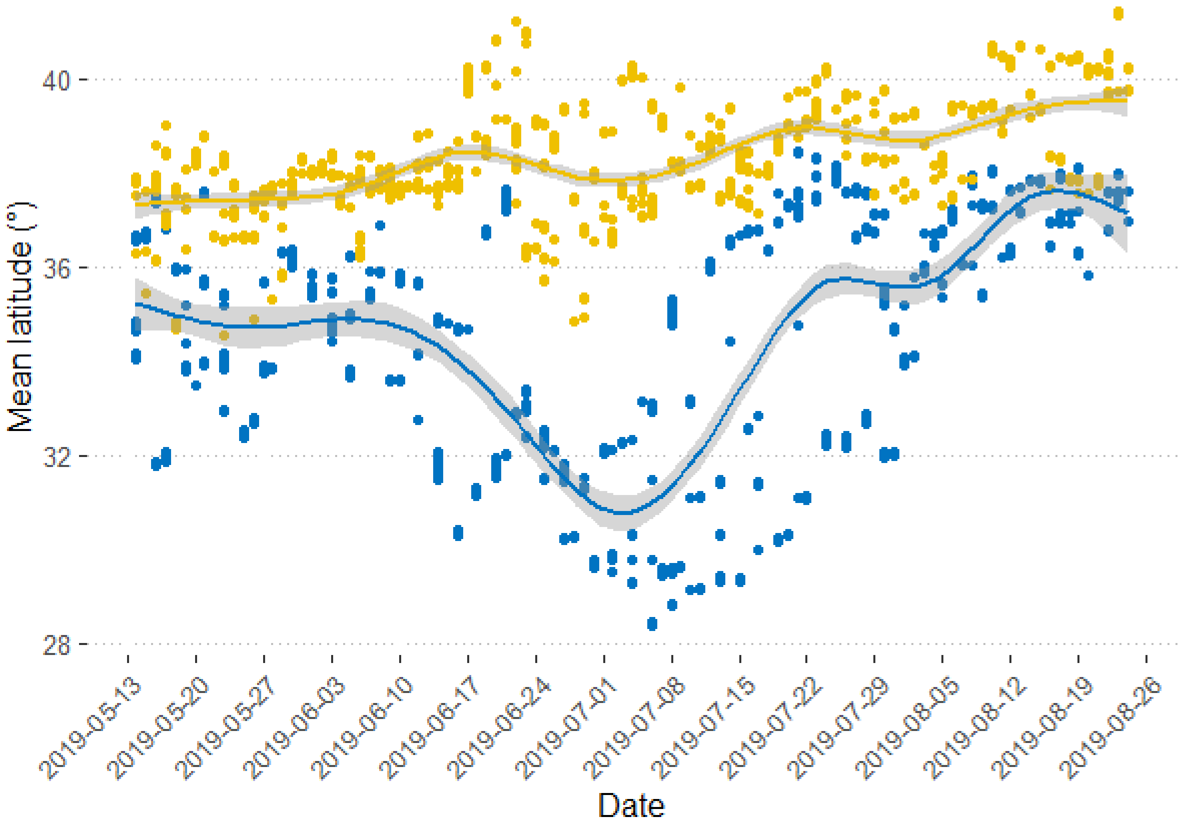


Figure S5. Daily latitude for Black-capped Petrels captured at sea off Cape Hatteras, North Carolina, USA and tracked from May 2019 to August 2019. Blue: dark form; yellow: light form. Shaded area is 95% confidence interval.

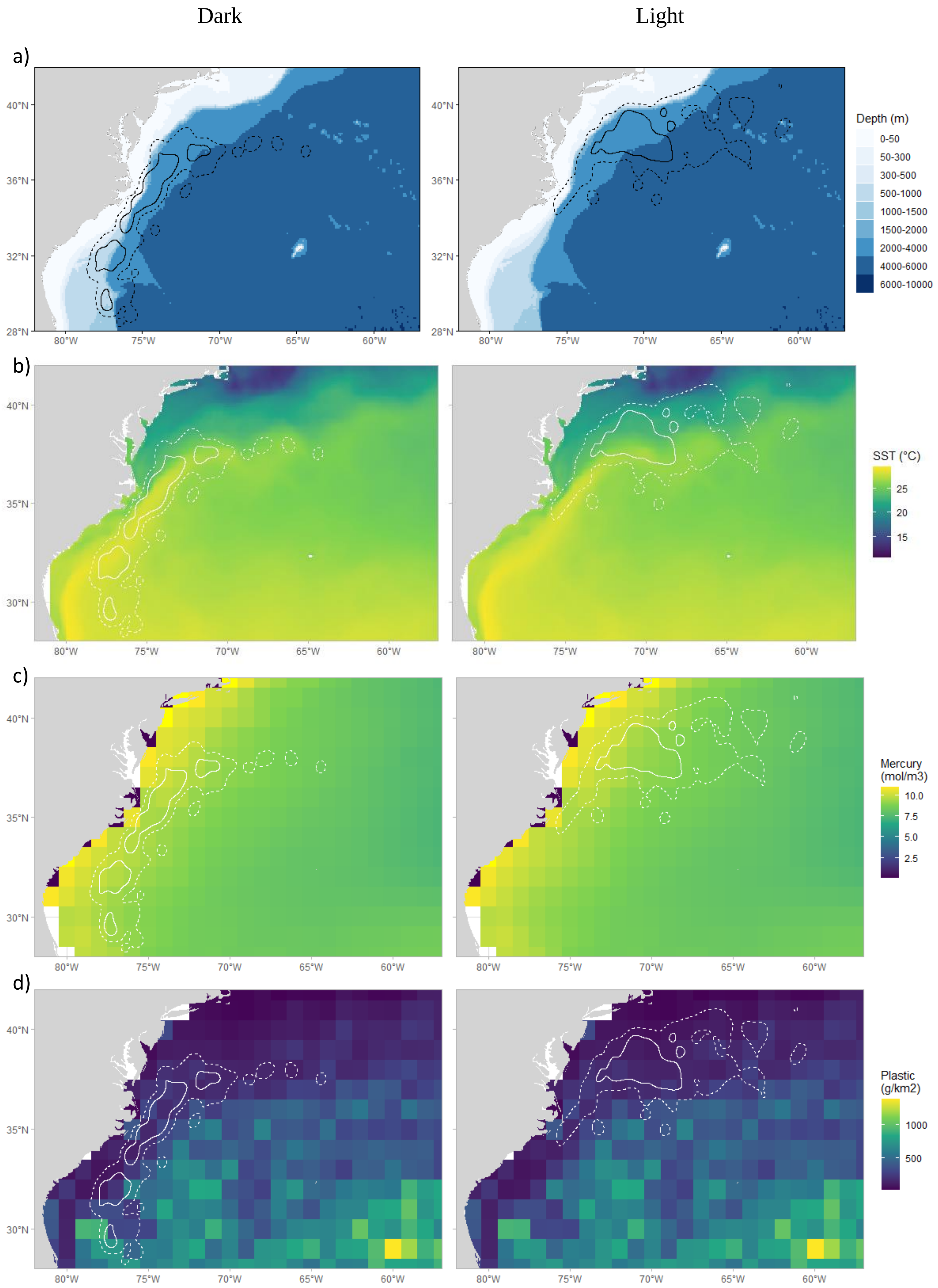
Dark
Light
a)
b)
c)
d)
Depth (m)
0-50
50-300
300-500
500-1000
1000-1500
1500-2000
2000-4000
4000-6000
6000-10000
SST (°C)
25
20
15
Mercury (mol/m3)
10.0
7.5
5.0
2.5
Plastic (g/km2)
1000
500
40°N
36°N
32°N
28°N
35°N
30°N
80°W
75°W
70°W
65°W
60°W

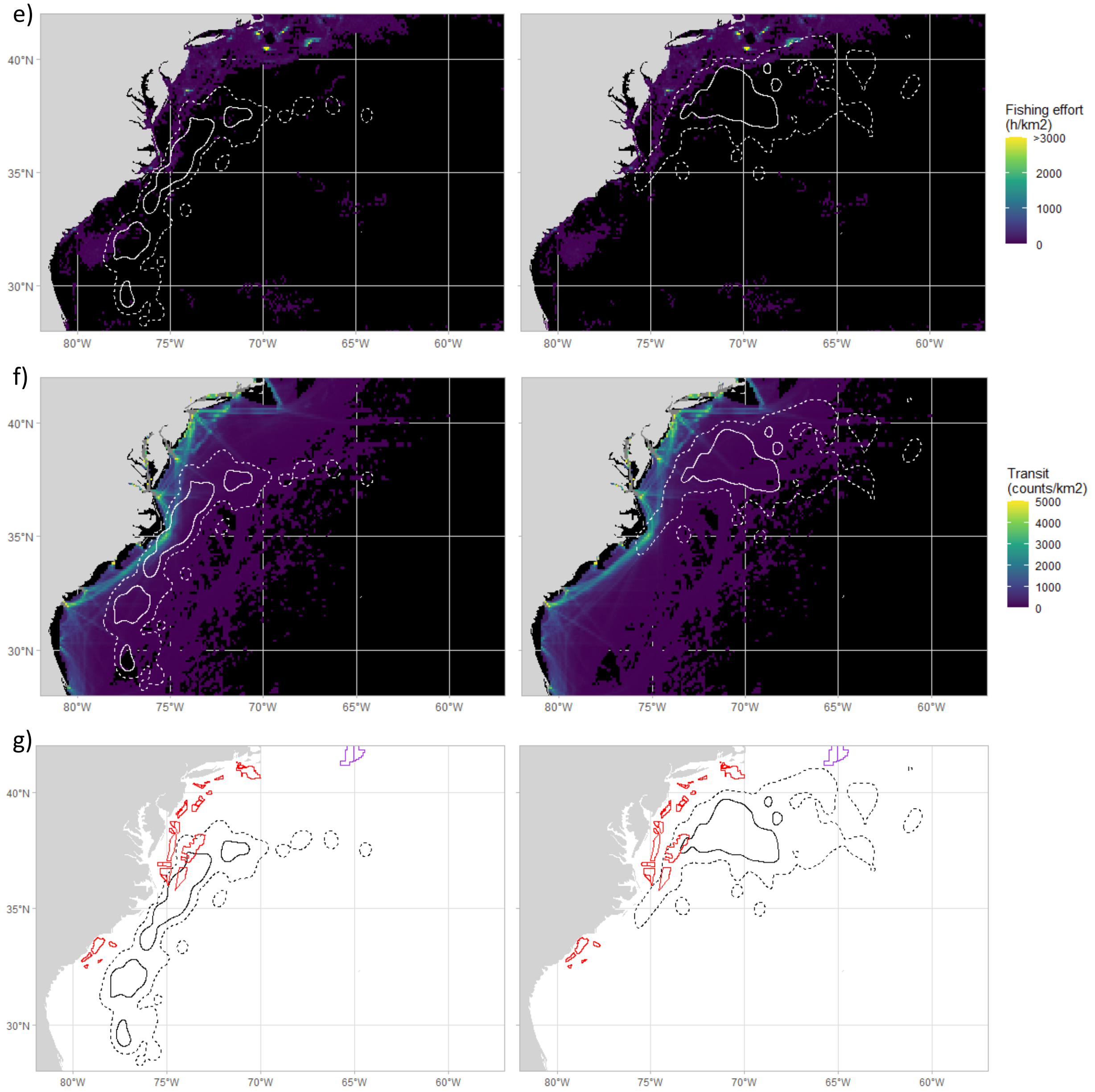


Figure S6. Maps of habitat attributes and marine threats to Black-capped Petrels in the western North Atlantic. a) Bathymetry, b) Sea surface temperature, c) Mercury concentration, d) Microplastics concentration, e) Fishing effort, f) Ship traffic, and g) Marine energy. Left panel: dark forms; right panel: light forms. In all panels, solid black or white lines represent core areas (50% UD) and dashed black or white lines represent home range (90% UD) of petrels tracked from May 2019 – August 2019. In panel G, purple polygons indicate the location of petroleum leases, and red polygons offshore wind leases. UD = Utilization distribution.

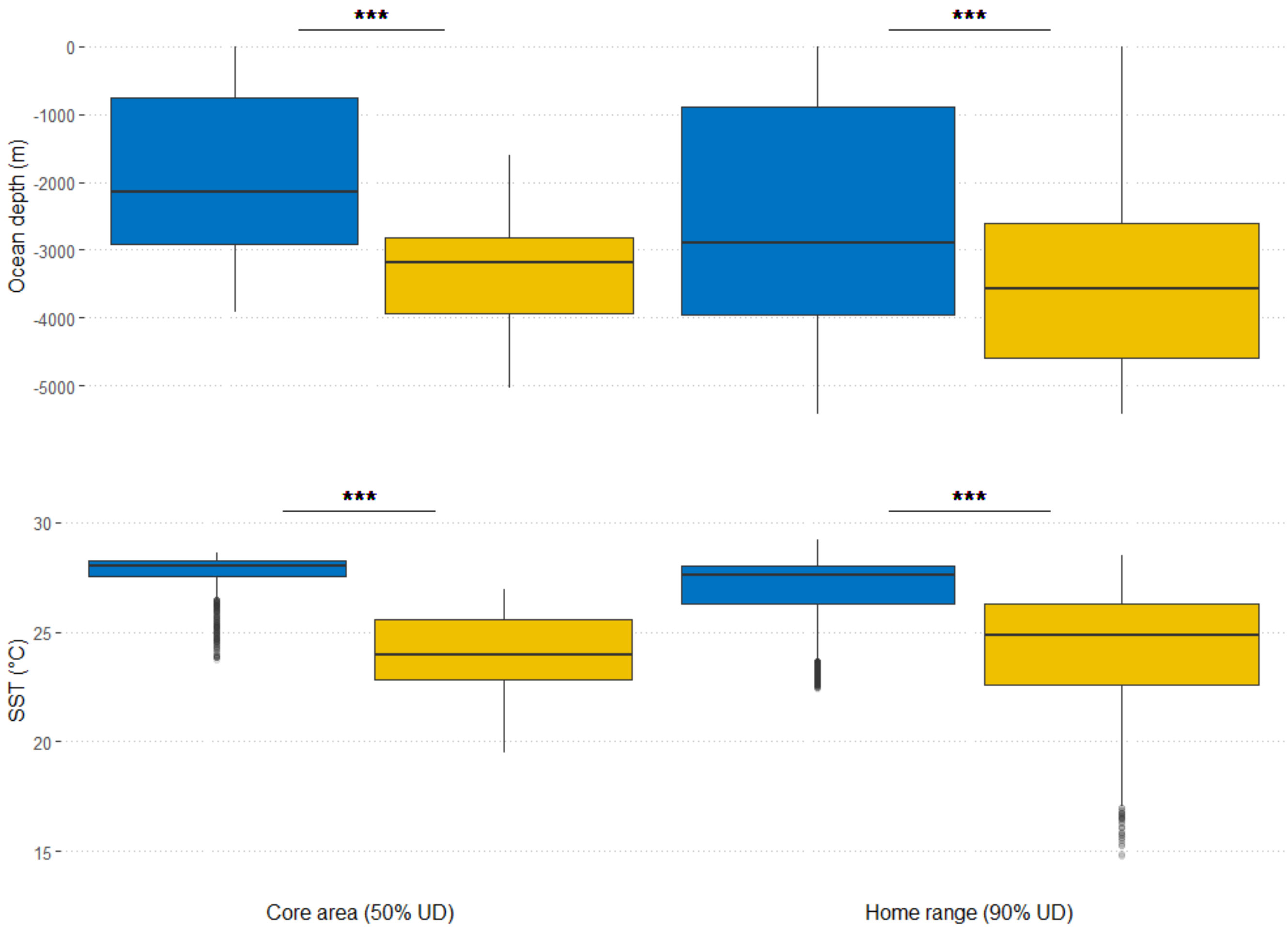


Figure S7. Distribution of values of ocean depth and sea surface temperature in core area and home range of Black-capped Petrels tracked from May 2019 – August 2019. Blue represents dark forms and yellow represents light forms. P-value of Wilcoxon sum rank test: *** <0.005. Boxes depict the median and quartiles, whiskers depict the 5th and 95th percentiles, and circles depict data beyond the 5th and 95th. UD = Utilization distribution.

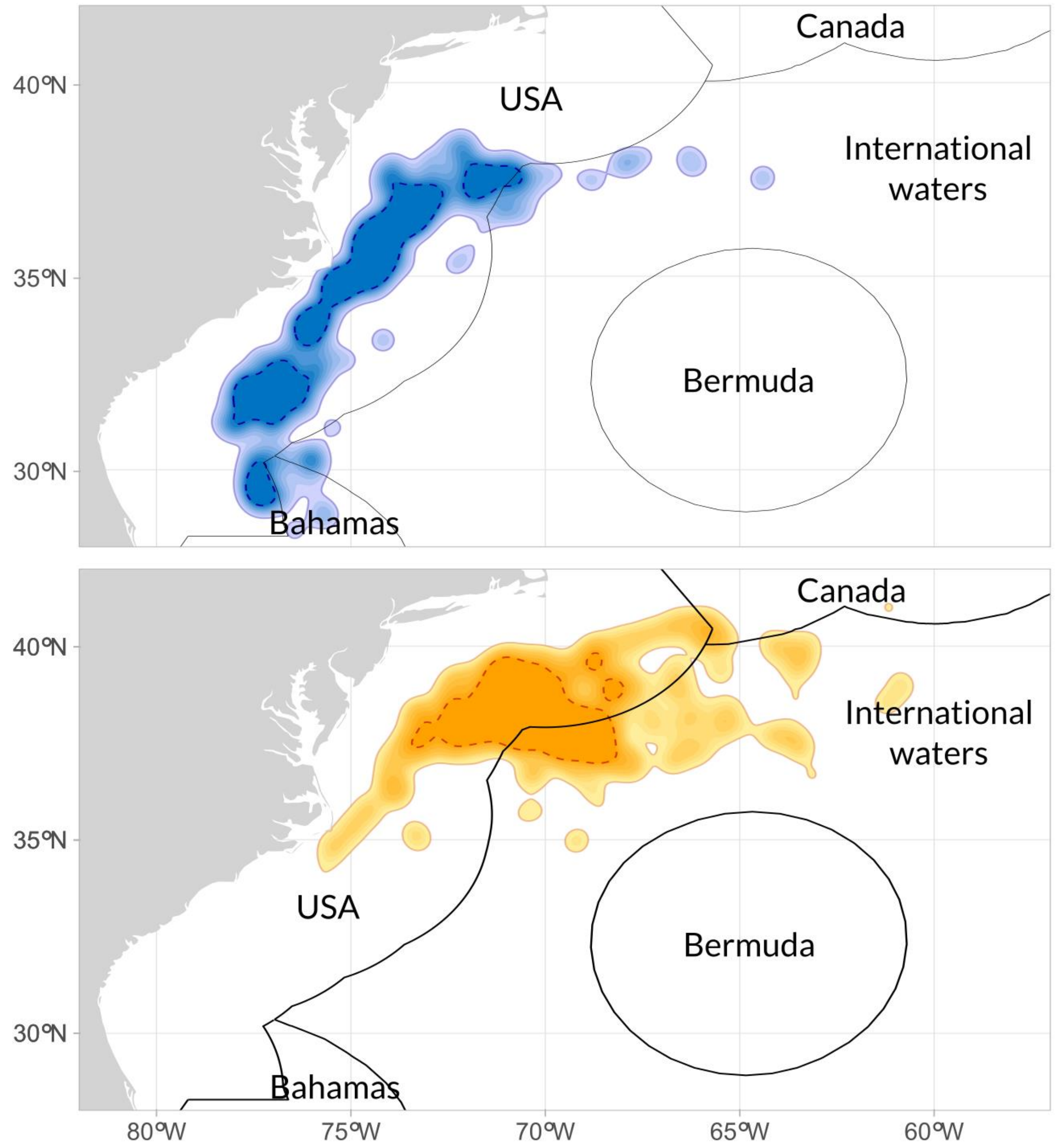


Figure S8. Overlap between exclusive economic zones in the western North Atlantic and the distribution of Black-capped Petrels tracked from May 2019 – August 2019. Top: dark form; bottom: light form. Black lines delineate exclusive economic zones. Colored dashed lines indicate core areas.

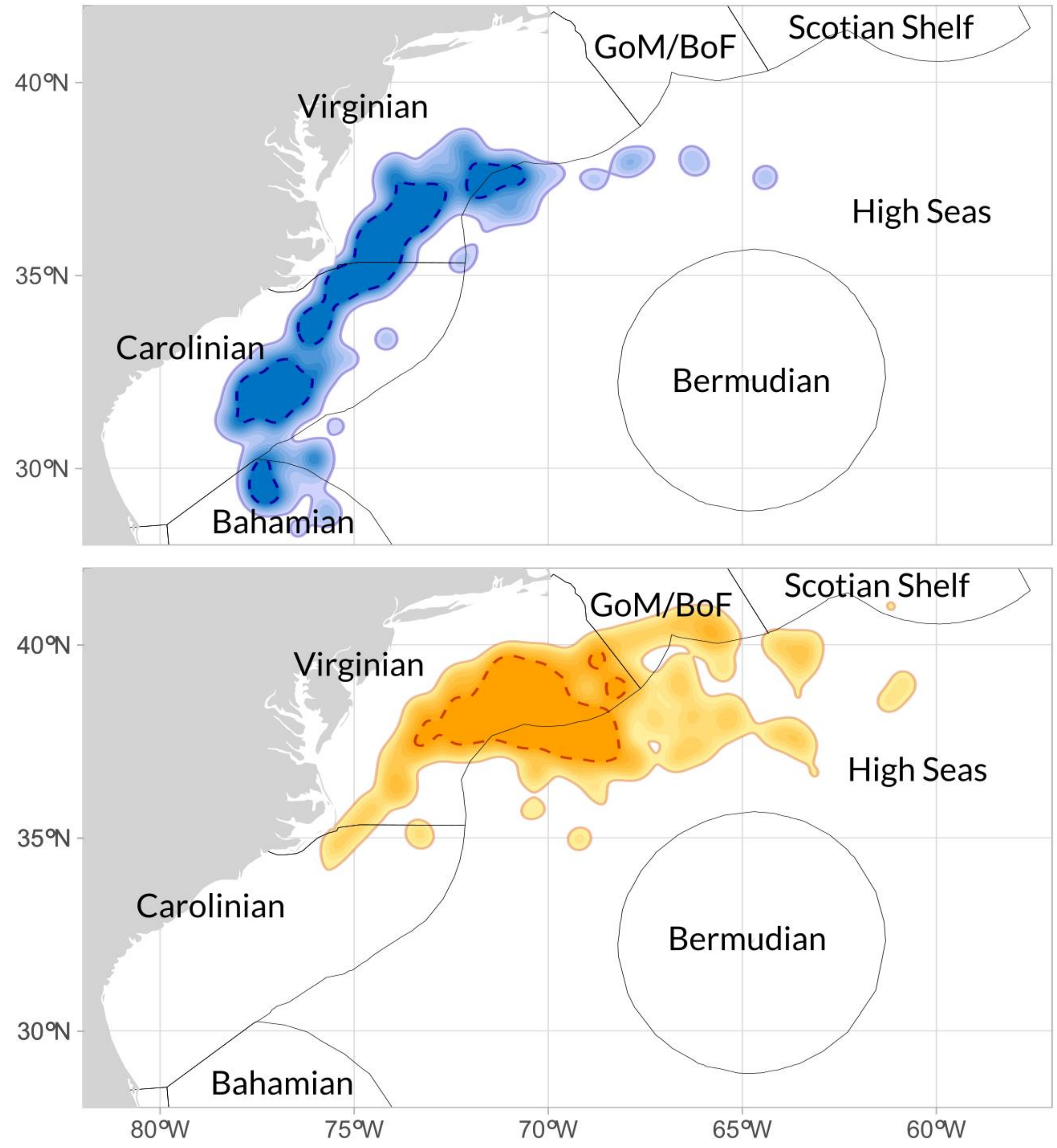


Figure S9. Overlap between marine ecoregions in the western North Atlantic and the distribution of Black-capped Petrels tracked from May 2019 – August 2019. Top: dark form; bottom: light form. Black lines delineate marine ecoregions (Spalding et al. 2007). Colored dashed lines indicate core areas. GoM/BoF is Gulf of Maine/Bay of Fundy.

Reference:

Spalding, M. D., Fox, H. E., Allen, G. R., Davidson, N., Ferdaña, Z. A., Finlayson, M. A. X., ... & Robertson, J. (2007). Marine ecoregions of the world: a bioregionalization of coastal and shelf areas. *BioScience*, *57*(7), 573-583.

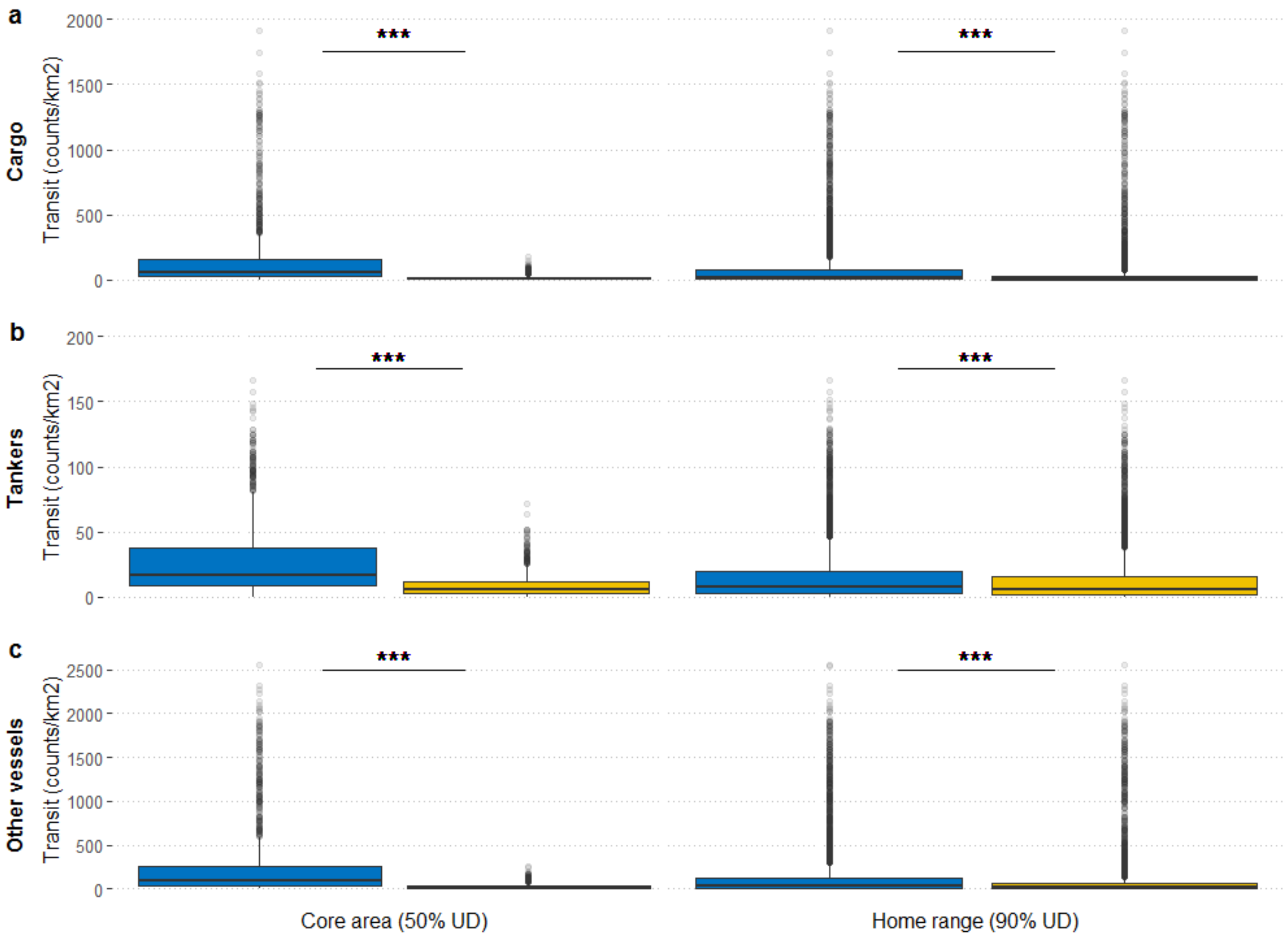


Figure S10. Distribution of values of exposure to vessel transit in core area and home range of Black-capped Petrels tracked from May 2019 – August 2019. Blue represent dark forms and yellow represents light forms. P-value of Wilcoxon sum rank test: *** <0.005. Boxes depict the median and quartiles, whiskers depict the 5th and 95th percentiles, and circles depict data beyond the 5th and 95th. UD = Utilization distribution.

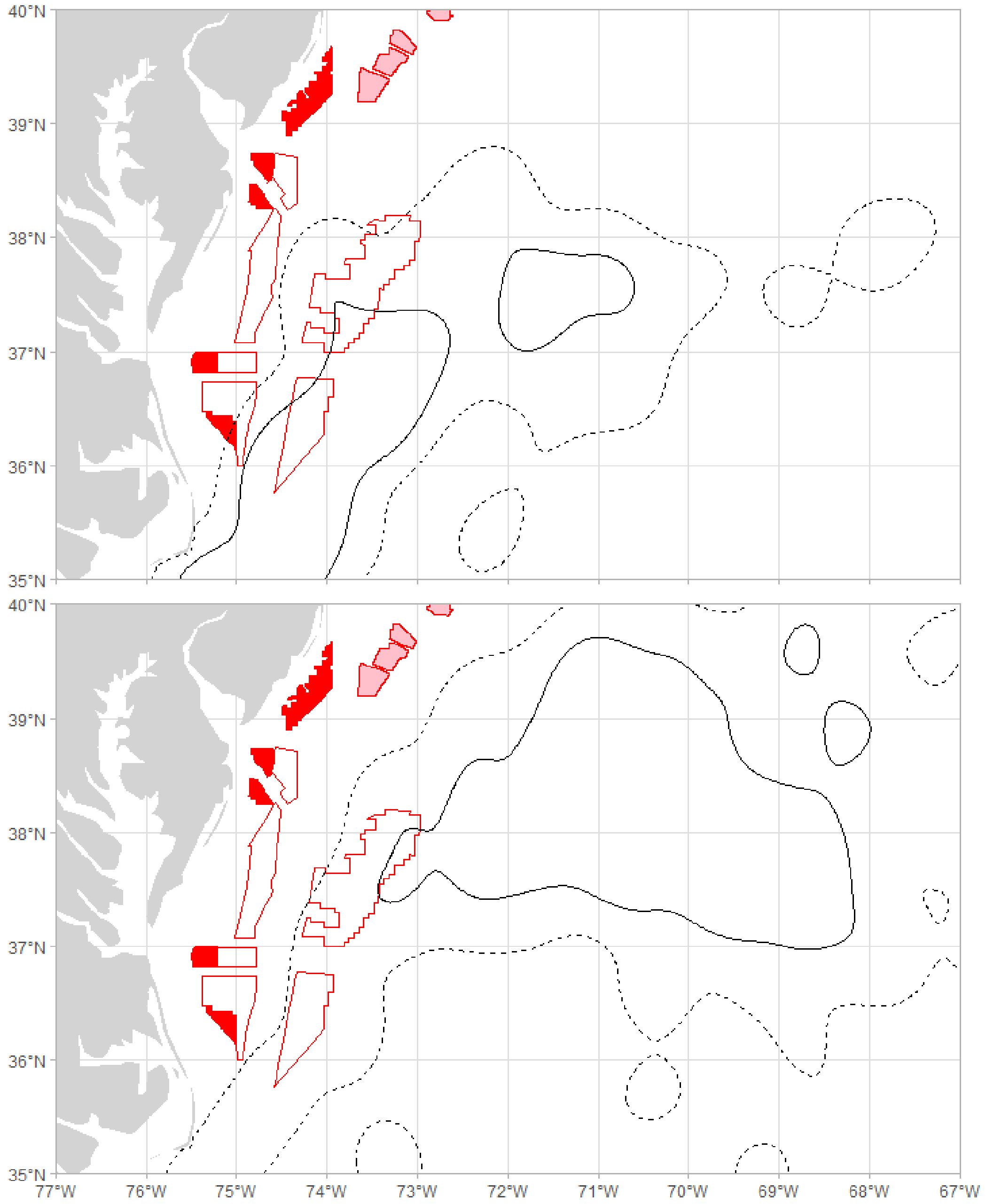


Figure S11. Overlap between lease areas for wind energy production and Black-capped Petrels use areas in the western North Atlantic. Top panel: utilization distributions (UD) of dark forms; bottom panel: utilization distributions of light forms. Solid black lines represent core areas (50% UD) and dashed black lines represent home range (90% UD) of petrels tracked from May 2019 – August 2019. Red-filled polygons indicate the location of active leases, pink-filled polygons indicate planned areas (New York Bight), and outlined polygons indicate draft areas (Central Atlantic).

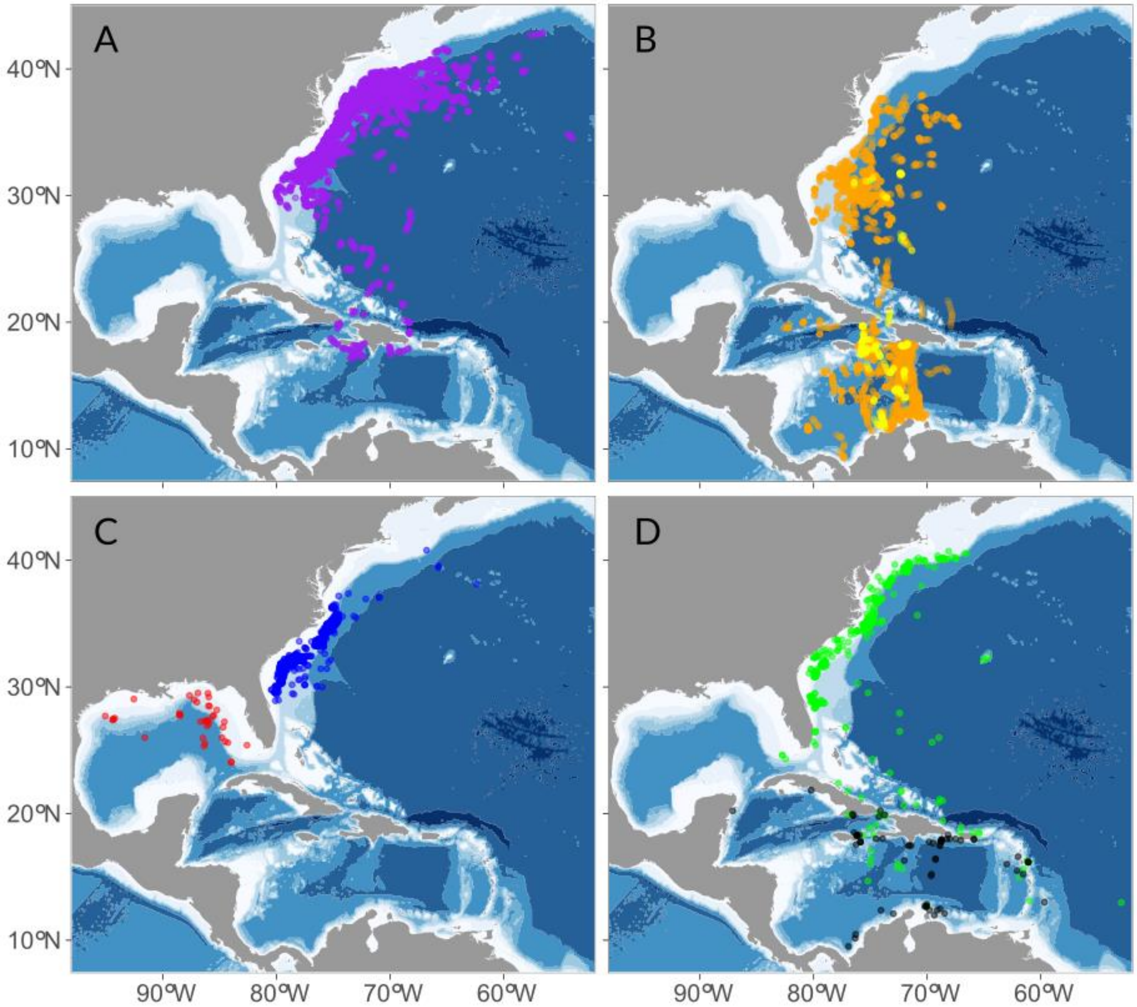


Figure S12. Existing datasets of Black-capped Petrel tracking and observations at sea. A) This study. B) Orange: Jodice et al. (2015); yellow: Satgé et al. (2019). C) Blue: Sussman and U.S. Geological Survey (2014); red: Jodice et al. (2021). D) Black: Leopold et al. (2019), green: eBird (2021).

References:

eBird (2021). eBird: An online database of bird distribution and abundance. eBird, Cornell Lab of Ornithology, Ithaca, New York. Accessed from http://www.ebird.org on 01 November 2021.

Jodice, P. G., Ronconi, R. A., Rupp, E., Wallace, G. E., & Satgé, Y. (2015). First satellite tracks of the endangered black-capped petrel. *Endangered Species Research*, *29*(1), 23-33.

Jodice, P. G., Michael, P. E., Gleason, J. S., Haney, J. C., & Satgé, Y. G. (2021). Revising the marine range of the endangered black-capped petrel *Pterodroma hasitata*: occurrence in the northern Gulf of Mexico and exposure to conservation threats. *Endangered Species Research*, *46*, 49-65.

Leopold, M. F., Geelhoed, S. C., Scheidat, M., Cremer, J., Debrot, A. O., & Van Halewijn, R. (2019). A review of records of the Black-capped Petrel *Pterodroma hasitata* in the Caribbean Sea. *Marine Ornithology*, *47*, 235-241.

Satgé, Y. G., Rupp, E., & Jodice, P. G. (2019). *A preliminary report of ongoing research of the ecology of Black-capped Petrel (*Pterodroma hasitata*) in Sierra de Bahoruco, Dominican Republic–I: GPS tracking of breeding adults*. South Carolina Cooperative Fish and Wildlife Research Unit.

Sussman, Allison & US Geological Survey (2014). *Atlantic Offshore Seabird Dataset Catalog, Atlantic Coast and Outer Continental Shelf, from 1938-01-01 to 2013-12-31 (NCEI Accession 0115356)*. NOAA National Centers for Environmental Information. Dataset. https://accession.nodc.noaa.gov/0115356. Accessed [2020-11-01].
